# Genetic Incorporation of Two Mutually Orthogonal Bioorthogonal Amino Acids That Enable Efficient Protein Dual-Labeling in Cells

**DOI:** 10.1101/2021.04.12.439361

**Authors:** Riley M. Bednar, Subhashis Jana, Sahiti Kuppa, Rachel Franklin, Joseph Beckman, Edwin Antony, Richard B. Cooley, Ryan A. Mehl

**Affiliations:** Department of Biochemistry and Biophysics, Oregon State University, 2011 Agricultural & Life Sciences Building, Corvallis, Oregon 97331-7305, United States; Department of Biochemistry and Molecular Biology, Saint Louis University School of Medicine, Edward A. Doisy Research Center, 1100 South Grand Blvd., St. Louis, MO 63104

## Abstract

The ability to site-specifically modify proteins at multiple sites *in vivo* will enable the study of protein function in its native environment with unprecedented levels of detail. Here, we present a versatile two-step strategy to meet this goal involving site-specific encoding of two distinct noncanonical amino acids bearing bioorthogonal handles into proteins *in vivo* followed by mutually orthogonal labeling. This general approach, that we call <u>d</u>ual <u>e</u>ncoding <u>a</u>nd labeling (DEAL), allowed us to efficiently encoded tetrazine- and azide-bearing amino acids into a protein and demonstrate for the first time that the bioorthogonal labeling reactions with strained alkene and alkyne labels can function simultaneously and intracellularly with high yields when site-specifically encoded in a single protein. Using our DEAL system, we were able to perform topologically-defined protein-protein crosslinking, intramolecular stapling, and site-specific installation of fluorophores all inside living *Escherichia coli* cells, as well as study the DNA-binding properties of yeast Replication Protein A *in vitro*. By enabling the efficient dual modification of proteins *in vivo*, this DEAL approach provides a tool for the characterization and engineering of proteins *in vivo*.

## Introduction

As the fields of molecular and cellular biology develop, the demand for tools that enable defined, independent, and residue-level manipulation of proteins at multiple sites in their native context continues to grow. Recent advances in bioorthogonal chemistry have the potential to overcome this limitation and thereby provide the ability to interrogate protein function in the complex chemical environment of living cells through dual labeling strategies^1^. Many contemporary technologies such as self-labeling enzyme domains, chemoenzymatic labeling, and affinity tags have been used to encode bioorthogonal handles into proteins *in vivo*^2^. However, their bulky size, restrictive placement at the target protein’s termini, and potential to disrupt protein function greatly impinge their utility^3^. In contrast, genetically encoded noncanonical amino acids (ncAAs) bearing bioorthogonal handles can be site-specifically installed at any location in a protein *in vivo* via genetic code expansion (GCE) with minimal perturbations to protein structure and function^4^. Moreover, this GCE approach has been adapted to allow simultaneous incorporation of multiple distinct ncAAs, including bioorthogonally reactive ncAAs, into a single protein *in vivo*^5–7^. Nevertheless, while the GCE approach enables site-specific attachment with small handles, the higher demands it places on efficient and orthogonal labeling reactions that can function inside cells have not been met.

An important requisite to dual modifying protein *in vivo* is that the two labeling reactions must be mutually orthogonal to both one another and to other functional groups present in the cell (labeling orthogonality)^8^. The reactions should also be free of catalysts, high yielding, and rapid under physiological conditions, thereby enabling effective labeling at low protein concentrations and on biologically relevant time frames^9^. Few reactions meet these criteria better than the strain-promoted azide-alkyne coupling (SPAAC) and inverse electron demand Diels-Alder (IEDDA) reactions. The SPAAC reaction is a [3+2] cycloaddition reaction originally described by Bertozzi and colleagues that commonly occurs between azides and dibenzoannulated cyclooctynes (DBCO)^10^. Despite modest reaction kinetics (∼0.1 - 1 M^-1^ s^-1^), its biocompatibility has made it one of the most extensively used bioorthogonal reactions for *in vivo* labeling^8,10^. The IEDDA reaction is a [4+2] cycloaddition that occurs between an electron-deficient diene such as a 1,2,4,5-tetrazine and an electron-rich dienophile such as a strained alkene^1^. Bioorthogonal IEDDA reactions, such as those first described by Fox and colleagues, can reach rates upwards of 10^6^ M^-1^ s^-1^, allowing complete reaction within minutes at sub-micromolar concentrations^1,9,11–14^. Amino acid derivatives containing azide, cyclopropene, alkyne, *trans*-cyclooctene (TCO), and tetrazine functionalities have all been genetically encoded into proteins^15^, albeit not yet in a manner conducive to dual encoding and subsequent intracellular labeling *in vivo*.

While density functional theory predicts that SPAAC and IEDDA reactions should be mutually orthogonal and the *in vivo* compatibility of these reactions has been shown, site-specific encoding of the handles into the same protein and dual labeling *in vivo* has not yet been demonstrated^16–18^. A recent attempt succeeded at dual encoding two different handles into a protein using GCE, however, the functional groups required the use of copper catalyst, thereby relegating labeling to the cell surface due to copper toxicity^19^. These examples highlight the challenge of developing efficient and orthogonal dual encoding and labeling approaches. An effective solution to this challenge should be encoding azide and tetrazine moieties, since they exhibit high biostability, participate in biocompatible, mutually orthogonal reactions, and are not anticipated to cross-react during the long incubation periods required for GCE^9,16,20^. Provided tetrazine- and azide-bearing amino acids can be encoded efficiently into the same protein, and the mutual orthogonality of the SPAAC and IEDDA reactions persists *in vivo*, this dual protein labeling approach would open avenues to manipulate and study molecular processes with greater scrutiny than is currently accessible.

To test this hypothesis, we developed the first genetic code expansion system that enables the simultaneous and site-specific incorporation of the two bioorthogonally reactive ncAAs, *para*-azidophenylalanine (pAzF) and a tetrazine-containing ncAA (Tet3.0), into proteins *in vivo*. Our optimized GCE system enables robust production of dual-ncAA containing proteins and allowed us to characterize the dual labeling efficiency and orthogonality on various proteins *in vivo* and *in vitro*. We demonstrate the utility and versatility of our <u>d</u>ual <u>e</u>ncoding <u>a</u>nd labeling (DEAL) system by showcasing a diverse array of *in vivo* abilities, such as site-specific installation of FRET pairs, topologically-defined protein-protein crosslinking, and site-specific intramolecular protein stapling (Fig. 1), as well as applying our system to the study of the challenging *Saccharomyces cerevisiae* Replication Protein A (*Sc*RPA) complex. These examples illustrate the precision and power of our optimized DEAL system to meet the demands of rapidly advancing biological fields.

**Figure 1.**
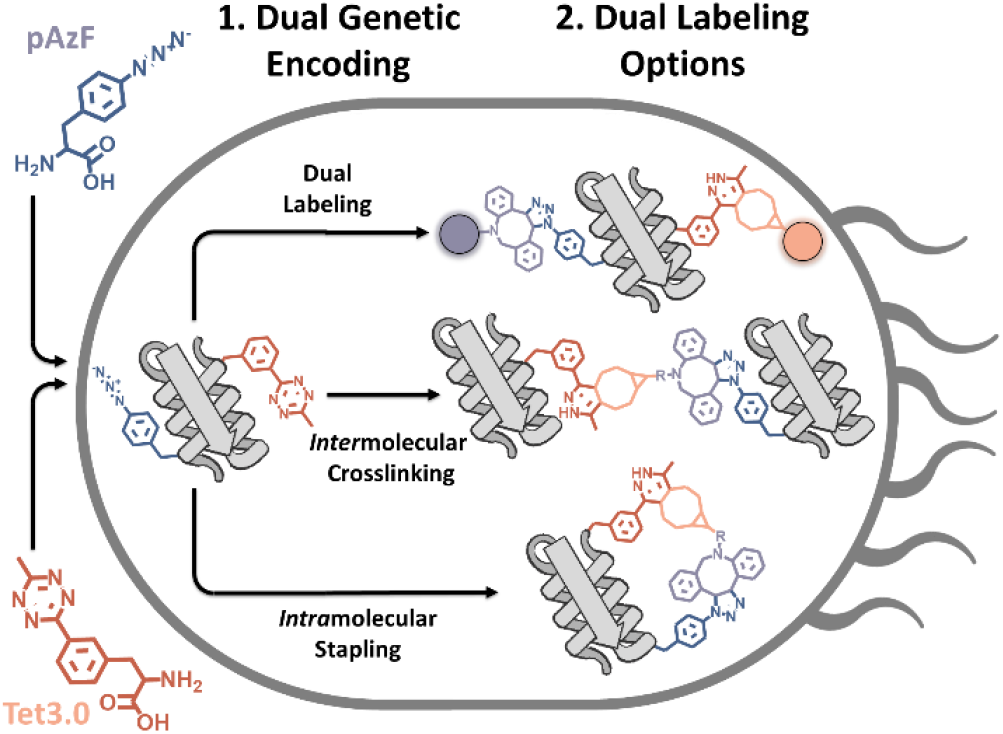
Schematic of *in vivo* dual encoding and labeling. Noncanonical amino acids (ncAAs) *para*-azidophenylalanine (pAzF; blue) and a meta-substituted tetrazine-containing phenylalanine derivative (Tet3.0; orange) are encoded into proteins using genetic code expansion via mutually orthogonal, dual nonsense suppression systems. Controlled bioorthogonal labeling at both sites within proteins enables site-specific dual labeling, topologically-defined intermolecular crosslinking and intramolecular stapling *in vivo*.

## Results and Discussion

### Organization of Genetic Code Expansion Components

Generating a dual-ncAA suppression system capable of supporting the simultaneous encoding of two distinct ncAAs requires two mutually orthogonal suppression systems (subsystems), each comprised of four components: 1) the ncAA to be encoded, 2) the aminoacyl tRNA synthetase (aaRS) engineered to load the ncAA onto 3) the cognate tRNA and 4) codon that will be suppressed by the ncAA-tRNA during translation of the protein of interest. As such, we elected to use a popular *Methanocaldococcus jannaschii* tyrosyl aminoacyl tRNA synthetase/tRNA pair (*Mj*TyrRS/tRNA) that was originally generated for the incorporation of *para*-cyanophenylalanine, but that also shows polyspecificity towards pAzF^21^, to encode this ncAA at amber stop codons (herein referred to as pAzFRS/tRNA_CUA_). To drive incorporation at ochre stop codons, we turned to an engineered *Methanosarcina barkeri* Pyrrolysyl aminoacyl tRNA synthetase (*Mb*PylRS/tRNA_UUA_ that efficiently recognizes a meta-substituted 1,2,4,5-tetrazine-containing phenylalanine derivative (Tet3.0RS/tRNA_UUA_) previously selected by our laboratory^13^. The choice of anticodons was based on two factors. First, the tRNA anticodon loop is a known identity factor for *Mj*TyrRS and, as such has historically been utilized as a UAG suppression system with known success^22^, while the tRNA anticodon is not an identity factor for *Mb*PylRS and can therefore tolerate codon reassignment of its cognate tRNA^23^. Second, The UGA codon was not considered because it is known to experience near-cognate suppression from endogenous 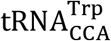^24^.

We performed optimization experiments to identify the most efficient combination of cell strains and vectors for hosting the two suppression subsystems and reporter gene under single stop codon suppression conditions (Fig. S2). These experiments revealed that the pAzFRS/tRNA_CUA_ and Tet3.0RS/tRNA_UUA_ are most efficient when in the pEVOL^25^ and pUltraI^26^ plasmids, respectively, and that these both function most optimally when the reporter gene is expressed from a pET28 vector in BL21(DE3) cells (for a detailed discussion of these results, see the supplemental section “Evaluation of UAG and UAA Single Site Suppression”). As such, our finalized dual suppression system is a modular, three-vector configuration, with each suppression system placed on a separate plasmid, and the gene of interest contained on a third plasmid (Figure S1).

### Dual incorporation of pAzF and Tet3.0

Earlier reports on dual ncAA incorporation have identified points in translation where orthogonality may become compromised, such as the substrate specificity of the aaRS, the interactions between aaRS and tRNA, and the specificity of tRNAs during decoding^26,27^ (Fig. S1). The orthogonality of the individual aaRS/tRNA pairs in our systems, *Mj*TyrRS/tRNA_CUA_ and *Mb*PylRS/tRNA_UUA_, has previously been established^26^. Therefore, we characterized the orthogonality of our dual-encoding subsystems at the ncAA-aaRS and tRNA-codon levels. We observed that, in the absence of ncAA for the pAzFRS/tRNA_CUA_ subsystem, UAG codons can be suppressed by near-cognate suppression by endogenous decoding systems, and that the UAA suppressor tRNA for the Tet3.0RS/tRNA_UUA_ subsystem can decode UAG stop codons through wobble-pairing (Fig. S3)—both of which have previously been observed^26,28^. Importantly, these breaches in encoding orthogonality are situational, and do not manifest when both subsystems are present and functioning properly (see section “Evaluation of GCE Orthogonality” in the supplemental section for a full discussion).

Having established that our optimized pAzFRS/tRNA_CUA_ and Tet3.0RS/tRNA_UUA_ subsystems are contextually and mutually orthogonal, we sought to combine them for the dual encoding of pAzF and Tet3.0 into proteins. To do so, we paired a <u>S</u>mall <u>U</u>biquitin-like <u>Mo</u>difier-<u>s</u>uperfolder <u>G</u>reen <u>F</u>luorescent <u>P</u>rotein (SUMO-sfGFP) fluorescent reporter possessing UAG and UAA codons at sites 35 and 102, respectively, with both subsystems and observed a 10% yield as compared to SUMO-sfGFP^WT^ when both ncAAs were added (Fig. 2A). We also reversed the order of the nonsense codons by introducing UAA and UAG codons at positions 35 and 102 and observed a 22% yield (Fig. 2A), indicating that the order of the stop codons can be reversed and that this orientation may improve protein yields. Unsurprisingly, we also observed above-background fluorescence when pAzF and/or the pAzFRS/tRNA_CUA_ subsystem were absent (Fig. 2A), which we attribute to the situational breaches in encoding orthogonality previously mentioned (Fig. S3). To confirm the fidelity of ncAA incorporation, sfGFP possessing both pAzF and Tet3.0 at positions 134 and 150 (sfGFP^Dual^) was expressed and purified yielding 26 mg protein per L of expression media, approximately 12% of sfGFP^WT^ (Table S5). Electrospray ionization (ESI) mass spectrometry analysis on sfGFP^Dual^, sfGFP^pAzF^, sfGFP^Tet3.0^, and sfGFP^WT^ expressed and purified under analogous conditions showed masses consistent with the expected ncAA-substitutions (Fig. 2B). These results verify that our dual-encoding system can be used to achieve efficient, simultaneous, and homogenous site-specific incorporation of pAzF and Tet3.0 into the same protein.

**Figure 2.**
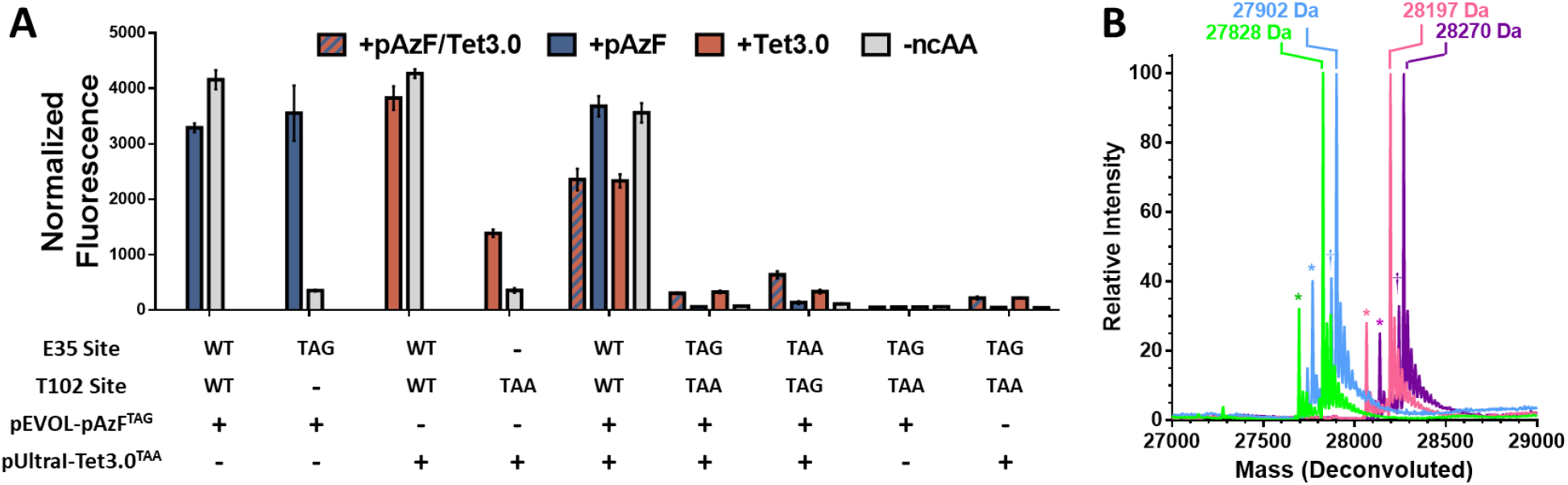
Validation of dual suppression. (A) Normalized SUMO-sfGFP fluorescence for single and dual UAG-UAA suppression at 24 hours in the presence of both pAzF and Tet3.0 (hashed blue and orange), pAzF alone (blue), Tet3.0 alone (orange), and in the absence of ncAAs (gray). (B) Overlaid ESI mass spectra of sfGFP containing either pAzF (blue) or Tet3.0 (pink) at site 150, or together at sites 134 (pAzF) and 150 (Tet3.0) (purple), or neither (WT; green). Observed masses are indicated above each major peak. Peaks corresponding to the loss of N-terminal methionine are indicated by “*”, while peaks corresponding to pAzF reduction are indicated by “†”. Mass measurement error is ± 1 Da, see Table S6 for expected masses.

### Evaluation of dual labeling *in vitro*

Dual encoding of pAzF and Tet3.0 into a protein should enable efficient one-pot dual labeling. To test this, we produced a series of SUMO-sfGFP fusion proteins with either pAzF or Tet3.0 individually in each domain at either site 35 or 253 (SUMO^pAzF^-sfGFP or SUMO-sfGFP^Tet3.0^) or together (SUMO^pAzF^-sfGFP^Tet3.0^) in yields ranging from ∼100 to ∼11 mg/L (Table S5). Upon exposure to a DBCO-TAMRA fluorophore, only proteins possessing pAzF exhibited labeling, as determined by SDS-PAGE in-gel fluorescence (Fig. 3B). Likewise, exposure to a strained *trans*-cyclooctene-terminated 5 kDa polyethylene glycol polymer (sTCO-PEG_5000_) led to a mobility shift only for SUMO-sfGFP^Tet3.0^ (Fig. 3B). When SUMO^pAzF^-sfGFP^Tet3.0^ was exposed to both DBCO-TAMRA and sTCO-PEG_5000_ we observed the formation of a fluorescent, mobility-shifted product (Fig. 3B). To verify labeling orthogonality, we added ULP1 protease to cleave the labeled SUMO domain, and as expected, observed a fluorescently-labeled SUMO fragment and a mobility-shifted sfGFP fragment (Fig. S4).

**Figure 3.**
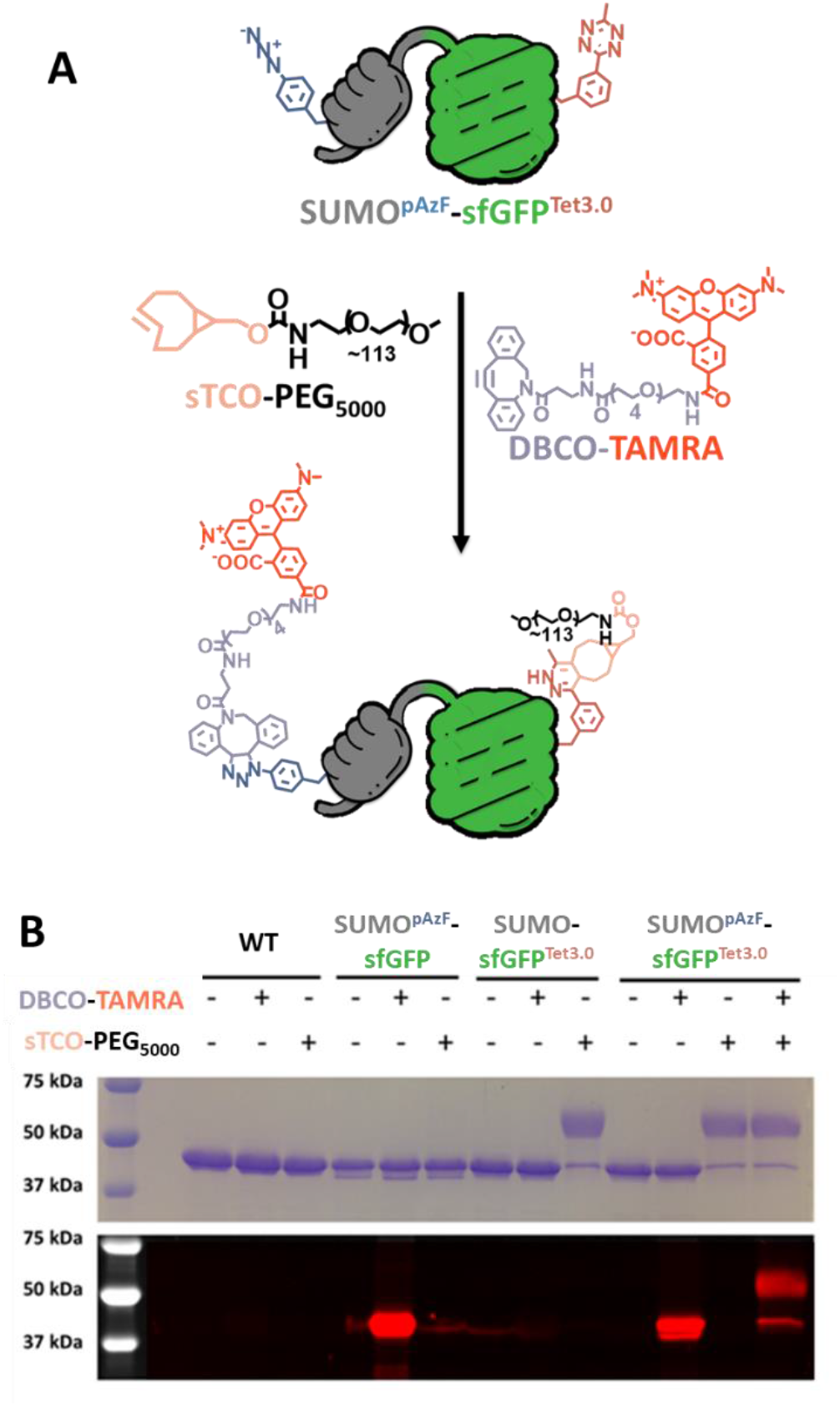
*In vitro* dual labeling of dual encoded SUMO-sfGFP. (A) Reaction scheme of SUMO^pAzF^-sfGFP^Tet3.0^ dual labeling. (B) SDS-PAGE of SUMO^pAzF^-sfGFP^Tet3.0^ exposed to DBCO-TAMRA and/or sTCO-PEG_5000_ imaged by Coomassie staining (top) and in-gel fluorescence (bottom). In these experiments, pAzF is incorporated at position 35 (in the SUMO domain), and Tet3.0 at position 253 (in the sfGFP domain). The contents of each lane are indicated above.

To quantify the labeling extent of SPAAC and IEDDA we utilized a modified blocking assay developed by Murrey and colleagues^29^ with a series of sfGFP constructs that positions pAzF or Tet3.0 at site 150 (sfGFP^pAzF^ and sfGFP^Tet3.0^) or together at sites 134 and 150, respectively (sfGFP^Dual^). This assay consists of an initial non-visualizable labeling step with DBCO-NH_2_ and/or sTCO-OH, followed by a quenching step with excessive DBCO-TAMRA and/or sTCO-JF669 dyes. Quantifying the loss in fluorescence during step 2 as a result of the reaction in step 1, we determined that SPAAC and IEDDA proceeded efficiently for these constructs *in vitro*, with yields of roughly 40-80% and >90%, respectively (Fig. 4A-B). It should be noted that this assay only allows us to quantify relative reaction extent, since compromised handles are unreactive in both labeling steps of the assay. In addition, we observed that 24-hour SPAAC labeling between pAzF and DBCO-NH_2_ followed by 15-minute IEDDA labeling between Tet3.0 and sTCO-OH maximized *in vitro* labeling (>80%) while minimizing cross-reactivity (<4%) (Fig. S5A-B), consistent with observations made by Karver et al., who noted that azides exhibit slight reactivity towards TCO^16^.

**Figure 4.**
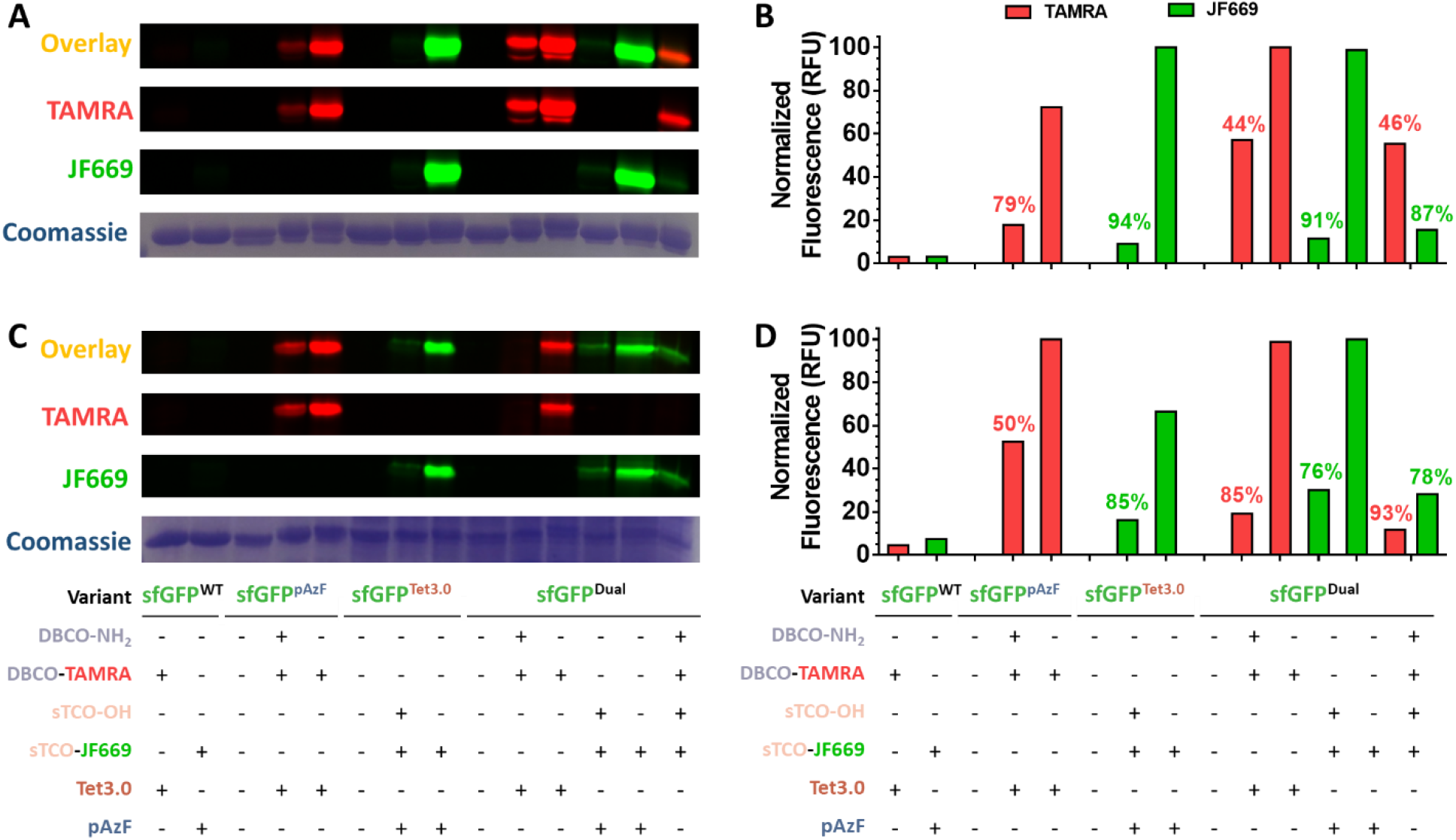
Quantification of dual labeling reactions *in vitro* and *in vivo* using two step blocking labeling (see methods). (A) SDS-PAGE of *in vitro* reactions on sfGFP variants labeled with sTCO-OH and/or DBCO-NH_2_ followed by sTCO-JF669 and/or DBCO-TAMRA prior to in-gel fluorescence imaging; top panel is the overlay of TAMRA and JF669 signals followed by each individual channel, and Coomassie staining. Encoding positions for sfGFP^Tet3.0^ and sfGFP^pAzF^ are at site 150, while encoding of sfGFP^Dual^ includes Tet3.0 at site 134, and pAzF at site 150. (B) Densitometry quantification of the fluorescent gels in panel A (red bars are quantifications of the TAMRA channel, and green bars are quantifications of the JF669 channel). The DBCO-NH_2_ and/or sTCO-OH labeling yields are listed above the corresponding bar. (C) *In vivo* labeling analysis analogously presented to panel A. (D) Densitometry quantification of fluorescent gel channels in panel C, processed and presented analogously to panel B.

Using these labeling conditions, we reacted the aforementioned sfGFP constructs with DBCO-NH_2_ and sTCO-OH *in vitro* and analyzed the products by mass spectrometry. Peaks corresponding to dual-labeled sfGFP were observed as the dominant product (Fig. S6), consistent with the observed orthogonality (Figs. 3 and S5). Minor peaks are consistent with pAzF reduction to *para*-aminophenylalanine, a common modification produced during protein expression^30^ or salt adducts (see supplemental section “Mass Spectrometry Analysis”). Together these results indicate that we have identified optimal labeling conditions that enable the SPAAC and IEDDA reactions to proceed efficiently and orthogonally on proteins containing pAzF and Tet3.0.

### Evaluation of dual labeling *in vivo*

Despite numerous examples of dual encoding of ncAAs into proteins *in vivo*^5–7,19^, to our knowledge there are no reports of site-specific dual labeling of proteins inside a cell. To demonstrate *in vivo* DEAL with our system, we utilized our modified blocking assay to quantify labeling of our previously used sfGFP constructs in cells. Similar to *in vitro* labeling, *in vivo* labeling proceeded efficiently, approaching 50-93% and 76-85% for SPAAC and IEDDA reactions, respectively (Fig. 4C-D), confirming that the cellular environment does not pose a major impetus to dual-labeling with these ncAAs. To evaluate labeling orthogonality of the SPAAC and IEDDA reactions *in vivo* we used our *in vitro* cross-reactivity analysis method by exposing cells containing sfGFP^pAzF^ and sfGFP^Tet3.0^ to sTCO-OH and DBCO-NH_2_, respectively (Fig. S5C-D). Again, we detected minimal cross-reactivity not exceeding 6%, indicating that under these conditions the SPAAC and IEDDA reactions retain their orthogonality.

### Dual Encoding and Labeling *in vivo*

Having confirmed our new DEAL system functions efficiently *in vivo*, we next sought to explore the utility of our system by showcasing three *in vivo* capabilities: (1) intermolecular protein-protein crosslinking, (2) intramolecular protein stapling, (3) and dual-fluorophore labeling (Fig. 1). To aid in our analysis we developed a second fluorescent reporter protein, the blue fluorescent protein mTagBFP2, which has excellent spectral overlap with sfGFP^31^. We screened 6 sites for their ability to tolerate UAG and UAA suppression and selected site 105, with a yield of 79 mg/L when pAzF is encoded (Table S5, Fig. S7; see supplementary discussion “*mTagBFP2 Screening and Characterization*”). All constructs including mTagBFP2 encode pAzF at this relative position, while all constructs including sfGFP encode Tet3.0 at relative position 150.

#### Intermolecular Protein-Protein Crosslinking in vivo

Our DEAL system also offers a plethora of potential *in vivo* applications if dual encoding is applied to two distinct proteins, such as programmable supramolecular protein-complex formation. Here we demonstrate this through covalent intermolecular hetero-protein protein crosslinking between sfGFP^Tet3.0^ and mTagBFP2^pAzF^ in *E. coli*. To do so, we generated a pETduet vector that most balances the expression of these constituent proteins under dual encoding conditions, and confirmed orthogonal labeling of the ncAAs therein (Fig. S8) (see the supplemental section “*sfGFP/mTagBFP2 and sfGFP-mTagBFP2 Expression and Characterization*” for a full discussion).

We first verified that these two ncAA-containing proteins could be successfully crosslinked *in vitro*, by incubating purified sfGFP^Tet3.0^ and mTagBFP2^pAzF^ with a heterobifunctional sTCO-DBCO crosslinker followed by SDS-PAGE and analytical size-exclusion chromatography (Fig. 5A-C). A distinct higher molecular weight product was produced with a retention volume of 14 mL (compared to ∼16 mL for the unlinked proteins) that absorbed at both 399 and 485 nm, the absorption maxima of mTagBFP2 and sfGFP, respectively (Fig. 5B-C). We isolated this product and determined that it possesses spectral properties consistent with sfGFP and mTagBFP2, and displays detectable FRET between the two, supporting the formation of a crosslink between the two proteins (Fig. S9).

**Figure 5.**
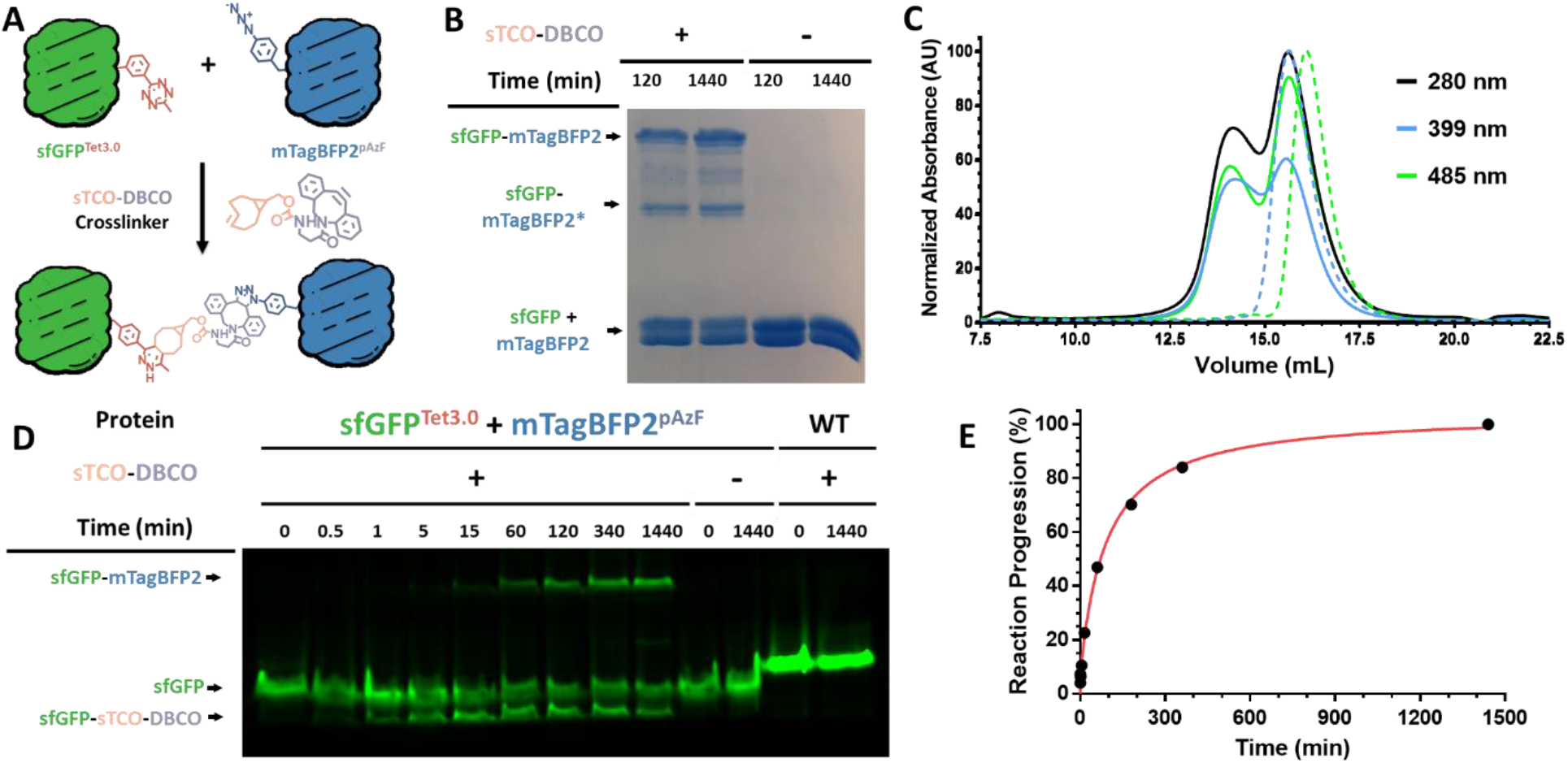
*In vitro* and *in vivo* intermolecular protein-protein crosslinking through DEAL. (A) Reaction scheme of crosslinking between sfGFP^Tet3.0^ and mTagBFP2^pAzF^ via an sTCO-DBCO linker. (B) SDS-PAGE analysis of *in vitro* reaction between purified sfGFP^Tet3.0^ and mTagBFP2^pAzF^ at 2 and 24 hours after sTCO-DBCO addition. The lower mobility crosslinked product, sfGFP-mTagBFP2^*^, matches the mass loss for photolyzed mTagBFP2^*^ known to accumulate over time^31^. (C) Size exclusion chromatogram of crosslinking reaction mixture at 24 hours post sTCO-DBCO addition and monitored by absorbance at 280 nm (black), 399 nm (mTagBFP2 λ_max_; blue), 485 nm (sfGFP λ_max_; green). Dashed lines are elution profiles of purified sfGFP^WT^ (dashed green) and mTagBFP2^WT^ (dashed blue). (D) SDS-PAGE in-gel fluorescence image of *in vivo* crosslinking reaction time course following the addition of sTCO-DBCO to *E. coli* cells containing expressed either sfGFP^Tet3.0^ and mTagBFP2^pAzF^, or sfGFP^WT^ and mTagBFP2^WT^. (E) Densitometry quantification of panel D with reaction time course shown as a percentage of maximal crosslinked product over time. The contents of each lane are indicated above each gel.

To track *in vivo* crosslinking of sfGFP^Tet3.0^ and mTagBFP2^pAzF^ we directly observed the formation of the crosslinked product over time using SDS-PAGE in-gel fluorescence^32^ (Fig. 5D-E). Immediately following the addition of sTCO-DBCO we observed a small downward mobility shift from the sfGFP^Tet3.0^ as it reacted quickly with sTCO-DBCO followed by a slower coupling reaction with mTagBFP2^pAzF^ to form the crosslinked sfGFP-mTagBFP2 complex (Fig. 5D, see supplemental discussion “*sfGFP/mTagBFP2 and sfGFP-mTagBFP2 Expression and Characterization*”). The reaction appeared to reach 50% completion at about ∼70 minutes after the addition of sTCO-DBCO to cells containing sfGFP^Tet3.0^ and mTagBFP2^pAzF^ and approached completion (∼35% total yield as determined by densitometry) after 6 hours (Fig. 5D-E). Importantly, no detectable crosslinking was observed in the absence of sTCO-DBCO, or when the crosslinker was added to cells containing expressed proteins lacking ncAAs (Fig. 5D-E). These results confirm that our DEAL system can be used to achieve site-specific crosslinking between two different proteins, thereby enabling precise control over supramolecular topography in the complex milieu of live cells.

#### Intramolecular Protein Crosslinking in vivo

Our *in vivo* DEAL system should also be capable of intramolecular protein crosslinking often referred to as “protein stapling”. While peptide stapling has been used extensively to alter the properties of peptides^33^, no general protein stapling methodology has been demonstrated in live cells.

Prior to pursuing protein stapling in cells, we verified *in vitro* stapling was feasible using an sTCO-PEG_4_-DBCO crosslinker on purified, dual-encoded protein. To do so, we evaluated the extent of stapling for two sfGFP-mTagBFP2 fusion proteins (sfGFP^Tet3.0^-mTagBFP2^pAzF^ and mTagBFP2^pAzF^-sfGFP^Tet3.0^; Fig. S10) when exposed to an sTCO-PEG_4_-DBCO crosslinker (Figs. 6A-B). We found that in order to observe distinctive stapling via in-gel fluorescence the crosslinked product had to be cleaved by TEV protease at a cut site in the linker to resolve the linked and unlinked products (Figs. 6A-B, S10). Both constructs showed similar degrees of stapling, achieving ∼50% after 120 minutes (Fig. 6A-B). Due to its higher levels of expression (Fig. S10B), we elected to proceed with the sfGFP^Tet3.0^-mTagBFP2^pAzF^ orientation to explore *in vivo* stapling.

**Figure 6.**
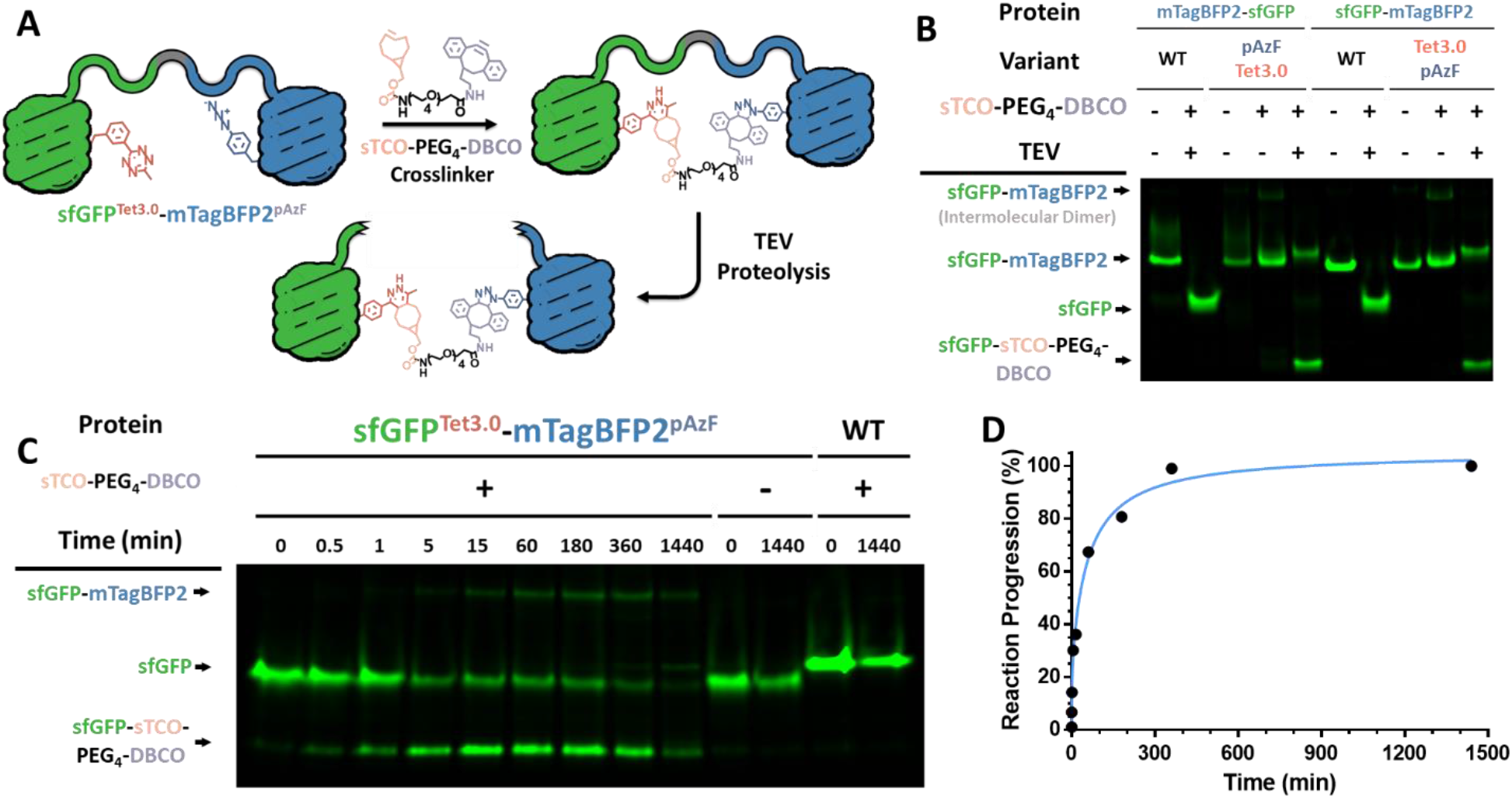
*In vitro* and *in vivo* intramolecular protein stapling via DEAL. (A) Reaction scheme of intramolecular stapling of sfGFP^Tet3.0^-mTagBFP2^pAzF^ with sTCO-PEG_4_-DBCO and subsequent TEV cleavage. (B) Analysis of *in vitro* protein stapling of sfGFP^Tet3.0^-mTagBFP2^pAzF^ and mTagBFP2^pAzF^-sfGFP^Tet3.0^ with sTCO-PEG_4_-DBCO, using SDS-PAGE fluorescent imaging. (C) SDS-PAGE in-gel fluorescence image of *in vivo* reaction time course following the addition of sTCO-PEG_4_-DBCO to *E. coli* cells containing expressed sfGFP^Tet3.0^-mTagBFP2^pAzF^ or sfGFP^WT^- mTagBFP2^WT^. (D) Densitometry quantification of panel C, reaction time course shown as a percentage of maximal crosslinked product over time. The contents of each lane are indicated above each gel.

To demonstrate *in vivo* protein stapling with our DEAL system, *E. coli* cells containing sfGFP^Tet3.0^-mTagBFP2^pAzF^ were exposed to sTCO-PEG_4_-DBCO and the degree of stapling was determined at various time points. Stapling progress, as monitored by SDS-PAGE in-gel fluorescence, showed that *in vivo* stapling reached 50% of its maximal extent (∼18% total yield) after about ∼20 minutes and was nearly complete after 6 hours (Figs. 6C-D), highlight the ability of our DEAL system to enable intramolecular “stapling” of proteins in live *E. coli* cells.

#### Dual Fluorescent Labeling in vivo

The ability to site-specifically install two different fluorescent dyes into a protein in their native context is highly desired to monitor conformational changes^5^, track localization^34^, and observe protein-protein interactions^35^, among other applications. To demonstrate *in vivo* DEAL with fluorescent dyes, we simultaneously applied DBCO-TAMRA and sTCO-JF669 dyes (1 µM) to cells containing sfGFP^Tet3.0^-mTagBFP2^pAzF^. After 60 and 5 minutes the SPAAC and IEDDA reactions were quenched and labeling was observed by SDS-PAGE in-gel fluorescence. As anticipated, we observed distinct and specific labeling with minimal cross-reactivity (Fig. 7). These results demonstrate that our DEAL system can be used to site-specifically and simultaneously dual-label proteins *in vivo* within biologically-relevant concentrations and times.

**Figure 7.**
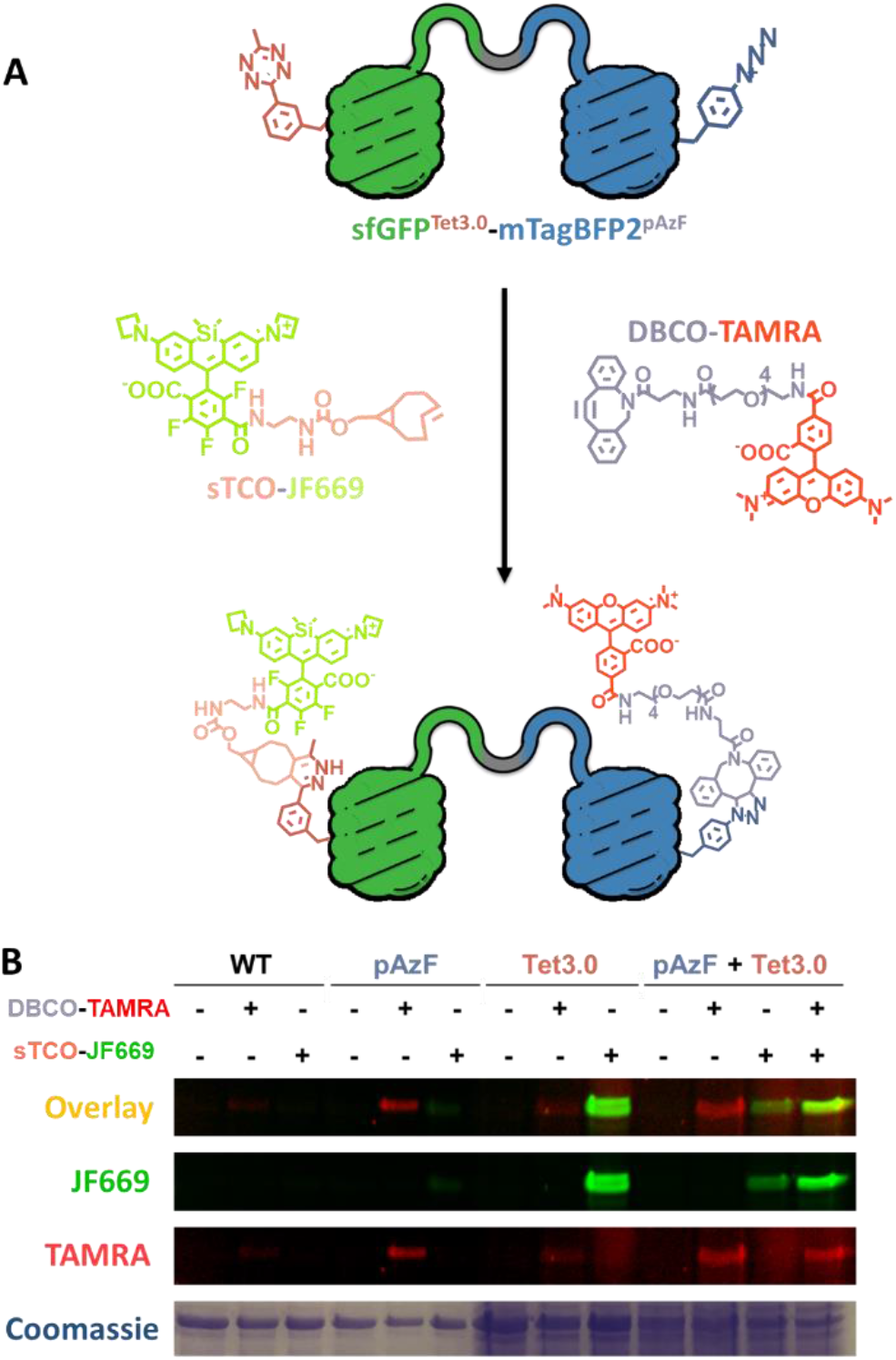
*In vivo* dual fluorophore labeling of dual encoded protein in *E. coli*. (A) Reaction scheme of dual fluorophore labeling of sfGFP^Tet3.0^-mTagBFP2^pAzF^ with sTCO-JF669 and DBCO-TAMRA. (B) SDS-PAGE of resulting *E. coli* cell lysate after *in vivo* reactions in cells containing expressed sfGFP^Tet3.0^-mTagBFP2^pAzF^ labeled with sTCO-JF669 and DBCO-TAMRA; top panel is the overlay of TAMRA and JF669 channels followed by each individual channel, and Coomassie staining. Cells exposed to fluorescent labels were quenched with excess pAzF and/or Tet3.0 prior to sample preparation.

### Dual labeling of *Saccharomyces cerevisiae* Replication Protein A DNA-Binding Domains *in vitro*

Next, we sought to determine if our DEAL system could address a more challenging and biologically relevant problem: monitoring the interaction between *Saccharomyces cerevisiae* Replication Protein A (RPA) and single-stranded DNA (ssDNA) via FRET. RPA is a heterotrimeric ssDNA-binding protein complex that is essential for almost all aspects of DNA metabolism^36^. RPA functions to sequester transiently open ssDNA in the cell and serves as a hub for the recruitment of over three dozen enzymes^37^. Structurally, RPA is composed of five ssDNA binding domains (DBDs) and two protein-protein interaction domains (PIDs) which are tethered by several flexible linkers^38^ (Fig. 8A). Owing to these linkers, RPA can adopt multiple configurations on ssDNA and experimental tools to directly monitor transitions in these configurations are currently limited. Moreover, the RPA complex is replete with Cys residues and cannot be reconstituted *in vitro*, making installation of fluorophores by DEAL a necessary approach. We used DEAL to site-specifically introduce organic dyes onto the two terminal DBDs such that the change in configuration can be monitored by FRET upon binding to ssDNA (Fig. 8B).

**Figure 8.**
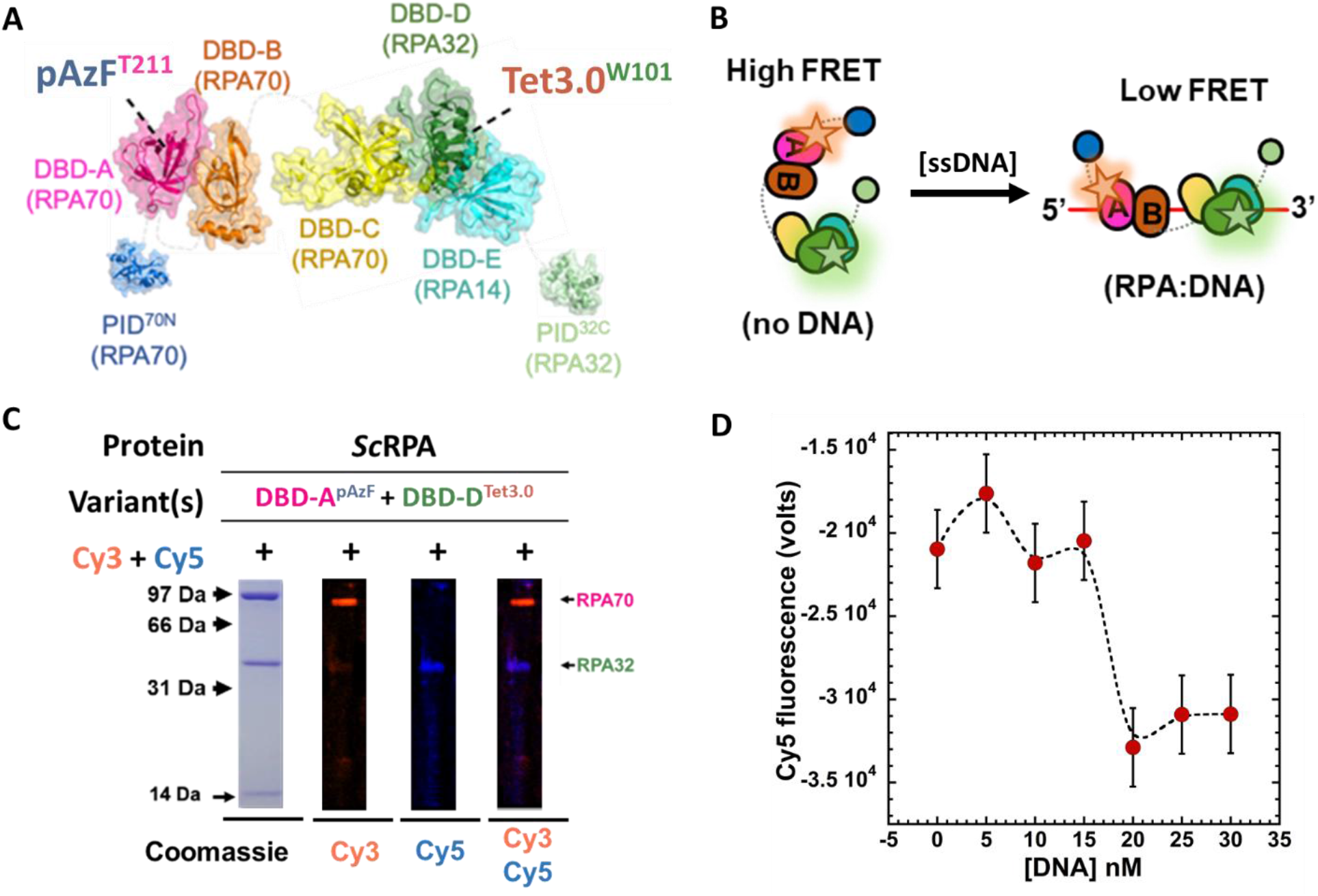
Characterization of *Sc*RPA labeling and FRET analysis of DNA binding. (A) Structural model of *S. cerevisiae* RPA complex (PDB: 6I52) highlighting the location of encoded pAzF and Tet3.0 in DBD-A and DBD-D, respectively. (B) Scheme depicting RPA binding ssDNA and the predicted FRET changes associated with these conformational states. (C) SDS-PAGE in-gel fluorescence analysis of *Sc*RPA after labeling by DBCO-Cy3 and TCO-Cy5. Each dye signal is imaged individually and as an overlay. (D) Resulting Cy5 fluorescence intensity from dual Cy3/Cy5 labeled *Sc*RPA^Dual^ as a function of DNA concentration.

To generate fluorescently dual labeled RPA, we encoded pAzF at site 211 on DBD-A (in RPA70) and Tet3.0 at site 101 on DBD-D (in RPA32)^39,40^. Following purification of full-length RPA-DBD-A^pAzF^-DBD-D^Tet3.0^ the complex was labeled with DBCO-Cy3 and TCO-Cy5. In-gel fluorescence was used to confirm DEAL of DBD-A^pAzF-Cy3^ and DBD-D^Tet3.0-Cy5^, respectively, with no detectable cross-reactivity (Fig. 8C). We next monitored FRET changes in dual-labeled RPA as a function of ssDNA concentration (Fig. 8D). In its unbound state RPA is expected to adopt an ensemble of configurations that give rise to a high FRET signal; however, upon ssDNA binding the complex is expected to stabilize into a splayed, low FRET configuration where the DBDs are linearly arranged on the ssDNA template^40^ (Fig. 8B). In agreement with this prediction, we observed high of FRET between Cy3 and Cy5 at low ssDNA concentrations and a transition to low FRET as more DNA is added (Fig. 8D). These results highlight the robustness of our DEAL system to address challenging encoding and labeling problems on meaningful protein systems.

## Conclusions

We have reported here that, when combined with GCE, the SPAAC and IEDDA bioorthogonal labeling reactions can be used to simultaneously and site-specifically label a protein in live cells. To do so, we developed and optimized a DEAL system that encodes pAzF and Tet3.0 into proteins and relies on their orthogonal reactivity to achieve labeling. Our DEAL system enables homogenous production of full-length proteins containing pAzF and Tet3.0 at ∼10% of their natural counterparts, and affords mutually orthogonal reactions *in vitro* and *in vivo* with typical labeling yields upwards of 50% and 90% respectively. We showcased the utility and versatility of this DEAL system for *in vivo* applications through three vignettes. First, through the simultaneous incorporation of pAzF and Tet3.0 into two distinct proteins within the same cell, we demonstrated the first example of *in vivo* site-specific bioorthogonal protein-protein crosslinking through the addition of a heterobifunctional crosslinker. Second, we achieved *in vivo* protein stapling by performing intramolecular crosslinking between pAzF and Tet3.0 handles located within the same protein. Third, we demonstrated efficient dual labeling in *E. coli* using two fluorescent dyes with minimal cross-reactivities. Moreover, to tackle a traditionally intractable problem, DEAL was applied to monitor the relative configurational changes of *S. cerevisiae* RPA by FRET as it binds ssDNA. These applications conclusively demonstrate that the SPAAC and IEDDA reactions are efficient and mutually orthogonal on the same protein both *in vitro* and within living *E. coli* cells.

The GCE systems were selected in this work to maximize DEAL on proteins from *E. coli* and to demonstrate the superior efficiency of combining these bioorthogonal handles on proteins. Due to the success of these applications, we hypothesize that transplanting the ncAAs and reactions developed here into eukaryotic cells using currently available systems^13,41^ will have a similar level of mutual orthogonality and provided efficient DEAL for a broad range of applications. For example, while we illustrated dual *in vivo* labeling with two fluorophores, one could also install virtually any combination of probes or moieties, such as protein ligands and inhibitors to modulate protein function, photosensitive probes for spatiotemporal control, or secondary probes with shifted spectral properties, all of which could enable simultaneous monitoring and control of protein functions in real-time in their native context^42–44^. As demonstrated, the reactive handles can be installed on separate proteins *in vivo*, which could allow one to monitor protein-protein interactions via FRET without needing bulky fluorescent proteins that may ablate interactions^45^. Likewise, DEAL could be used to construct supramolecular structures or to switch protein function in living cells through installation of crosslinks within a protein structure with residue-specific precision, similar to how we were able to achieve bioorthogonal protein-protein crosslinking and protein stapling^46,47^. While this work represents the first examples of DEAL that are site-specific and function effectively *in vivo*, we expect DEAL system utility to increase with advances in GCE technology and bioorthogonal chemistry.

## Materials and Methods

A detailed materials and methods section can be found in the supplementary materials.

## Supporting information

Dual-ncAA Supplemental (JACS Final)

## Supplementary Materials

All supporting material including methods, discussion, figures, schemes, and tables can be found in the supplemental figures.

## Author Contributions

R.M.B., R.A.M., and R.B.C. designed of the project. R.M.B. performed the experiments. The experiments specific to RPA were designed by E.A. and S.K and performed by S.K.. Organic reagents generated for the experiments were synthesized by S.J.. R.F. and J. B. analyzed the MS data. R.M.B., R.A.M., and R.B.C. wrote the manuscript with input from E.A., S.K., S.J., R.F., and J.B..

## Acknowledgements

We would like to kindly thank Dr. Luke Lavis of Janelia Research Campus, HHMI, for providing the JF669-CO_2_H used in this study. We would also like to thank Dr. Colin Johnson for use of his probe sonicator and fluorescence spectrometer.

---

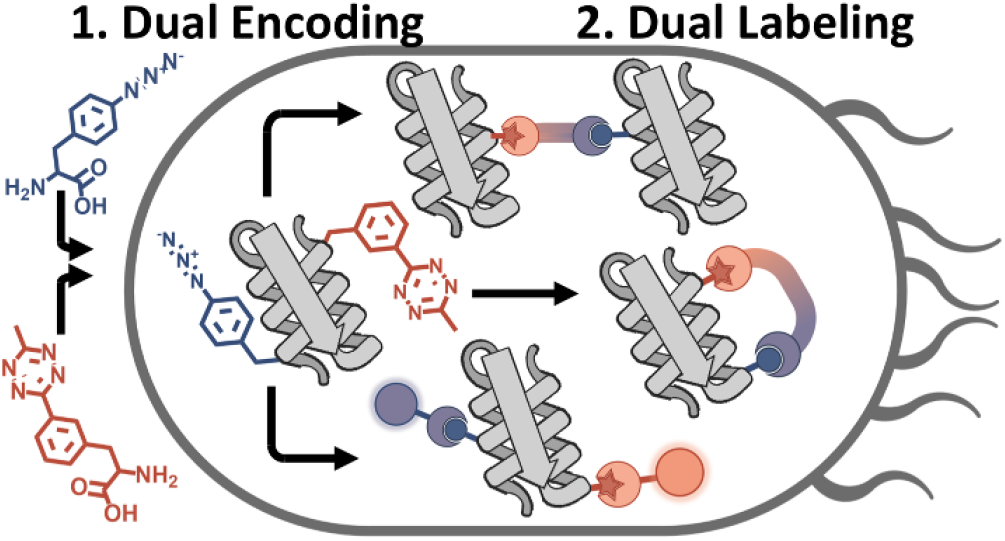

**For Table of Contents Only**

