## Supplementary material for "Genetic Incorporation of Two Mutually Orthogonal Bioorthogonal Amino Acids That Enable Efficient Protein Dual-Labeling in Cells": Dual-ncAA Supplemental (JACS Final)

**This PDF file includes:**

Materials and Methods
Supplementary Discussion
Schemes S1–S3
Figures S1 to S31
Tables S1-S6
References 1–32

### Materials and Methods

##### *General Synthetic Methods*

All purchased chemicals were used without further purification**.** DBCO-Amine, DBCO-$PEG_4$-Amine, and DBCO-TAMRA were purchased from Click Chemistry Tools. JF669-$CO_2H$ was a gift from Luke Lavis at Janelia Research Campus, HHMI. Anhydrous dichloromethane was used after overnight stirring with calcium hydride and distillation under argon atmosphere. Thin-layer chromatography (TLC) was performed on silica 60F-254 plates. The TLC spots containing alkenes were charred by potassium permanganate staining. Flash chromatographic purification was performed with silica gel 60 (230-400 mesh size). $^{1}H$ NMR spectra were recorded with Bruker 400MHz and 700 MHz, while $^{13}C$ NMR spectra were recorded at 175 MHz. Coupling constants (J values) are reported in hertz. The chemical shifts are shown in ppm and are referenced to the residual non-deuterated solvent peak $CDCl_3$ ($\delta$ =7.26 in $^{1}H$ NMR, $\delta$ = 77.23 in $^{13}C$ NMR), $CD_3OD$ ($\delta$ =3.31 in $^{1}H$ NMR, $\delta$ = 49.2 in $^{13}C$ NMR), as an internal standard. Splitting patterns of protons are designated as follows: s-singlet, d-doublet, t-triplet, q-quartet, m-multiplet, bs- broad singlet, dd- doublet of doublets.

*Chloride salt of (S)-2-amino-3-(3-(6-methyl-1,2,4,5-tetrazin-3-yl)phenyl)propanoic acid (Tet3.0, 1)*: The synthetic procedure was followed as previously described with minor alterations[1]. In a

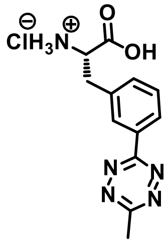

flame dried, 15 mL heavy walled reaction tube under argon atmosphere the starting material Boc-protected 3-cyano phenylalanine (250 mg, 0.86 mmol) was added, along with $Ni(OTf)_2$ catalyst (153 mg, 0.43 mmol) and acetonitrile (0.5 ml, 9 mmol). Anhydrous hydrazine (1.4 mL, 43 mmol) was slowly added to the reaction mixture while stirring and purged with argon for 5 to 10 minutes and the tube was immediately sealed. The reaction mixture was then heated to 50 $^{o}C$ for 24 hr. The reaction mixture then was cooled to room temperature, and the apparatus opened slowly to add 20 equivalents of 2 M $NaNO_2$ solution along with 5 mL of water. The aqueous phase was washed with ethyl acetate (20 ml) to remove the homo coupling product. The collected aqueous phase was then acidified with 4 M HCl (pH ~2) under ice cold temperatures and extracted with ethyl acetate (3x 20 mL). The combined organic layer was washed with brine, dried with anhydrous $Na_2SO_4$ and concentrated under reduced pressure. Silica gel flash column chromatography (30-35% ethyl acetate in hexanes with 1% acetic acid) yielded 235 mg of **Boc-protected-Tet3.0** (235 mg, 0.65 mmol) in the form of a pinkish red gummy material. Yield 76%. $^{1}H$ NMR (400MHz, $CDCl_3$) $\delta$ 8.41 (2H, t, $J$ = 7.2 Hz), 7.51-7.44 (2H, m), 5.17 (1H, d, $J$ = 7.4), 4.71 (1H, d, $J$ = 5.2), 3.33 (1H, dd, $J$ = 13.6, 5.2 Hz), 3.21 (1H, dd, $J$ = 13.2, 6.4 Hz), 3.07 (3H, s), 1.39 (9H, s). $^{13}C$ NMR (175MHz, $CDCl_3$) $\delta$ 175.6, 167.3, 164.1, 155.5, 137.7, 133.8, 132.1, 129.5, 129.1, 126.7, 80.4, 54.4, 38.1, 28.4, 21.1.

The purified Boc-protected Tet-3.0 amino acid (200 mg, 0.55 mmol) was then dissolved in 2 mL ethyl acetate and charged with 1 mL HCl gas saturated 1,4 Dioxane under argon atmosphere. The reaction mixture was allowed to stir at room temperature until the starting materials was consumed, as determined by TLC (normally 2 to 3 hours.). This product was concentrated under reduced pressure and re-dissolved in ethyl acetate (2x 10 mL) and similarly concentrated to remove excess HCl gas which resulted in pink colored solid material of **Tet3.0** in quantitative yield (97%). $^{1}H$ NMR (400MHz, $CD_3OD$) $\delta$ 8.52-8.49 (2H, m), 7.66-7.60 (2H, m), 4.38 (1H, dd, $J$ =7.2, 6 Hz), 3.46 (1H, dd, $J$ = 14.4, 5.6 Hz), 3.35 (1H, dd, $J$ = 14.4, 7.2 Hz), 3.05 (3H, s). $^{13}C$ NMR (175 MHz,

$CD_3OD$) $\delta$ 171.1, 169.1, 165.2, 137.1, 134.7, 134.4, 131.2, 129.8, 128.4, 55.1, 37.3, 21.1. ESI-MS calculated for $C_{12}H_{14}N_5O_2$ ($[M + H]^+$) 260.1142, found 260.1133.

*4-Nitro Phenyl activated sTCO (2)*: The synthetic procedure was followed as previously described[1]. $^1$H NMR (400MHz, $CDCl_3$) $\delta$ 8.27 (2H, d, *J* = 9.6 Hz), 7.37 (2H, d, *J* = 9.6 Hz), 5.88-5.82 (1H, m), 5.18-5.14 (1H, m), 4.18 (2H, d, *J* = 7.2 Hz), 2.43-2.39 (1H, m), 2.35-2.22 (3H, m), 1.96-1.90 (2H, m), 0.94-0.83 (1H, m), 0.69-0.64 (1H, m), 0.62-0.49 (3H, m).

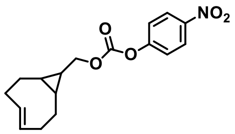

*DBCO-sTCO (3)*: In 3 mL of anhydrous dichloromethane, DBCO-Amine (10 mg, 0.036 mmol), the activated ester of sTCO **2** (14 mg, 0.045 mmol) and triethylamine (30 µL, 0.20 mmol) were combined under argon atmosphere. The reaction mixture was stirred at room temperature for 18 hours. The solvent was then concentrated onto silica gel under reduced pressure and the resulting product, molecule **3** (11 mg, 0.024 mmol), was purified by silica gel column chromatography (30-35% methanol in dichloromethane). Yield 67%. $^1$H NMR (400MHz, $CD_3OD$-$CDCl_3$-mix.) $\delta$ 7.67 (1H, d, *J* = 7.2 Hz), 7.48-7.41 (4H, bs), 7.39-7.31 (2H, m), 7.26 (1H, d, *J* = 7.2 Hz), 5.88-5.81 (1H, m), 5.15-5.07 (2H, m), 3.83 (2H, d, *J* = 6.8 Hz), 3.72 (1H, d, *J* = 14 Hz), 3.21-3.15 (1H, m), 3.11-3.02 (1H, m), 2.54-2.42 (1H, m), 2.32-2.12 (2H, m), 2.07-1.99 (1H, m), 1.94-1.84 (2H, m), 1.34-1.28 (2H, dd, *J* = 7.6, 2.4 Hz ), 0.91-0.80 (2H, m), 0.62-0.51 (2H, m), 0.41-0.34 (1H, m).

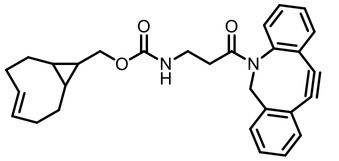

*DBCO-PEG4-sTCO (4)*: Following the above procedure, using 10 mg (0.019 mmol) of DBCO-PEG$_4$-amine, 7 mg (0.023 mmol) of the activated ester of sTCO **2** and 30 µL triethylamine (0.2 mmol) produced 9 mg (0.012 mmol) of the title molecule **4**. Yield 63%. $^1$H NMR (400MHz, $CD_3OD$-$CDCl_3$-mix.) $\delta$ 7.64 (1H, d, *J* = 6.8 Hz), 7.42 ( 4H, bs), 7.39-7.28 (2H, m), 7.25 (1H, d, *J* = 9.2 Hz), 5.87-5.79 (1H, m), 5.14-5.05 (2H, m), 3.89 (2H, d, *J* = 6.4 Hz), 3.69 (2H, d, *J* = 14 Hz), 3.63-3.55 (14H, m), 3.51 (4H, 7, *J* = 5.6 Hz), 3.25 (3H, t, *J* = 5.6 Hz), 3.19-3.12 (1H, m), 2.53-2.45 (1H, m), 2.29-2.20 (3H, m), 2.07-1.99 (1H, m), 1.93-1.85 (2H, m), 0.90-0.79 (2H, m), 0.55 (2H, m), 0.46-0.39 (1H, m).

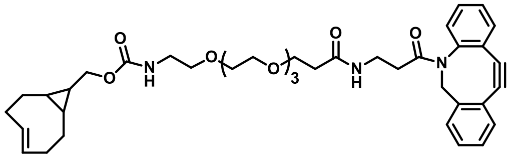

*sTCO-amine (5)*: Following the above procedure, 100 mg of the activated ester of sTCO **2** (0.315 mmol), 75 µL (1.1 mmol) of ethylenediamine and 200 µL of triethylamine (1.5 mmol) made 45 mg (0.19 mmol) of the title molecule **5**. Yield 60%. Compound was purified by silica gel column chromatography (30-35% methanol in dichloromethane). $^1$H NMR (400MHz, $CDCl_3$) $\delta$ 5.83-5.75 (1H, m), 5.09-5.01 (1H, m), 3.85 (2H, d, *J* = 6.0 Hz), 3.18 (2H, t, *J* = 5.2 Hz), 2.77 (2H, t, *J* = 6.0 Hz), 2.30-2.26 (1H, m), 2.21-2.12 (3H, m), 1.87-1.80 (2H, m), 0.79-0.74 (1H, m), 0.50-0.43 (2H, m), 0.37-0.34 (2H, m).

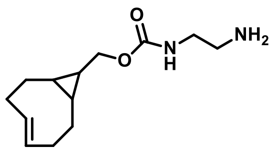

*sTCO-JF669 (6)***:** In dry DCM (2 mL), JF669-$CO_2$H (5 mg, 0.01 mmol), 1-Ethyl-3-(3-dimethylaminopropyl) carbodiimide (3 mg, 0.015 mmol) and N-hydroxysuccinimide (2.0 mg, 0.015 mmol) were added under argon atmosphere and stirred for 15 minutes under ice cold conditions. Next, sTCO -amine **5** (4 mg, 0.015 mmol) was added, followed by N,N-diisopropylethylamine (20 µl, 0.15 mmol). After 15 minutes the ice bath was removed and stirring was continued for another 24 hours at room temperature. The solvent was concentrated onto silica gel under reduced pressure and the title compound **6** (4 mg, 0.005 µmol) was purified by silica gel column chromatography (20-25% methanol in dichloromethane). Yield 53%. $^1$H NMR (400MHz, $CD_3OD$) $\delta$ 6.90 (2H, d, $J$ = 8.8 Hz), 6.78 (2H, d, $J$ = 2.4 Hz), 6.37 (2H, dd, $J$ = 6.0, 2.0 Hz), 5.89-5.81 (1H, m), 5.16-5.04 (1H, m), 4.03 (6H, t, $J$ = 7.2 Hz), 3.91 (2H, t, $J$ = 6.8 Hz), 3.76 (1H, d, $J$ = 6.8 Hz), 3.67 (1H, s), 3.49 (2H, t, $J$ = 6.0 Hz), 3.23-3.15 (2H, m), 2.46-2.39 (2H, m), 2.34 (1H, t, $J$ = 6.4 Hz), 2.27-2.19 (4H, m), 1.95-1.85 (2H, m), 1.82-1.73 (1H, m), 0.91-0.86 (2H, m), 0.55 (3H, s ), 0.51 (3H, s ), 0.46-0.39 (2H, m), 0.27-0.21 (1H, m).

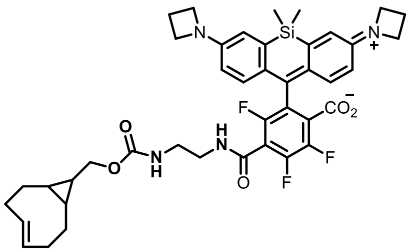

***Molecular Cloning***

Molecular cloning of plasmids used in this study was carried out using a combination of overlap extension PCR[2], QuikChange PCR[3], and SLiCE cloning[4]. In short, fragments containing at least 25 bp of homology at their flanking ends to either the vector backbone or other fragments were amplified using touchdown PCR[5], the resulting products were separated on 0.8-1.2 % (w/v) agarose gels, and purified using GeneJet Gel Extraction Kit (ThermoFischer Scientific, USA) according to the manufacturer's instructions. Vector backbones were prepared through either restriction digestion or PCR amplification and similarly purified through gel extraction, as previously mentioned. Fragments and vector backbones were then ligated using the SLiCE cloning protocol[4] and transformed into chemically-competent DH10B *E. coli* cells and selected on LB agar plates containing the appropriate antibiotic. In several cases, multiple fragments were fused using overlap extension PCR prior to ligation by SLiCE (see Table 1). For codon reassignment (i.e. for generation of pDule1-Tet3.0$^{TAA}$), anticodon sequences were mutated via QuikChange PCR followed by DpnI digestion prior to transformation into *E. coli* DH10B cells. Colonies were selected and propagated prior to purification. Genetic sequences of each plasmid were confirmed using Sanger sequencing. The primers and template used for PCR amplification, as well as the vector backbone and restriction enzymes used for linearization are summarized for each plasmid in Tables 1 and 3. Plasmids not prepared in this study were acquired either through Addgene (see Addgene ID number in Table 1) or were prepared previously (see references in Table 1). It is important to note that all genes that were cloned were done so such that they were terminated by an opal stop codon.

To generate the strains listed in Table 2, ~50 ng of each plasmid was combined with either chemically-competent DH10B, or electro-competent BL21(DE3) *E. coli* cells. The mixture was incubated for approximately 15 minutes prior to transformation; for chemically-competent cells, the mixture was incubated at 42°C for 45 seconds; for electro-competent cells, the mixture was electroporated using an Eporator electroporator (Eppendorf, KGaA, Germany; formerly New Brunswick Scientific, USA). Freshly transformed cells were immediately resuscitated in 2xYT media for approximately one hour at 37°C at 250 rpm, and plated on LB agar containing the proper antibiotics, and incubated overnight at 37°C. DH10B strains were stored as frozen glycerol stocks at -80°C, whereas BL21(DE3) strains were transformed fresh and stored at 4°C for no more than two weeks. Note that in general strains containing pBAD vectors were prepared in DH10B cells[6], while T7-based vectors (pET and pRSF) were prepared in BL21 cells[7] since this cell line possesses the trans-acting T7 polymerase to drive expression of these reporters.

***Culture Conditions***

Unless otherwise stated cultures were grown in sterile baffled culture flasks at 50 mL scale in an I26 incubator-shaker (Eppendorf, KGaA, Germany; formerly New Brunswick Scientific, USA) set to 37°C and 250 rpm in autoinduction media (AIM; prepared as detailed by Studier[8]) supplemented with arabinose (0.05% w/v) and lactose (0.02% w/v) and appropriate antibiotics for approximately 24 hours. 5 mL overnight starter cultures in noninducing media[8] supplemented with the appropriate antibiotics were used to inoculate expression cultures at 2% (v/v) dilution. For BL21(DE3) strains overnight starter cultures were inoculated by scraping a swath of cells from a fresh LB agar plate. For DH10B strains, overnight cultures were inoculated by picking a single colony from a LB agar plate, or by inoculation from a frozen glycerol stock. 100 mM Tet3.0 and pAzF stock solutions were prepared fresh in DMF and $H_2O$ with 2 equivalents of NaOH, respectively, and were diluted into cultures at 0.5 and 1.0 mM, respectively prior to inoculation by overnight starters.

***Determination of Suppression Efficiency***

For the determination of suppression efficiency, strains were grown in triplicate at the 500 µL scale in sterile 96-well blocks in a similar fashion as to above. To measure culture OD, a 20 µL aliquot was removed and diluted into 200 µL of water in a 96-well plate and the absorbance at 600 nm was determined using a Synergy 2 Microplate Reader (BioTek). To measure fluorescence, a 50 µL aliquot of culture was removed and diluted into 200 µL of water in a 96-well plate and the sfGFP fluorescence was measured on the same microplate reader using a 485/20 excitation filter and 528/20 emission filter with the optics positioned at the top 50%. For mTagBFP2, fluorescence was measured using a 380/20 excitation filter and 460/40 emission filter. To determine suppression efficiency, the fluorescent signal is first normalized to OD (Fluorescence/OD) for each culture and then averaged across the three replicates. The mean normalized fluorescence for cultures grown in the presence of ncAA(s) is then corrected by subtracting the mean normalized fluorescence of corresponding cultures grown in the absence of ncAA(s). The suppression efficiency is determined as a percentage of the corrected mean normalized fluorescence relative to the mean normalized fluorescence of the corresponding WT cultures.

***Protein Expression and Purification***

Proteins were expressed according to the culture conditions stipulated in the "*Culture Conditions*" section. After growth, cultures were harvested by pelleting at ~5,500 rcf for 10 minutes. The culture supernatant was removed and pelleted cells were either used immediately or were stored frozen at -80°C until needed. To lyse, pelleted cultures were resuspended to 5 mL in 50 mM $Na_2PO_4$, 500 mM NaCl, 5 mM imidazole pH 7.0 wash buffer. These resuspended cells were then microfluidized in a single pass at 18,000 psi using a M-110P microfluidizer system (Microfluidics Corp, USA). The resulting lysate was cleared by centrifugation at ~21,000 rcf for 30 minutes in a chilled centrifuge. The cleared lysate was combined with 250-500 µL of TALON cobalt NTA resin (Takara Bio, Japan) and incubated with agitation at 4°C for at least 1 hr. The resin was then washed using 30 mL (60-120 column volumes) of wash buffer and was eluted with 2.5 mL of 50 mM $Na_2PO_4$, 500 mM NaCl, 250 mM imidazole pH 7.0 elution buffer. The eluted protein solution was desalted into PBS buffer (50 mM $Na_2PO_4$, 100 mM NaCl, pH 7.0) using a PD-10 delating column according to the manufacturer's instructions. If necessary, the protein solution was concentrated by using a 10 kDa MWCO Vivaspin spin-concentration filter (GE Health Sciences). Protein concentration was determined by absorbance at 280 using the molar extinction coefficients in table S5. Purified protein was stored at 4°C until needed.

***SDS-PAGE in-gel Fluorescence and Densitometry***

Unless otherwise stated, samples were prepared at 2 µg for purified protein, or 15 µL at an OD of 10 for crude lysate in PBS buffer, and combined with 4x SDS sample buffer (250 mM Tris, 10% (w/v) sodium dodecyl sulfate, 50% (v/v) glycerol, 20% (v/v) β-mercaptoethanol, 0.1% (w/v) bromophenol blue, pH 6.8) prior to incubation at 95°C for approximately 5 minutes. When preparing samples for sfGFP in-gel fluorescence experiments, this boiling step was omitted. Samples were then loaded onto 15% SDS-PAGE gels and run at 200 V for approximately 60 minutes. For in-gel fluorescence detection, the SDS-PAGE gel was imaged using a ChemiDoc XRS+ imager (Bio-Rad Laboratories Inc., USA) using the following light source:emission filters. For sfGFP in-gel fluorescence, Epi-blue(450-490 nm):530/28. For TAMRA dyes, Epi-green(520-545 nm):605/50. For JF669 dye, Epi-red(625-650 nm):695/55. Automatic exposure times were determined by the instrument to optimize faint band detection. Following imaging, SDS-PAGE gels were soaked in Coomassie stain (50% (v/v) methanol, 40% (v/v) water, 10% (v/v) acetic acid, 0,02% (w/v) Coomassie G250) for approximately 20 minutes followed by an approximately 20 minute soak in destain solution (50% (v/v) methanol, 40% (v/v) water, 10% (v/v) acetic acid). When necessary, densitometry measurements were made using the ImageJ software suite.

***Protein Mass Spectrometry***

Proteins were purified as previously stated (see methods section "*Protein Expression and Purification*"). After purification, proteins were de-salted two times into liquid chromatography-mass spectrometry (LC-MS) grade ultrapure water using either a PD-10 or NAP-5 de-salting column, and when necessary, spin-concentrated to a concentration above 15 µM using a 2 mL, 10 kDa MWCO centrifugal filter (Millipore). Prior to mass analysis, all proteins were diluted to 10µM before injection onto PLRP-S 1000A reverse phase chromatography column using an AdvanceBio LC system. The proteins were eluted from the column with a gradient of water/acetonitrile + 0.1% formic acid. An Agilent 6545XT Q-ToF mass spectrometer was used for in-tact mass analysis. Proteins were ionized with a Dual AJS electrospray ionization source and then directed into the mass spectrometer inlet. The ions were detected in high-resolution mode (2GHz) in an extended mass range up to 3200m/z. The Agilent MassHunter BioConfirm Software was used for deconvolution of the resulting spectra.

***In vitro Labeling***

Unless otherwise stated, proteins were labeled in 15 µL reactions in 1.5 mL polypropylene microcentrifuge tubes at room temperature and visualized by SDS-PAGE (see section "SDS-PAGE in-gel Fluorescence and Densitometry" for details of how SDS-PAGE gels were run). In general, protein concentrations ranged from 8-10 µM, with small molecule reactants being added at 10-fold molar equivalencies (excess), unless otherwise stated. For Tet3.0-sTCO reactions, the reaction was allowed to proceed for 15 minutes before being quenched with excess Tet3.0 (~6-7 mM). For pAzF-DBCO reactions, the reaction was allowed to proceed for 2-24 hours before being quenched with excess pAzF (~6-7 mM). Unless otherwise stated, when performed simultaneously, pAzF-DBCO reactions were initiated prior to Tet3.0-sTCO reactions in a one-pot format, while quenching was performed simultaneously at the end of the reaction period. ULP1 cleavage was carried out as previously described[9]

*Quantification of in vitro Labeling*

To quantify *in vitro* labeling, we adapted an approach originally developed by Murrey et al.[18]. In short, 10 µM sfGFP was exposed to DBCO-$NH_2$ and/or sTCO-OH (66.7 µM) for 24 hours and 15 minutes, respectively (unless otherwise stated) at room temperature. Following this initial non-fluorescent labeling phase, DBCO-TAMRA and/or sTCO-JF669 dyes were added (666.7 µM) as quenchers and allowed to react for 120 minutes at room temperature. To prevent inadvertent cross-reactivity on $sfGFP^{Dual}$ under single-labeling conditions during processing (i.e. sTCO-JF669 reacting with free pAzF during boiling, when Tet3.0-sTCO-OH reactivity is being assessed), excessive inverse-cognate ncAA (i.e. addition of free pAzF when assessing IEDDA reaction efficiency to minimize pAzF-sTCO-JF669 labeling) was added (6.67 mM) at this step as well. Samples were then boiled for 5 minutes in loading buffer and approximately 2.8 µg of protein was loaded onto a 15% SDS-PAGE gel, and electrophoresis was applied at 120 V for approximately 65 minutes to remove excess fluorophores. In-gel fluorescence was then detected using a Bio-Rad ChemiDoc imaging system (Bio-Rad Laboratories, Inc., USA) using the following channels: TAMRA (Ex: Epi-green (520-545 nm), Em: 605/50 nm), JF669 (Ex: Epi-red (625-650), Em: 695/55 nm) set to auto-exposure time setting. The resulting images were then quantified using densitometry (see section "SDS-PAGE in-gel Fluorescence and Densitometry"). Quantified fluorescent band intensities were corrected for non-specific labeling by subtraction of a $sfGFP^{WT}$ sample exposed to the dye in question. The relative yield was determined using the following equation:

$$Yield\ (\%) = (1 - \left(\frac{I_{block}}{I_{quench}}\right)) \cdot 100$$

Where $I_{block}$ is the band intensity of a sample that was exposed first to a non-fluorescent label (i.e. DBCO-$NH_2$ and/or sTCO-OH) prior to fluorescent quenching, and $I_{quench}$ is the band intensity of a sample that was exposed exclusively to fluorescent quenching (with no blocking).

***In vivo Labeling***

*Cell Preparation and General Labeling Conditions*

Cultures expressing ncAA-containing proteins were cultured as previously described (see section "*Culture Conditions*"). After approximately 24 hours of growth, cultures were harvested by centrifugation at 4000 rcf for 10 minutes in a chilled centrifuge. To remove excess ncAA, the resulting supernatant was decanted and the cell pellet resuspended in PBS buffer, and centrifuged for 5 minutes at 4000 rcf. This process was repeated two additional times for a total of three washes. Following washing, cells were resuspended in PBS and to an OD of 10 and were stored at 4°C until needed, being stored no longer than 24 hours. In general, all *in vivo* labeling experiments were performed on resuspended cells at an OD of 10 (unless otherwise stated), with labeling carried out at room temperature at analogous concentrations, times, and orders of addition to *in vitro* labeling. After labeling, quenching was initiated by the addition of excessive ncAA (pAzF and/or Tet3.0 when necessary), and cells were lysed by probe sonication with the amount loaded being determined so as to be consistent on the basis of sfGFP fluorescence.

*Quantification of in vivo Labeling*

Cultures expressing ncAA-containing proteins were harvested, washed, and resuspended as previously described. DBCO-$NH_2$ and/or sTCO-OH (66.7 µM) were added to 100 µL cell suspensions at OD 10, and allowed to react for 24 hours and 15 minutes, respectively at room temperature, as previously described. The resulting cell suspensions were then lysed using a probe sonicator, and DBCO-TAMRA and/or sTCO-JF669 (666.7 µM) and Tet3.0 and/or pAzF (6.67 mM) were added and allowed to react for 120 minutes at room temperature, as was performed for *in vitro* quantification. Sample OD's were then adjusted on the basis of previously determined protein yield, and boiled for 5 minutes prior to electrophoresis via SDS-PAGE as previously described. Samples were then imaged and quantified analogously to what was done for *in vitro* quantification.

*Dual Dye Labeling*

Following harvesting and washing, cultures expressing sfGFP$^{WT}$-mTagBFP2$^{WT}$ and sfGFP-mTagBFP2$^{pAzF}$ (at sites 363) were diluted to an OD of 2.5, while cultures expressing sfGFP$^{Tet3.0}$-mTagBFP2 (at site 150) and sfGFP$^{Tet3.0}$-mTagBFP2$^{pAzF}$ (at sites 150 and 363, respectively) remained at an OD of 10 to facilitate consistent loading of the sfGFP-mTagBFP2 protein. To these samples, sTCO-JF669 and DBCO-TAMRA were added at a final concentration of 1 µM and the reaction was allowed to proceed for 60 minutes. The reaction between sTCO-JF669 and Tet3.0 was quenched after 5 minutes by the addition of excessive Tet3.0, while the reaction between DBCO-TAMRA and pAzF was quenched at 60 minutes by the addition of excessive pAzF. Loading buffer was added and samples were boiled as previously described. To achieve consistent loading, samples containing sfGFP$^{WT}$-mTagBFP2$^{WT}$ and sfGFP-mTagBFP2$^{pAzF}$ were diluted a further 2-fold prior to loading. Samples were run and imaged in the TAMRA and JF669 channels before being stained in Coomassie (see section "SDS-PAGE in-gel Fluorescence and Densitometry").

*Intermolecular Crosslinking*

Cultures expressing sfGFP$^{Tet3.0}$ (at site 150) and mTagBFP2$^{pAzF}$ (at site 105) or their WT equivalents were harvested and washed as mentioned previously. sTCO-DBCO crosslinking agent was added to 5 mL of cell suspension at a 1:1 ratio to sfGFP (typically 20 µM, with DMSO at a final v/v concentration of 3%), with the necessary amount being determined empirically using an sfGFP fluorescence dequenching titration assay (analogous to the method described previously by Blizzard et al.[10]) at the time of experimentation using the current batch of cells containing sfGFP$^{Tet3.0}$. As controls, an equivalent volume of sTCO-DBCO or vehicle (DMSO) as added to WT and ncAA-protein-containing cultures, respectively. 250 µL aliquots were collected at various time points and quenched by the addition of excess pAzF and flash frozen in liquid $N_2$ until needed. All samples were then run simultaneously on a single SDS-PAGE gel without boiling, imaged in the sfGFP channel, and quantified (see section "SDS-PAGE in-gel Fluorescence and Densitometry").

*Intramolecular Crosslinking*

Following harvest, cultures expressing sfGFP-mTagBFP2$^{WT}$ were diluted to an OD of 2.5, while those expressing sfGFP$^{Tet3.0}$-mTagBFP2$^{pAzF}$ (at sites 150, and 363, respectively) remained at an OD of 10. Excess sTCO-$PEG_4$-DBCO was then added to each 5 mL cell suspension to a final concentration of 20 µM (final DMSO v/v concentration at 3%), and 250 µL aliquots were withdrawn at various time points, quenched with excess pAzF, then flash frozen in liquid $N_2$ and stored until needed. After the time course was complete, all samples were thawed, lysed by sonication, and centrifuged at 20,000 rcf for 15 minutes to remove the insoluble fraction. TEV protease was added to the clarified lysate to a final concentration of 40 µg/mL, and was digested for approximately 24 hours at 4°C. Samples containing sfGFP-mTagBFP2$^{WT}$ were then diluted a further 8-fold to achieve consistent loading of the fusion protein. Digested samples were then run simultaneously on a single SDS-PAGE gel without boiling, imaged in the sfGFP channel, and quantified (see section "SDS-PAGE in-gel Fluorescence and Densitometry").

#### *Characterization of RPA*

##### *Plasmids and Strains for RPA Purification and Dual Labeling*

Three plasmids were sequentially transformed into *E. coli* BL21Ai competent cells in three steps: pUltra1-Tet3.0$^{TAA}$ was first transformed and competent cells were generated. pEVOL-pAzF$^{TAG}$ was next transformed into cells from step 1 and competent cells, now carrying both plasmids, were generated. Finally, plasmid pRSF-Duet-ScRPA-T211TAG-W101TAA, was transformed into cells generated in step 2. A codon-optimized version of the three RPA genes was synthesized (Genscript Inc.) and encode for a C-terminal poly-histidine tag on RPA32. The amber suppressor codon (TAG) and site for pAzF incorporation is positioned corresponding to T211 in the RPA70 subunit[11]. Similarly, the ochre stop codon (TAA) and site for Tet 3.0 incorporation is positioned corresponding to W101 in the RPA32 subunit[12]. Transformants carrying all three plasmids were isolated and used for protein expression.

##### *RPA purification*

Dual-labeled RPA was generated based on our procedures developed for singly-labeled *Saccharomyces cerevisiae* RPA[11]. Briefly, 3 L of ncAA-containing auto-induction media[8] cultures were grown for each protein preparation. Cells were induced by adding 0.4 mM IPTG and 0.05% (w/v) L-arabinose, along with 1 mM pAzF, and 0.5 mM Tet3.0 when the cultures reached $OD_{600}$ of ~2.0 and grown for an additional 3 hr at 37 °C. Harvested cells were resuspended in 75 mL cell resuspension buffer (30 mM HEPES, pH 7.8, 300 mM KCl, 0.1 mM EDTA, protease inhibitor cocktail, 1 mM PMSF, 10% (v/v) glycerol and 10 mM imidazole). Cells were lysed using 300 mg/ml lysozyme followed by sonication. Clarified lysates were fractionated on a $Ni^{2+}$-NTA agarose column. Protein was eluted using cell resuspension buffer containing 400 mM imidazole. Fractions containing RPA were pooled and diluted three-fold with buffer $H^0$ (30 mM HEPES, pH 7.8, 0.1 mM EDTA, 0.02% Tween-20 and 10% (v/v) glycerol). The diluted protein sample was then fractionated over a Q-sepharose column equilibrated with buffer $H^{100}$ (buffer $H^0$ with 100 mM KCl). Protein was eluted with a linear gradient $H^{50}$ – $H^{100}$ (superscript denoting final KCl concentration in the buffer). Fractions containing RPA were pooled and diluted with $H^0$ buffer to match the conductivity of buffer $H^{100}$, and further fractionated over a Heparin column. Protein was eluted using a linear gradient $H^{100}$–$H^{1500}$, and fractions containing RPA were pooled and concentrated using an Amicon spin concentrator (30 kDa molecular weight cut-off). RPA was further purified on a S200 size exclusion column using RPA storage buffer (30 mM HEPES, pH 7.8, 30 mM KCl, 0.02% Tween-20, 0.2mM EDTA and 10% (v/v) glycerol). Purified RPA was next labeled with DBCO-Cy3 on DBD-A$^{pAzF}$ and TCO-Cy5 on DBD-D$^{Tet3.0}$. Approximately 500 nM RPA was mixed with a 2-fold molar excess of DBCO-Cy3 and TCO-Cy5 each (Click Chemistry Tools Inc.) in 1 ml reactions in labeling buffer (30 mM HEPES, pH 7.8, 300 mM KCl,

0.02 % Tween-20, 0.25 mM EDTA and 10 % (v/v) glycerol). The reaction was incubated overnight (~16 hours in the dark, at 4 °C). The labeled protein was separated from free dye using Biogel P4 column and storage buffer. The labeled protein was then flash frozen using liquid nitrogen, and stored at –80 °C. RPA concentration, and labelling efficiency were measured as previously described[12,13] using $A_{280}$ of 98500 $M^{-1}$ $cm^{-1}$ for RPA, $A_{550}$ of 150,000 $M^{-1}cm^{-1}$ for Cy3, and $A_{650}$ of 255,000 $M^{-1}cm^{-1}$ for Cy5.

*FRET experiments*
DNA binding to dual-labeled RPA was measured using a PTI QM40 fluorimeter (Horiba Scientific, Edison, NJ, USA). 100 nM RPA-$A^{Cy3}$-$D^{Cy5}$ was mixed with increasing concentrations of DNA (0-120 nM $(dT)_{35}$ oligonucleotide) in reaction buffer (30 mM HEPES, pH 7.8, 100 mM KCl, 5 mM MgCl2, 1 mM β-mercaptoethanol and 6% (v/v) glycerol). FRET changes between the two ends of RPA was monitored by exciting the Cy3 at 535 nm and measuring Cy5 emission at 645 nm, at 25 °C. The data were collected as an average of 30 replicates.

### Supplementary Discussion

***Evaluation of UAG and UAA Single Site Suppression***
Before combining all components to generate a dual suppression system, we first sought to establish which genetic combinations maximized single nonsense suppression efficiency. Efficiency and orthogonality within GCE systems are dependent on the relative concentrations and activities of each of the GCE components during translation. Key factors that dictate the concentrations of the GCE genetic components are the plasmid copy number, the promotor, and the gene dosage. The importance of these factors, inspired us to evaluate how these factors affect two-site incorporation by genetic code expansion. To optimize dual-ncAA incorporation, we investigated the compatibility of several combinations of vectors for housing the various GCE genetic components under both single and dual suppression contexts. The pAzFRS/$tRNA_{CUA}$ system was housed on either a pEVOL[14] or pUltraI[15] plasmid, while the Tet3.0RS/$tRNA_{UUA}$ system was housed on either a pUltraI or pDule1[10] plasmid. As a fluorescent reporter of suppression efficiency in these experiments, we used a SUMO-sfGFP (Small Ubiquitin-like Modifier-super-folder Green Fluorescent Protein)[9] fusion protein containing a stop codon (either UAG or UAA) at position 35 in the SUMO domain. This site was chosen since it minimizes tetrazine-dependent fluorescent quenching of sfGFP, which could lead to an underestimation of suppression efficiency[10,16]. Each subsystem plasmid was paired with a UAG/UAA-interrupted SUMO-sfGFP reporter contained on either a pBAD or pET28 plasmid. The aforementioned plasmid combinations were assembled in either DH10B or BL21(DE3); however, due to strain-specific requirements, all combinations involving pBAD plasmids were assessed in DH10B cells, while all combinations involving pET28 plasmids were assessed in BL21(DE3) cells.

After expression, the amount of SUMO-sfGFP fluorescence produced was measured in the presence or absence of the corresponding ncAA and compared to an uninterrupted wild-type reporter (SUMO-$sfGFP^{WT}$). Robust UAG codon suppression efficiencies of 70% - 80% relative to SUMO-$sfGFP^{WT}$ was achieved when the pAzFRS/$tRNA_{CUA}$ pair was contained on either pEVOL or pUltraI plasmid in combination with a pET vector in the presence of pAzF (Fig S2A). In this case, we also noted a high level of background fluorescence for the pAzFRS/$tRNA_{UUA}$ in the absence of ncAA. Interestingly, a substantial drop in pAzF suppression efficiency was observed when the reporter was expressed from a pBAD vector in DH10B cells, highlighting the need to evaluate different plasmid and cell combinations when optimizing dual suppression systems (Fig. S2A). In the case of combinations involving the pAzFRS/tRNA$_{CUA}$ on pEVOL plasmids, we failed to observe expression even from our sfGFP$^{WT}$ reporter, indicating that this combination of plasmids is not conducive to heterologous protein expression. Similarly, for UAA suppression with Tet3.0, the greatest ncAA-dependent suppression (~21%) was observed from the Tet3.0RS/tRNA$_{UUA}$ system when on pUltraI and paired with a pET28 vector containing the reporter gene, while other combinations produced marginal, or no difference from background (Fig. S2B).

From these evaluations we surmised that optimal dual encoding would result from a plasmid combination with the gene of interest on a pET28 plasmid, the pAzFRS/tRNA$_{CUA}$ subsystem contained on either pEVOL or pUltraI vectors, and the Tet3.0RS/tRNA$_{UUA}$ subsystem contained on pUltraI. Due to plasmid compatibilities our finalized dual suppression system comprises of three-plasmids: (1) the pAzFRS/tRNA$_{CUA}$ subsystem on a pEVOL vector (pEVOL-pAzF$^{TAG}$), (2) the Tet3.0RS/tRNA$_{UUA}$ subsystem on a pUltraI vector (pUltraI-Tet3.0$^{TAA}$), and (3) the dual nonsense-encoding gene of interest on a pET28 vector (Fig. S1).

***Evaluation of GCE orthogonality***

As mentioned above and in the main text, we have observed several instances where the orthogonality of our dual-ncAA systems appear to be compromised, such as higher background fluorescence in the absence of ncAA for the pAzFRS/tRNA$_{CUA}$ system (Fig. S2A), and production of SUMO-sfGFP in the absence of the pAzFRS/tRNA$_{CUA}$ system (Fig. 2A). As such, we found it necessary to investigate these phenomena, even if they only manifest under expression conditions where one or more components of the dual-ncAA suppression system were absent. It has previously been demonstrated by others that the *Mj*TyrRS/tRNA$_{CUA}$ and *Mb*PylRS/tRNA$_{UUA}$ pairs are orthogonal on the aaRS-tRNA level, and as such, we did not assess the orthogonality of our systems at this level[15,17,18]. Nevertheless, orthogonality compromises have been observed at the ncAA-aaRS and tRNA-Codon levels[15].

A high background fluorescence for the pAzFRS/tRNA$_{CUA}$ could be indicative of (1) either higher levels of endogenous near-cognate suppression from the host decoding systems at the designated codon, or (2) of erroneous charging the orthogonal tRNA by either endogenous aaRSs or (3) misacylation of the orthogonal tRNA with natural amino acids on the behalf of the pAzFRS. If the latter case is true, this could mean that the pAzFRS exhibits polyspecificity towards alternative substrates other than pAzF, and as such, could potentially recognize Tet3.0 as a substrate as well. To test these possibilities, we paired each suppression system with a SUMO-sfGFP containing a cognate nonsense codon and observed the production of fluorescence in the presence of either pAzF, Tet3.0, or neither ncAA (Fig. S3A). We observed that, for the most part, each system seemed to produce robust fluorescence only in the presence of its cognate ncAA (Fig. S3A). Nevertheless, while not as intense as previously observed, the pAzFRS/tRNA$_{CUA}$ seemed to produce slightly more SUMO-sfGFP in the absence of ncAA than in the presence of Tet3.0, making it unlikely that pAzFRS can recognize Tet3.0 as a substrate. To further investigate the cause of the higher background fluorescence associated with the pAzFRS/tRNA$_{CUA}$ analyzed sfGFP$^{TAG}$ that was produced by the pAzFRS/tRNA$_{CUA}$ system in the presence of pAzF, Tet3.0, or in the absence of ncAA (Fig. S3B). What we observed was that when sfGFP$^{TAG}$ was suppressed through the action of the pAzFRS/tRNA$_{CUA}$ system in the presence of its preferred substrate, this protein exhibits a clean profile consistent with homogenous pAzF incorporation, with all other major peaks explained as either salt adducts, or through loss of N-terminal methionine[19] (Fig.

S3B). However, when sfGFP$^{TAG}$ is suppressed by the pAzFRS/tRNA$_{CUA}$ in the presence of Tet3.0 or absence of ncAA, we observed very similar spectra that differed in the identity of the ncAA at position 150 (Fig. S3B). Inspection of the dominant peak reveals that it possesses a mass identical to that of sfGFP$^{WT}$, indicating that either Asn, Asp, Leu, Ile, or a mixture of these amino acids are present at the 150 site (Fig. S3B). We also observed a minor peak that seemed to be consistent with Phe incorporation in both of these spectra (Fig. S3B). Since the pAzFRS active site was evolved from a TyrRS to recognize multiple aromatic ncAAs with modifications at the *para* position of Phe[20], we find it unlikely that this aaRS can recognize Asn, Asp, Leu, or Ile, and as such, the incorporation of these amino acids is likely due to near-cognate suppression from endogenous systems[21]. Incorporation of Phe, on the other hand, can likely be explained by recognition of this amino acid as an alternative substrate for the pAzFRS, which indicates that it still retains specificity for its ancestral aryl scaffold. Indeed, others have observed Phe incorporation using this and other aaRSs derived from the *Mj*TyrRS in the absence of ncAA[22,23]. We conclude that the abnormally high background associated with the pAzRRS/tRNA$_{CUA}$ pair can be explained through a combination of near-cognate suppression and misacylation of tRNA$_{CUA}$ with Phe by the pAzFRS, and note that in the presence of the preferred substrate, homogenous pAzF incorporation results.

We also elected to assess the orthogonality of our systems at the tRNA-Codon level since we observed production of SUMO-sfGFP$^{Dual}$ when only the Tet3.0RS/tRNA$_{UUA}$ system is present (Fig. 2A). The most parsimonious explanation for this observation is that the tRNA$_{UUA}$ is capable of suppressing both its cognate UAA codon, as well as the UAG codon through near-cognate suppression. To test this hypothesis, we combined each suppression pair individually with both SUMO-sfGFP$^{TAG}$ and SUMO-sfGFP$^{TAA}$ in the presence of cognate ncAA (Fig. S3C). We observed that pAzFRS/tRNA$_{CUA}$ only produced detectable fluorescence when paired with SUMO-sfGFP$^{TAG}$ in the presence of pAzF, indicating no detectable suppression of UAA codons by this system (Fig. S3C). Conversely, the Tet3.0/tRNA$_{UUA}$ pair led to detectable (albeit lower) fluorescence when paired with SUMO-sfGFP$^{TAG}$, in addition to the canonical pairing with SUMO-sfGFP$^{TAA}$, indicating that tRNA$_{UUA}$ is capable of decoding both TAA, and to a lesser extent, TAG codons (Fig. S3C). To investigate this further, we suppressed sfGFP$^{TAG}$ protein through the action of the Tet3.0RS/tRNA$_{UUA}$ suppression pair and analyzed it by mass spectrometry (Fig. S3D). As anticipated, we observed a spectrum consistent with homogenous Tet3.0 incorporation (and the corresponding -Met peak), indicating that the Tet3.0 is in fact incorporated by tRNA$_{UUA}$ through decoding of the UAG codon (Fig. S3D). Indeed, others have observed that methanogenic archaeal tRNA$_{UUA}$ is capable of suppressing UAG codons through established wobble-base interactions, so this is not entirely surprising[15]. Importantly, when sfGFP$^{TAG}$ was paired with the Tet3.0/tRNA$_{UUA}$ and the pAzFRS/tRNA$_{CUA}$ systems in the presence of their cognate ncAAs and produced through the action of their simultaneous suppression, we observed homogenous pAzF incorporation at position 150. These results indicate that, like the relative fidelity of pAzFRS, the UAG decoding by tRNA$_{UUA}$ only arises only in the absence of a dominant UAG suppression system. Therefore, we conclude that, under situations where both suppression systems are present and functioning properly, tRNA-Codon orthogonality is maintained and leads to homogenous incorporation of pAzF at UAG codons, and Tet3.0 at UAA codons (Fig. 2B).

***Mass Spectrometry Analysis***

Throughout this study, we have relied on mass spectrometric analysis to confirm ncAA incorporation, evaluate amino acid substitutions, and assess ncAA reactivity. It should be noted here that all mass spectrometry observations were made on sfGFP proteins, since this protein is easily detected during whole-protein electrospray ionization measurement, unlike some other proteins that we have employed, such as SUMO-sfGFP, which we found was not easily detected without cleavage of SUMO domain (data not shown). It should also be noted here that we used two different mutants of sfGFP during our mass spectrometry analysis: $sfGFP^{WT}$ and $sfGFP^{TAG}$ both lack C-terminal "LE" residues preceding the C-terminal 6x His tag, while $sfGFP^{TAA}$ and $sfGFP^{Dual}$ possess these residues, leading to an average mass increase of ~242 Da (see Table S4). All of the major peaks that we have observed can be described by known or predicted modifications, which we will discuss here. Of note here is that all protein samples were desalted at least twice into Type 1 ultrapure water before processing to minimize the formation of salt adduct peaks (notably sodium adducts), which, despite this treatment, are still observable in all spectra[24]. We also find it salient to mention that relative peak intensities are not quantitatively representative in the spectra presented herein, where modifications may affect protein ionization. For a tabulation of all peaks indicated on spectra throughout this work, please refer to Table S6 (salt adduct peaks were excluded from this table due to their ubiquitous nature).

The most notable modification that we have observed throughout our studies was the loss of the N-terminal methionine from sfGFP. This is a common modification observed in proteins heterologously expressed in *E. coli* that leads to the production of a peak with a mass of approximately -131 Da relative to the main parent peak, and is caused by the action of methionyl-aminopeptidase[19] . Moreover, this modification has been observed previously in sfGFP produced by *E. coli* [10–12]. Since this modification does not affect the reactivity of our ncAA-containing sfGFP variants, it can be seen associated with all major species generated through biorthogonal labeling. Another common modification that we see is the reduction of pAzF, which most commonly results in the formation of *para*-aminophenylalanine (pAmF) and a subsequent peak of approximately -26 Da. This modification is present in almost all spectra of pAzF-containing sfGFP, and is likely formed through the reduction of pAzF during *in vivo* expression of these proteins, wherein the cytosol can become a reducing environment due to the presence of biothiols, which, in the case of glutathione, can achieve concentrations of 17 mM[25–29]. Consequently, when exposed to DBCO-$NH_2$ these peaks do not undergo conjugation, indicating that they are indeed unreactive (Fig. S6).

Through a combination of these two modifications, all major peaks present in our spectra can be explained (Table S6). As was previously mentioned, aside from these, the major peaks that we observed were either fully-reacted protein, or unreacted pAzF-containing proteins (Fig. S6A, S6C, S6E). This result is expected based on our quantification of labeling, wherein we routinely saw labeling yields of ~45-85% *in vitro* (Figs. 4A-4B and S5A-S5B). Conversely, spectra wherein Tet3.0-containing proteins were exposed to sTCO-OH were devoid of discernable unreacted products (Figs. S6B, S6D, S6E), which is consistent with the higher labeling yield of the IEDDA reaction that commonly exceeded 90% (Figs. 4A-4B, S5A-S5B). Overall, these results corroborate our previous observations that the SPAAC and IEDDA reactions can be used to effectively and site-specifically label proteins in a one-pot fashion.

***mTagBFP2 Screening and Characterization***

To demonstrate the power of our dual encoding and labeling approach, we sought to perform site-specific *in vivo* protein-protein crosslinking (Fig. 5). To do this, we sought a partner protein that could be easily detected via in-gel fluorescence and could serve as a FRET donor with sfGFP. While this ability was not leveraged as heavily as was designed (i.e. detection of sfGFP alone was sufficient for most of our applications, and ensemble *in vivo* FRET was of lower intensity than anticipated), we decided to utilize the blue fluorescent protein mTagBFP2 due to its excellent spectral overlap with sfGFP and superior photostability[30]. However, this protein has never been the study of genetic code expansion, and so selecting a suitable incorporation site based on previous report was not feasible. As such, we sought to characterize the suitability of this protein for genetic code expansion.

We began by identifying sites tolerant towards pAzF and/or Tet3.0 incorporation. To do so, we selected 6 surface accessible sites located at positions distributed throughout the protein (Fig. S7A) and mutagenized these sites to both TAG and TAA nonsense codons. We paired each $\text{mTagBFP2}^{\text{TAG}}$ and $\text{mTagBFP2}^{\text{TAA}}$ variant with $\text{pEVOL-pAzF}^{\text{TAG}}$ and $\text{pUltraI-Tet3.0}^{\text{TAA}}$, respectively and observed the level of fluorescence produced during expression at the 500 µL scale (Fig. S7B). We observed a wide range of fluorescence in response to pAzF and Tet3.0 incorporation at these sites (Figs. S7B, S7C). Notably, Ochre suppression by the $\text{Tet3.0RS/tRNA}_{\text{UUA}}$ system seemed to result in less fluorescence at each site, consistent with the lower suppression efficiency we have previously observed for this system (Figs. S7B, S2B). In regard to pAzF incorporation, we observed that sites Y14, T105, and N206 produced the highest level of fluorescence, approaching ~50% or greater of the fluorescence of the WT control (Fig. S7B). We next purified many of these protein variants and estimated the yield in mg/L, and found that this yield corresponded well with the observed suppression efficiency and ranged from ~30-80 mg/L for pAzF-containing variants (Table S5). We next investigated if pAzF incorporation adversely affected mTagBFP2 fluorescence. To do so, we isolated the four highest-expressing $\text{mTagBFP2}^{\text{pAzF}}$ variants and measured their fluorescence spectra (Fig. S7D). What we observed is that, when normalized for concentration, none of the variants we tested exhibited any spectral shifts or decreases in fluorescence intensity (Fig. S7D). Based on these results, we conclude that pAzF incorporation into mTagBFP2 does not adversely affect the spectral properties of the protein. Since site T105 could be robustly produced, and this position occupies a site comparable to position N150 in sfGFP (i.e. located on the side of the barrel), and exhibits high yield, we elected to continue with this variant for downstream applications.

***sfGFP/mTagBFP2 and sfGFP-mTagBFP2 Expression and Characterization***

To showcase the some of the myriad abilities that our dual encoding and labeling method affords, we elected to express $\text{sfGFP}^{\text{Tet3.0}}$ and $\text{mTagBFP2}^{\text{pAzF}}$ either as two independent proteins (i.e. for intermolecular crosslinking), or as a fusion protein (i.e. intramolecular stapling). However, in these instances, there are two different orientations within which one could orient these two different genes, and as such we sought to explore which orientations were most optimal for protein production.

In the case of intermolecular crosslinking, we required the production of $\text{sfGFP}^{\text{Tet3.0}}$ and $\text{mTagBFP2}^{\text{pAzF}}$ as separate proteins. To do so, we elected to express both of these proteins from a pETduet vector, which contains two multiple cloning sites, each proceeding a T7 promotor. In this instance, we reasoned that the crosslinking reaction would occur optimally if the $\text{sfGFP}^{\text{Tet3.0}}$, $\text{mTagBFP2}^{\text{pAzF}}$, and heterobifunctional crosslinker were present in a 1:1:1 ratio, and as such, controlling the ratio of $\text{sfGFP}^{\text{Tet3.0}}$ to $\text{mTagBFP2}^{\text{pAzF}}$ to achieve the most balanced ratio is desirable. Since this construct lacks a terminator between the two promotor regions, we hypothesized that this configuration would produce polar effects that may affect how each protein is translated depending on whether it is positioned upstream or downstream of the other. In this situation, the open reading frame that is positioned downstream would be present on two different mRNA transcripts as either the downstream open reading frame of a dicistronic transcript containing both genes, and as a monocistronic transcript with only a single gene (Fig. S8A). As such, positioning sfGFP$^{TAA}$ in the upstream or downstream position, for example, may affect the suppression efficiency, and therefore the yield of sfGFP$^{Tet3.0}$ (Fig. S8A). To see which orientation produced the most balanced ratio of sfGFP$^{Tet3.0}$ to mTagBFP2$^{pAzF}$ we generated two different pETduet constructs that positioned the sfGFP$^{TAA}$ either upstream or downstream of mTagBFP2$^{TAG}$, with both genes possessing a C-terminal 6xHis tag sequence and observed the fluorescence signal produced in each scenario. As can be seen in Fig. S8B, when sfGFP$^{TAA}$ was positioned downstream we observed a greater fluorescent output. We next purified the proteins resulting from both orientations and estimated the yield of the protein through the absorption of each fluorophore (Table S5). We observed that mTagBFP2$^{pAzF}$ was present in greater abundance, likely due to the superior suppression efficiency of the pAzFRS/tRNA$_{CUA}$ suppression system. Moreover, we found that the orientation that positioned sfGFP$^{TAA}$ downstream of mTagBFP2$^{TAG}$ resulting in the most balanced ratio of sfGFP$^{Tet3.0}$:mTagBFP2$^{pAzF}$ at ~1:6.7, and yields of ~10 and 67 mg/L (Fig. S8B, Table S5). To demonstrate that the produced proteins were reactive, we exposed the resulting protein mixtures from each orientation to sTCO-PEG$_{5000}$ and DBCO-TAMRA, and observed a mobility-shifted product in lower abundance and a fluorescent band (Fig. S8C). These results suggest that the positioning of the gene can affect the yield of the resulting proteins, that positioning sfGFP$^{TAA}$ in the downstream position produces a more balanced ratio with respect to mTagBFP2$^{pAzF}$, and that the resulting proteins are reactive.

To achieve intramolecular protein stapling, we sought to leverage positions that we had previously demonstrated are reactive by pAzF and Tet3.0. As such, we sought to produce an sfGFP$^{Tet3.0}$-mTagBFP2$^{pAzF}$ fusion protein that contains Tet3.0 and pAzF in positions that we had previously confirmed were reactive. While designing constructs to fuse these two proteins together, we also elected to include a tobacco etch virus protease (TEV) cut site located within the linker region such that the two domains could be proteolytically separated. In the interest of optimizing yields for *in vivo* analysis, we sought to explore which relative orientation would produce the most protein, and also explore which orientation was most reactive. To do so, we produced two different sfGFP-mTagBFP constructs, one wherein sfGFP proceeds and mTagBFP2, and one wherein sfGFP precedes mTagBFP2 (Fig. S10A). In each instance, sfGFP possessed a TAA mutation at the site analogous to site 150, and mTagBFP2 possessed a TAG mutation at the site analogous to 105. We found that the orientation that positioned sfGFP$^{TAA}$ preceding mTagBFP2$^{pAZF}$ yielded more protein than the alternative configuration (Fig. S10B), consistent with our earlier observation that positioning UAA upstream of UAG produced more SUMO-sfGFP protein (Figs. S8B, 2A). As was mentioned in the main text, we found no marked difference between stapling efficiency between these two orientations (Fig. 6B, S11). We did note, however, that after cleavage by TEV that two distinct bands were detectable in the sfGFP channel: a slow migrating product of similar mobility to the uncleaved product, and a lower, more mobile band (Fig. 6B, lanes 5 and 10). As mentioned in the main text, the slowly migrating band is presumed to be cleaved, stapled sfGFP-mTagBFP2 protein. We have previously observed that the Tet3.0-sTCO reaction is high yielding (Figs. 3, 4), and it is known that the mobility of proteins may be altered substantially and unpredictably by post-translational modifications[31]. For these reasons we surmise that this faster migrating band is cleaved sfGFP$^{Tet3.0}$ that has reacted with the sTCO-PEG$_4$-DBCO, but has not subsequently reacted with mTagBFP2$^{pAzF}$. Upon close inspection we observed a faint band of similar mobility to cleaved sfGFP$^{WT}$ (Fig. 6B, lanes 1 and 6), which is likely cleaved, but unreacted sfGFP$^{Tet3.0}$. We see this product form analogously during *in vivo* stapling in a time- and crosslinker-dependent manner wherein it reaches maximal intensity after ~15 minutes, further supporting the notion of this product being a by-product of the stapling reaction (Fig. 6C). Moreover, we observed the time- and crosslinker-dependent formation of a similar product during *in vivo* crosslinking (Fig. 5D); however, due to differences in sequence (e.g. presence of 6xHis-tag, and absence of TEV cleavage scar), and the heterobifunctional linker composition (e.g. crosslinking by sTCO-DBCO as opposed to stapling with sTCO-PEG$_4$-DBCO) the mobility of this product differs. We conclude that the orientation that positions the TAA codon upstream of the TAG codon optimizes protein yield, and that these two different orientations produce similar levels of stapled product in response to the addition of heterobifunctional crosslinker.

### Supplementary Schemes

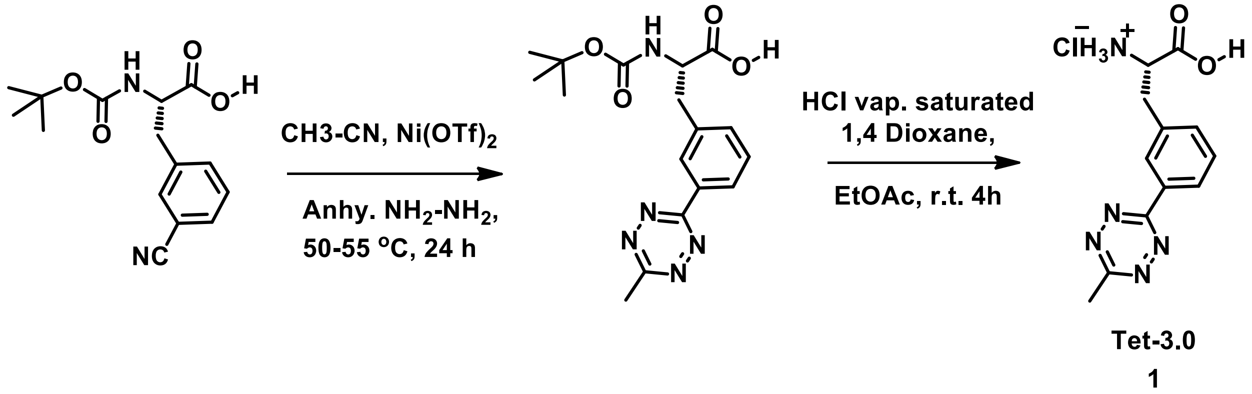

***Scheme S1.*** Synthesis of *meta*-substituted s-tetrazine derivative of phenylalanine (Tet-3.0).

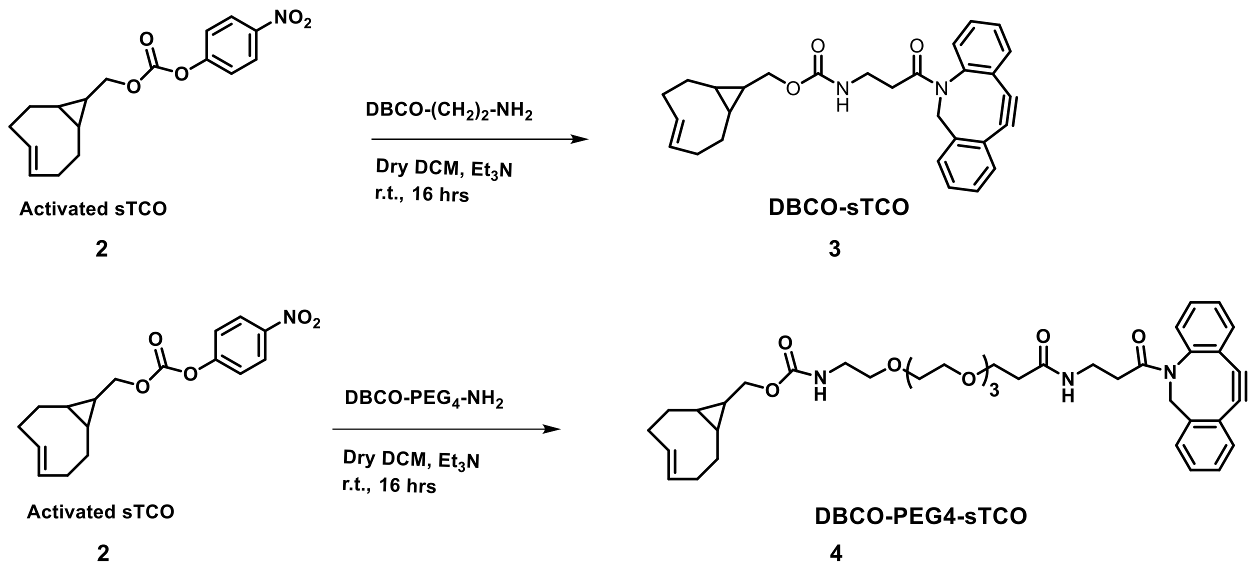

***Scheme S2***. Synthesis of DBCO-sTCO and DBCO-$PEG_4$-sTCO.

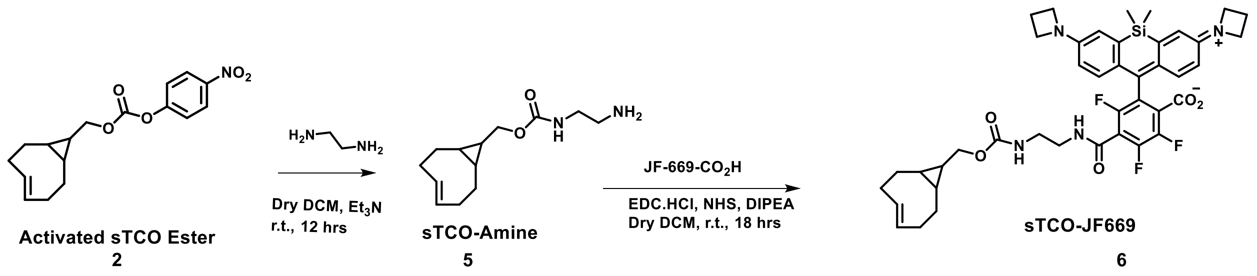

***Scheme S3.*** Synthesis of sTCO-JF669.

### Supplementary Figures

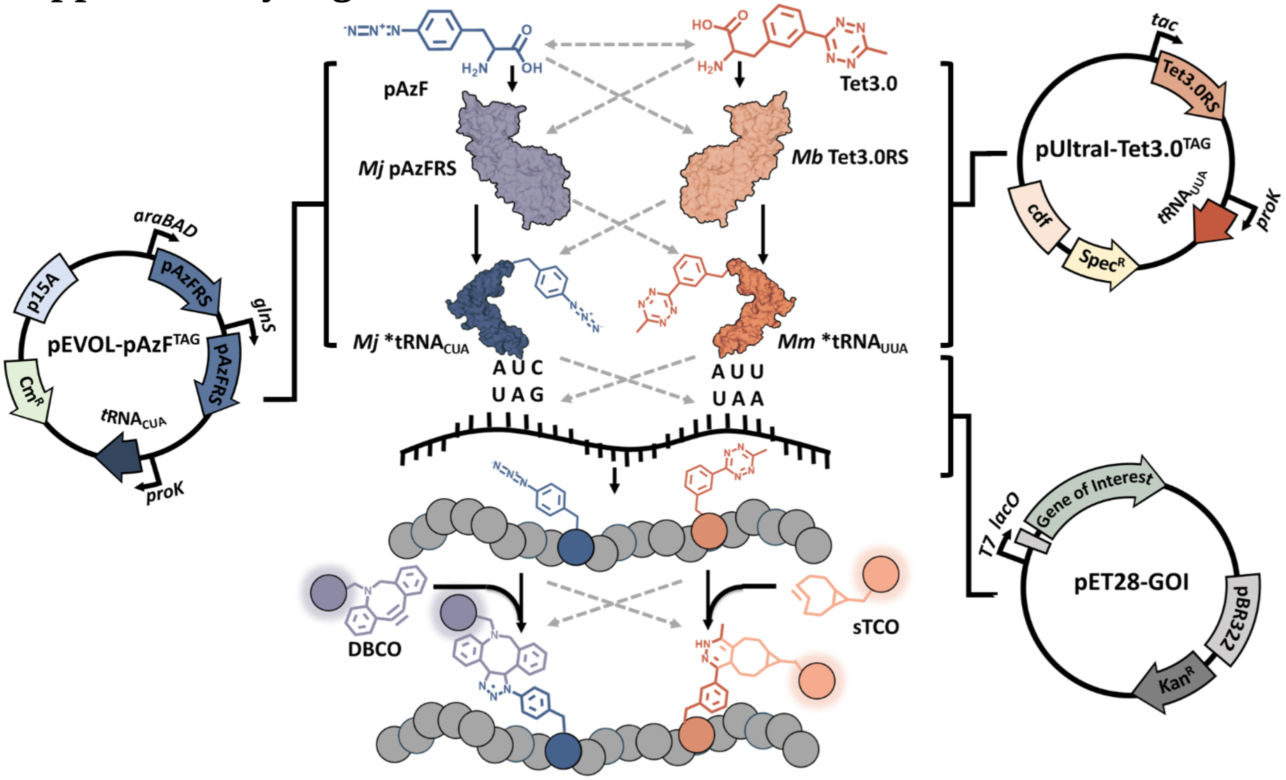

***Figure S1***. Overview of dual ncAA encoding and labeling subsystems used in this study. Above is a schematic illustrating genetic code expansion during dual-ncAA suppression using our system, with salient features depicted. Here, pAzF is recognized by an engineered *Methanocaldococcus janaschii* (*Mj*) pAzFRS and is charged onto *Mj*tRNA$_{CUA}$, forming *Mj**tRNA$_{CUA}$, which subsequently suppresses UAG codons to drive pAzF incorporation into a nascent protein during translation. Conversely, Tet3.0 is recognized by an engineered *Methanoscarcina barkeri* (*Mb*) Tet3.0RS and is charged onto *Methanosarcina mazei* (*Mm*) *Mm*tRNA$_{UUA}$, forming *Mm**tRNA$_{UUA}$, which subsequently suppresses UAA codons to drive incorporation of Tet3.0 into a nascent protein, in this case, downstream of pAzF. Solid black arrows indicate the desired flow of substrates, whereas dashed, gray arrows indicate undesirable, non-orthogonal substrate flux. Key points of critical orthogonality are denoted as dashed lines: at the dual encoding level between amino acids (top, lateral dashed gray arrows), at the ncAA-aaRS level (top crossed arrows), the aaRS-tRNA level (upper middle crossed arrows), at the tRNA-codon level (lower middle crossed arrows), or at the dual labeling level for strain-promoted azide-alkyne coupling reaction (SPAAC) between pAzF and DBCO and the inverse electron-demand Diels-Alder reaction (IEDDA) between Tet3.0 and sTCO (bottom crossed arrows). The plasmid maps at right and left depict the three plasmids used in our final, optimized configuration, including relevant features such as resistance, origin, tRNA and aaRS, and promoters.

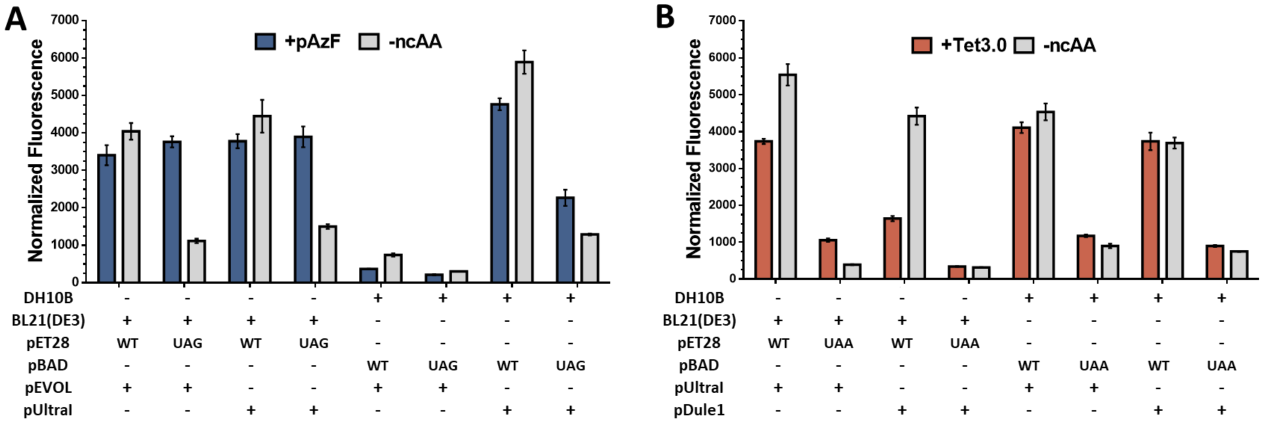

***Figure S2.*** Optimal vector and strain combinations for UAG and UAA suppression subsystems. Normalized SUMO-sfGFP fluorescence for (A) UAG suppression subsystems in the presence (blue) and absence (gray) of pAzF, and for (B) UAA suppression subsystems in the presence (orange) and absence (gray) of Tet3.0 after 24 hours of expression. UAG or UAA sites are located at position 35 in the SUMO domain.

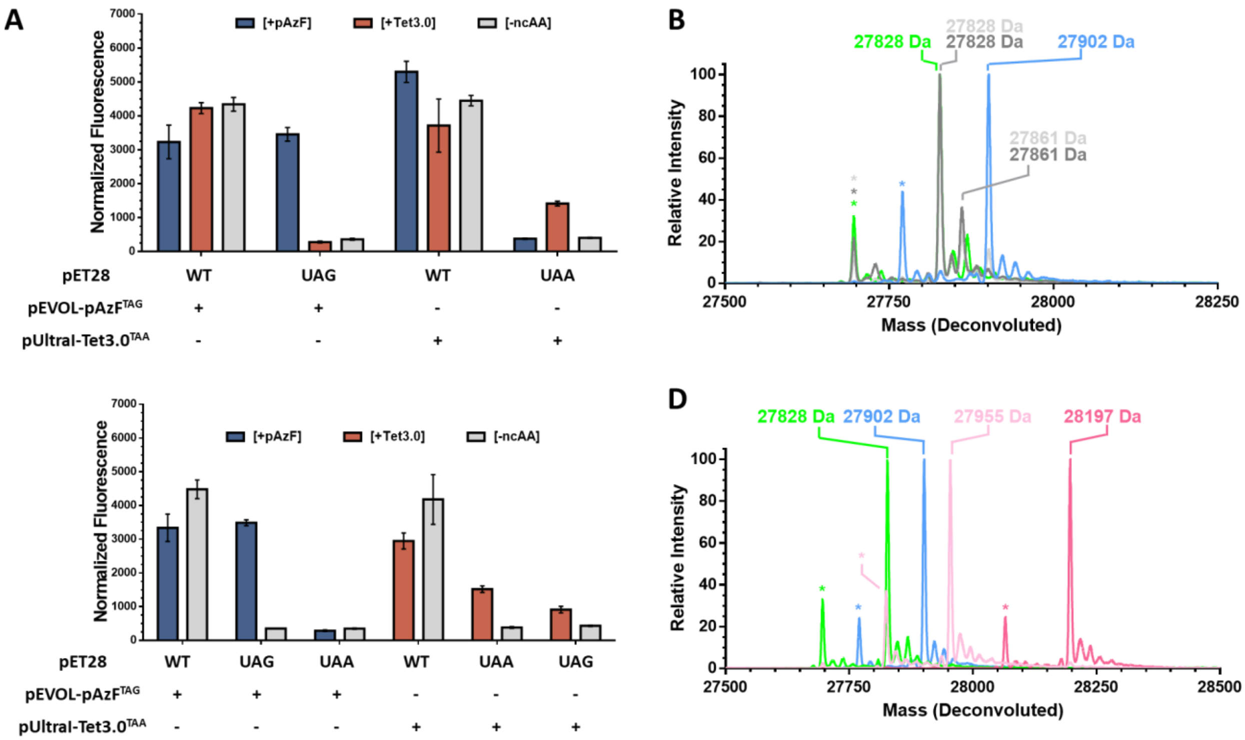

***Figure S3***. Assessment of orthogonality between genetic code expansion components in *E. coli* BL21(DE3) strains expressing SUMO-sfGFP variants. (A) Assessment of ncAA-aaRS orthogonality. Normalized SUMO-sfGFP fluorescence from expression of reporters with each cognate suppression subsystem in the presence of pAzF (blue), Tet3.0 (orange), or neither ncAA (gray). (B) Overlaid ESI mass spectra of sfGFP$^{TAG}$ suppressed by pAzFRS/tRNA$_{CUA}$ in the presence of pAzF (blue), Tet3.0 (light gray), or absence of ncAA (dark gray). sfGFP$^{WT}$ is included as a reference (green). (C) Assessment of tRNA-codon orthogonality. Normalized fluorescence from expression of SUMO-sfGFP$^{TAG}$ and SUMO-sfGFP$^{TAA}$ reporter in combination with each suppression subsystem in the presence or absence of its cognate ncAA (colors analogous to panel A). (D) Overlaid ESI mass spectra of sfGFP$^{TAG}$ suppressed either by pUltraI-Tet3.0$^{TAA}$ individually in the presence of Tet3.0 (light pink), or in combination with pEVOL-pAzF$^{TAG}$ in the presence of both ncAAs (blue). sfGFP$^{WT}$ (green) and sfGFP$^{Tet3.0}$ (pink) are included as references. Peaks corresponding to the loss of N-terminal methionine are indicated by "*", while peaks corresponding to pAzF reduction are indicated by "†". Mass measurement error is ± 1 Da, see Table S6 for expected masses.

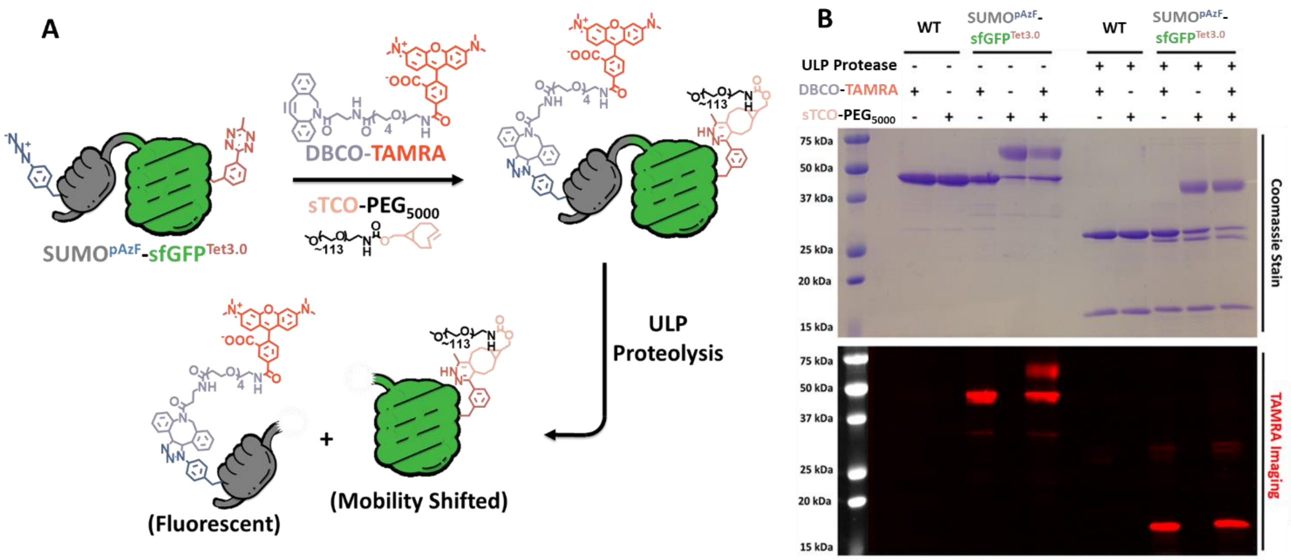

***Figure S4.*** *In vitro* dual labeling of dual encoded SUMO-sfGFP and ULP1 digestion. (A) Schematic depicting SUMO$^{pAzF}$-sfGFPT$^{et3.0}$ undergoing dual labeling by DBCO-TAMRA and sTCO-PEG$_{5000}$ followed by proteolytic cleavage by ULP1 protease. (B) SDS-PAGE of SUMO-sfGFP$^{WT}$ and SUMO$^{pAzF}$-sfGFP$^{Tet3.0}$ exposed to DBCO-TAMRA and/or sTCO-PEG$_{5000}$ with or without ULP1-mediated protein cleavage. The contents of each lane are indicated above each gel image. The upper inset shows Coomassie staining of the gel, while the lower inset shows fluorescent gel imaging in the TAMRA channel.

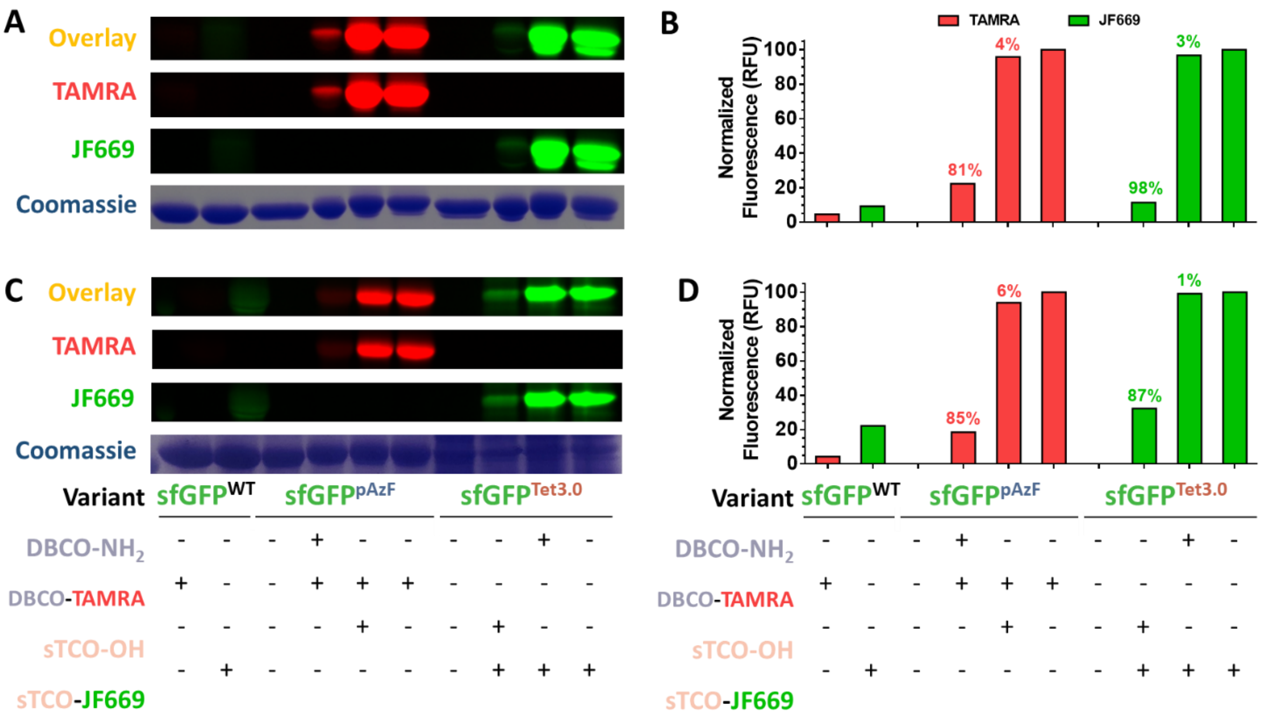

***Figure S5***. Assessment of IEDDA and SPAAC mutual orthogonality. (A) In-gel fluorescence of SDS-PAGE of *in vitro* reactions of sfGFP variants exposed to DBCO-$NH_2$ or sTCO-OH for 24 hours and 15 minutes, respectively, prior to quenching with DBCO-TAMRA or sTCO-JF669. The top image is an overlay of both TAMRA and JF669 fluorescent signals followed by the individual channels and Coomassie staining (lower). (B) Densitometry quantification of the fluorescent channels from panel A (green bars indicate quantification in the JF669 channel, while red bars indicate quantification in the TAMRA channel) with the calculated labeling yield indicated above corresponding bars. (C) *In vivo* labeling analysis performed and presented analogously to panel A. (D) Densitometry quantification of fluorescent gel channels in panel C, processed and presented analogously to panel B. The contents of each lane are indicated below each gel.

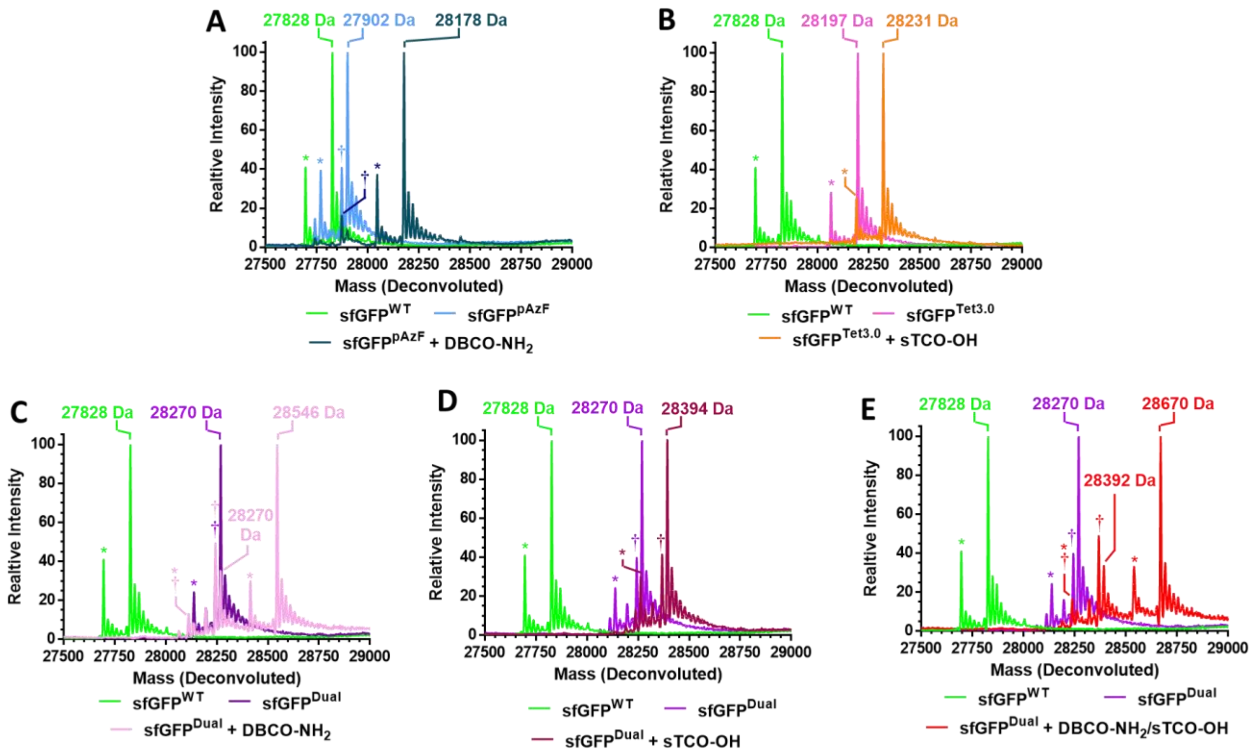

***Figure S6***. ESI Mass spectrometry of sfGFP variants labeled *in vitro*. Purified sfGFP containing pAzF were exposed to DBCO-$NH_2$ for 24 hours, while sfGFP containing Tet3.0 were exposed to sTCO-OH for 15 minutes in mass spectrometry-grade water and were desalted to remove excess labeling reagents prior to analysis. (A) Overlaid mass spectra of sfGFP$^{pAzF}$ before (light blue) and after DBCO-$NH_2$ addition (dark blue). (B) Overlaid mass spectra of sfGFP$^{Tet3.0}$ before (pink) and after sTCO-OH addition (orange). (C) Overlaid mass spectra of sfGFP$^{Dual}$ after DBCO-$NH_2$ addition (lilac trace). (D) Overlaid mass spectra of sfGFP$^{Dual}$ after sTCO-OH addition (mauve). (E) Overlaid mass spectra of sfGFP$^{Dual}$ after DBCO-$NH_2$ and sTCO-OH addition (Red trace). sfGFP$^{WT}$ (green; all panels) and unreacted sfGFP$^{Dual}$ (purple; panels C-E) are included as references. Peaks corresponding to the loss of N-terminal methionine are indicated by "*", while peaks corresponding to pAzF reduction are indicated by "†". Mass measurement error is ± 1 Da, see Table S6 for expected masses.

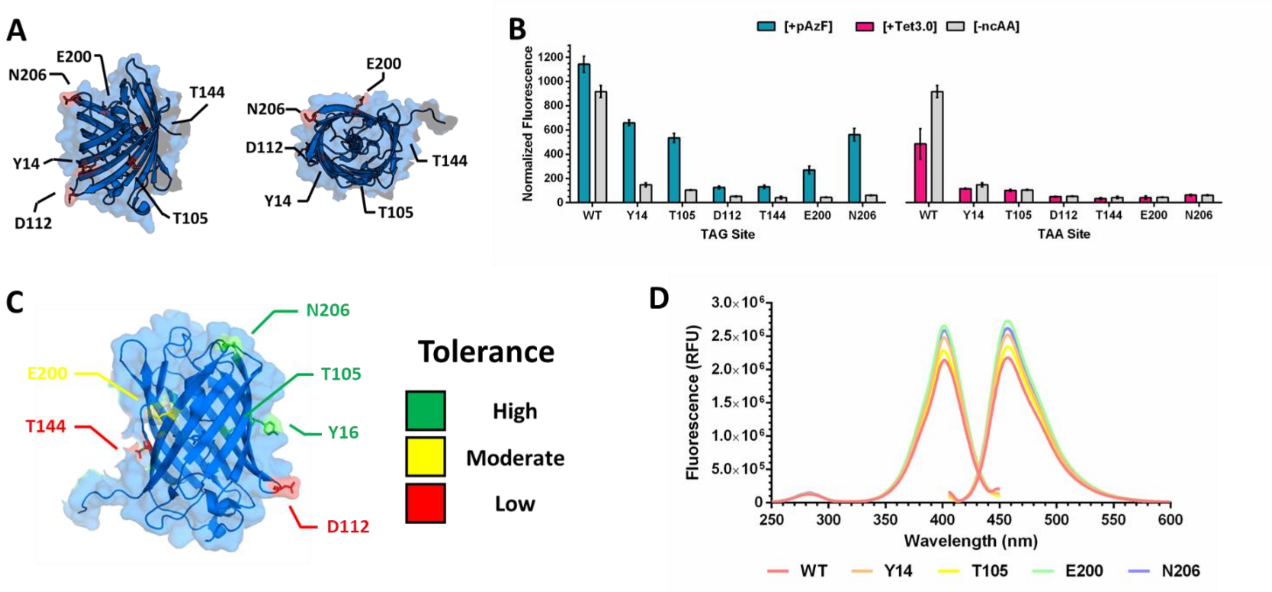

***Figure S7***. Characterization of mTagBFP2 as a fluorescent reporter for genetic code expansion. (A) Crystal structure of mTagBFP (PDB code 3M24) with the six residues selected for pAzF and Tet3.0 encoding highlighted in pink and labeled. (B) Normalized fluorescence of mTagBFP2 variants resulting from UAG suppression by pEVOL-pAzF$^{TAG}$ in the presence of pAzF (blue bars) or UAA suppression pUltraI-Tet3.0$^{TAA}$ in the presence of Tet3.0 (orange bars) or absence of ncAA (gray bars). (C) mTagBFP2 structure with six residues color coded to reflect pAzF encoding tolerance. As indicated at top right: green corresponds to high tolerance (>50% suppression efficiency), yellow corresponds to moderate tolerance (50% - 10% suppression efficiency), while red corresponds to low tolerance (<10% suppression efficiency). (D) Fluorescence spectra of five different mTagBFP2$^{pAzF}$ variants normalized by concentration showing minimal fluorescent change from ncAA encoding. Excitation spectra were scanned from 300 nm to 450 nm with emission detection fixed at 454 nm, while emission spectra were scanned from 405 nm to 600 nm with excitation fixed at 399 nm.

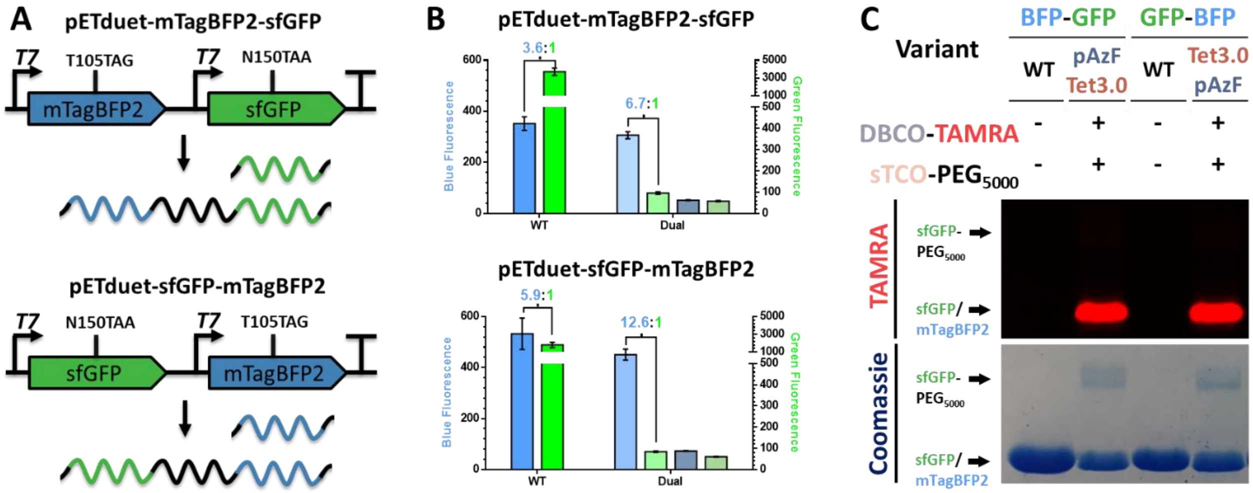

***Figure S8***. Characterization of sfGFP$^{Tet3.0}$ and mTagBFP$^{pAzF}$ co-expression and reactivity. (A) Graphic depicting the orientation of two mTagBFP2+sfGFP open reading frames on pETduet vectors and the resulting mRNA. (B) Normalized mTagBFP2 (blue bars; left axis) and sfGFP (green bars; right axis) fluorescence resulting from dual suppression of mTagBFP2$^{TAG}$+sfGFP$^{TAA}$ (top panel) and sfGFP$^{TAA}$+mTagBFP2$^{TAG}$ in the presence (light blue/green bars) and absence (muted blue/green bars) of ncAA. Maximal expression yield is measured from fluorescence of expressed WT equivalents (bright blue/green bars). Indicated above are the ratio of mTagBFP2:sfGFP as determined from purified proteins. (C) SDS-PAGE analysis demonstrating the reactivity of purified sfGFP$^{Tet3.0}$/mTagBFP2$^{pAzF}$ from both orientations after exposure to DBCO-TAMRA and sTCO-PEG$_{5000}$. Upper gel image corresponds to in-gel fluorescence observed under a TAMRA channel, while the lower image corresponds to the same gel visualized by Coomassie staining.

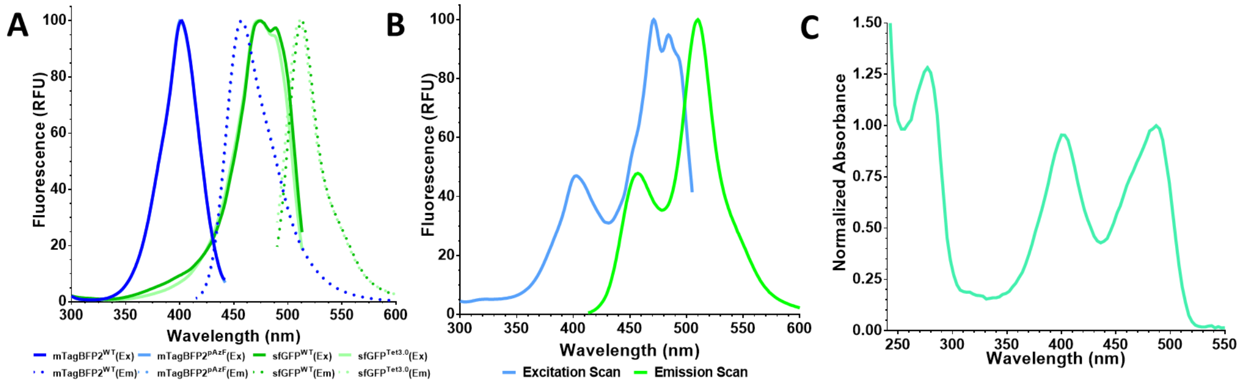

***Figure S9***. Fluorescence and absorbance properties of crosslinked sfGFP$^{Tet3.0}$-sTCO-DBCO-mTagBFP2$^{pAzF}$. (A) Excitation (solid lines) and emission spectra (dotted lines) of purified mTagBFP2$^{WT}$ (dark blue), mTagBFP2$^{pAzF}$ (light blue), sfGFP$^{WT}$ (dark green), and sfGFP$^{Tet3.0}$ (light green). Excitation spectra for mTagBFP2 were scanned from 300-442 nm, with fluorescence observed at 454 nm, while emission spectra were scanned from 415-600 nm, with excitation at 399 nm. Excitation spectra for sfGFP were scanned from 300-513 nm, with fluorescence observed at 513 nm, while emission spectra were scanned from 490-600 nm, with excitation at 485 nm. (B) Excitation (blue line) and emission spectra (green line) of isolated, crosslinked sfGFP$^{Tet3.0}$-sTCO-DBCO-mTagBFP$^{pAzF}$. Excitation spectrum was determined by scanning from 300-505 nm, with fluorescence observed at 513 nm; emission spectrum was determined by scanning from 404-600 nm, with excitation at 399 nm. (C) Absorbance spectra of isolated sfGFP$^{Tet3.0}$-sTCO-DBCO-mTagBFP$^{pAzF}$ showing strong absorbance at both 399 nm and 485 nm indicating crosslinking of sfGFP and mTagBFP2.

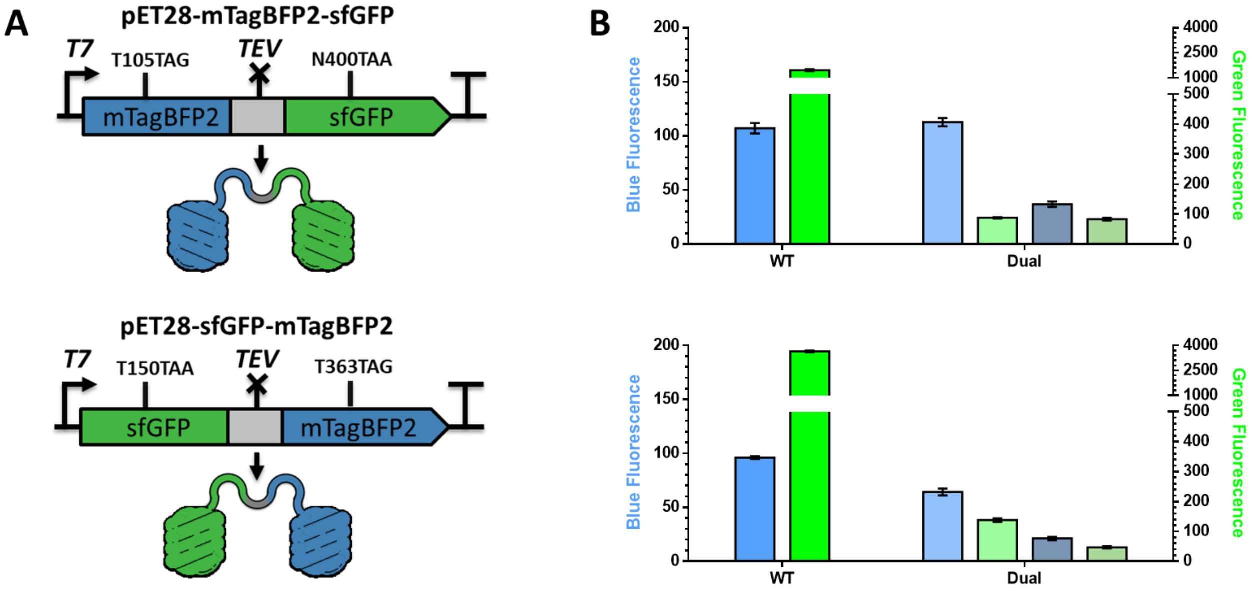

***Figure S10***. Expression characterization of sfGFP-mTagBFP2 fusion proteins**.** (A) Graphic depicting the orientation of the two different sfGFP-mTagBFP2 fusion proteins used in this study with salient features, as well as the resulting product produced shown. (B) Normalized mTagBFP2 (blue bars; left axis) and sfGFP (green bars; right axis) fluorescence resulting from dual suppression of mTagBFP2$^{TAG}$-sfGFP$^{TAA}$ (top panel) and sfGFP$^{TAA}$-mTagBFP2$^{TAG}$ (bottom panel) in the presence (light blue/green bars) and absence (muted blue/green bars) of ncAAs. Maximal expression yield is measured from fluorescence of expressed WT equivalents (bright blue/green bars).

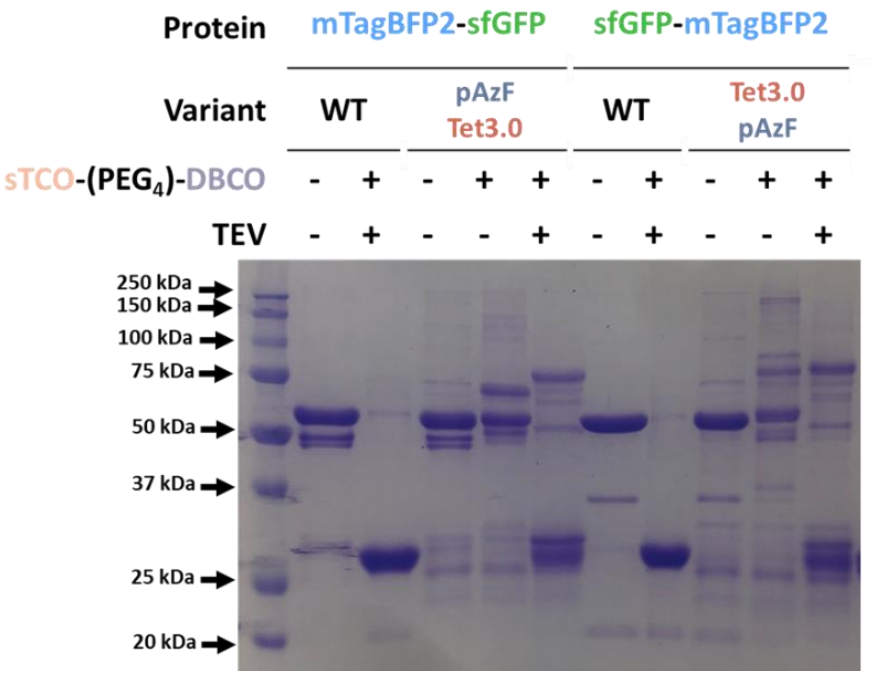

***Figure S11***. Assessment of *in vitro* intramolecular stapling on dual encoded reporters. SDS-PAGE of purified sfGFP$^{Tet3.0}$-mTagBFP2$^{pAzF}$ and mTagBFP2$^{pAzF}$-sfGFP$^{Tet3.0}$ exposed to sTCO-PEG$_4$-DBCO were analyzed after cleavage by TEV protease. Coomassie staining shows distinct mass changes associated with efficient intramolecular after TEV cleavage.

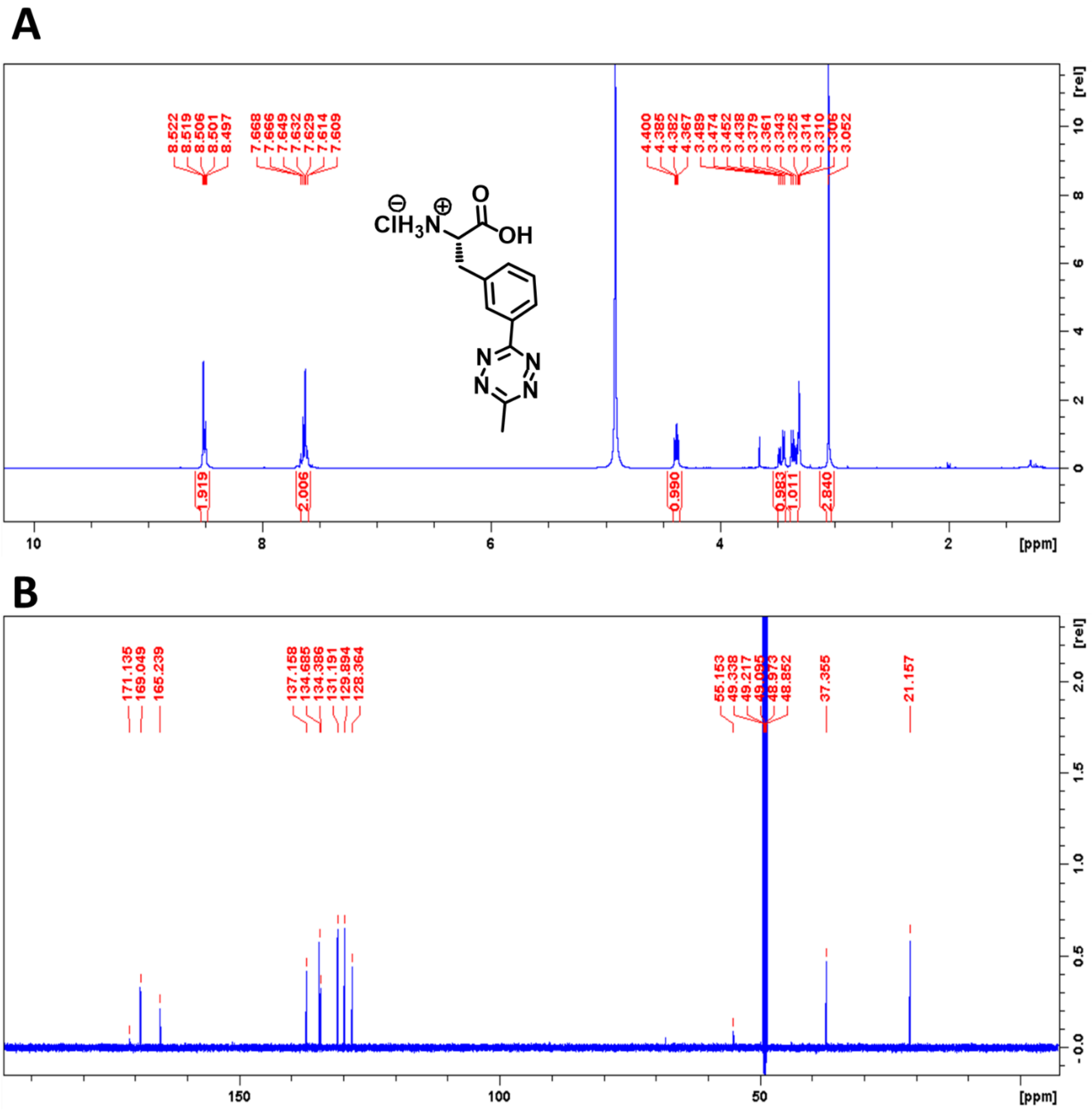

***Figure S12.*** NMR spectra of Tet3.0 (**1**). (A) $^{1}H$ spectra, (B) $^{13}C$ spectra.

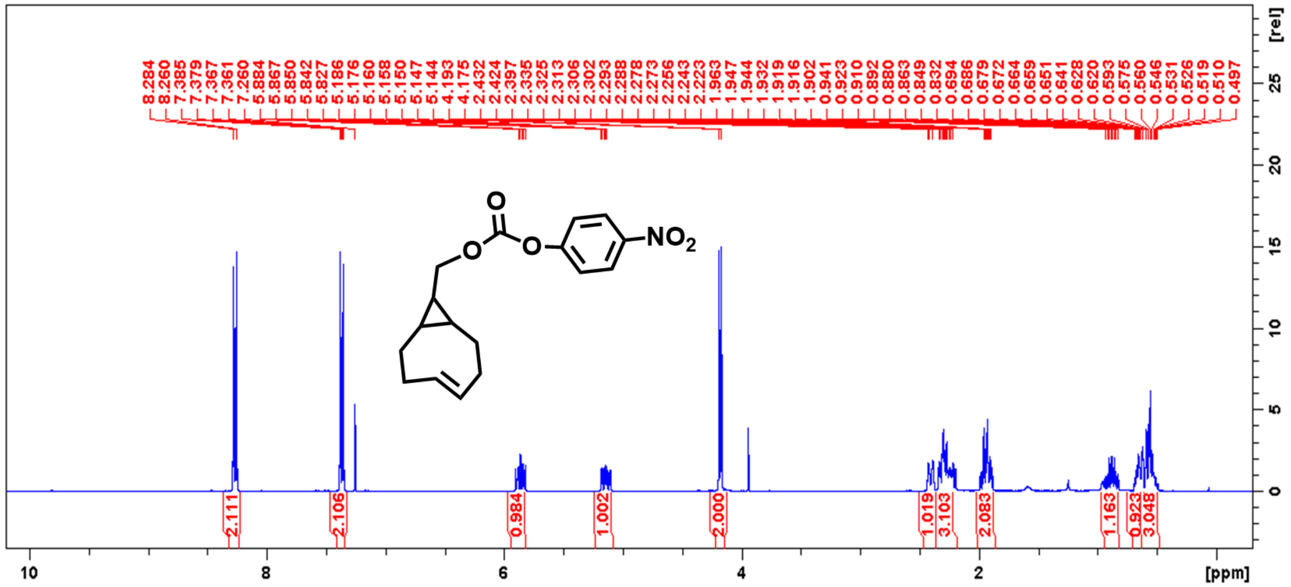

***Figure S13***. [1]H NMR spectra of activated ester of sTCO (**2**).

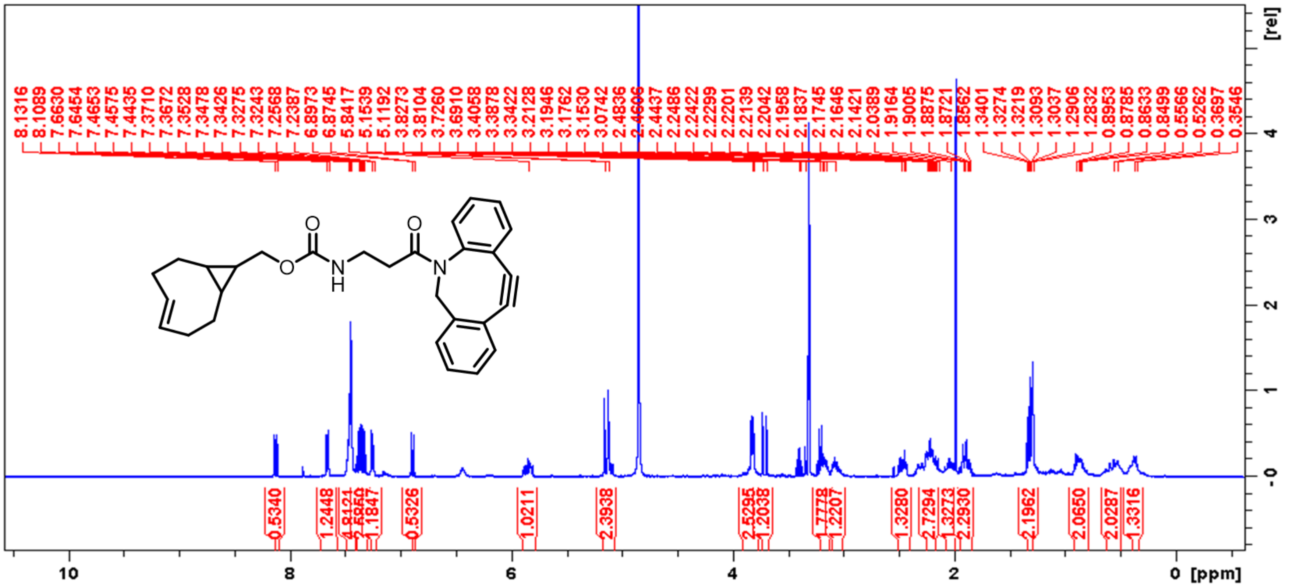

***Figure S14.*** [1]H NMR spectra of DBCO-sTCO (**3**).

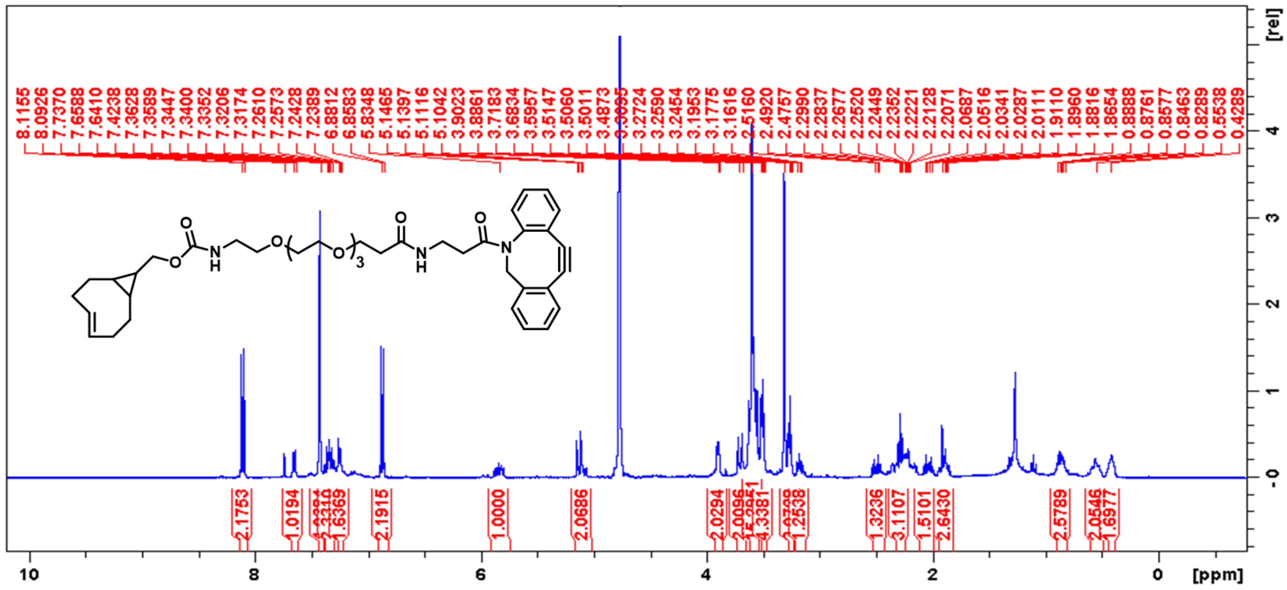

***Figure S15.*** $^1$H NMR spectra of DBCO-PEG$_4$-sTCO (**4**).

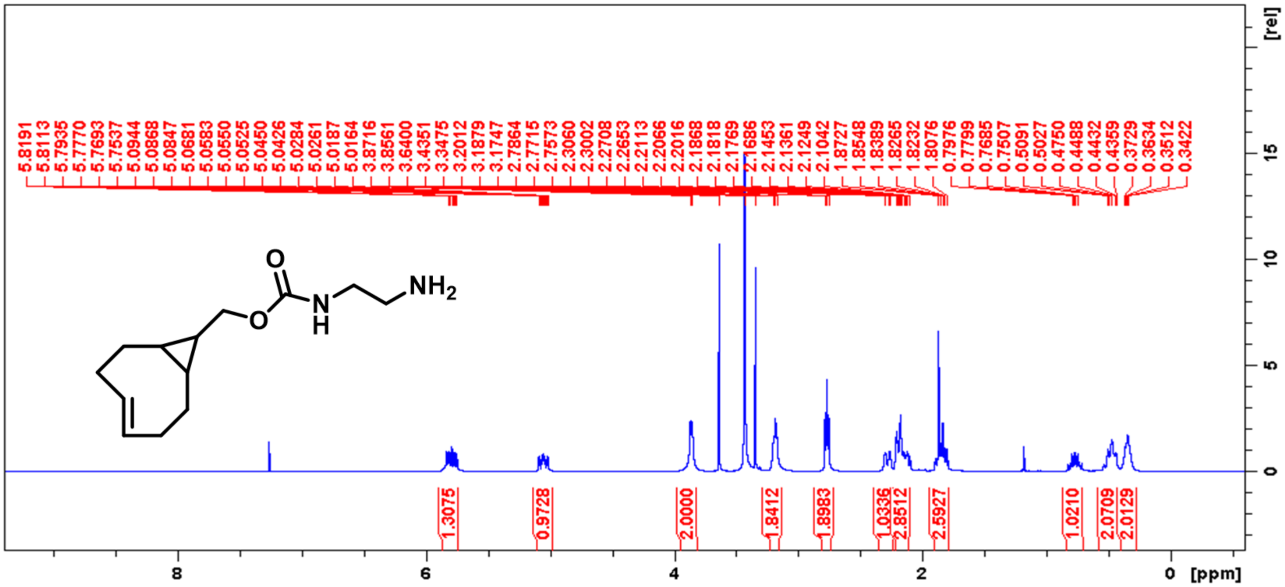

***Figure S16.*** $^{1}$H NMR spectra of sTCO-Amine (**5**).

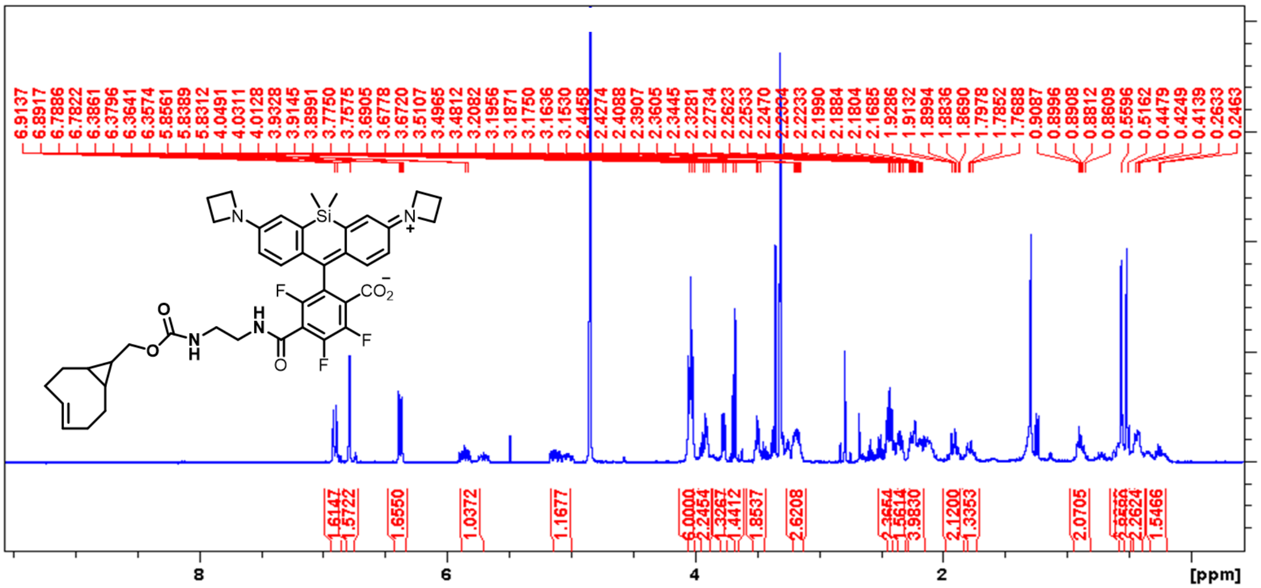

***Figure S17.*** $^{1}$H NMR spectra of sTCO-JF669 (**6**)

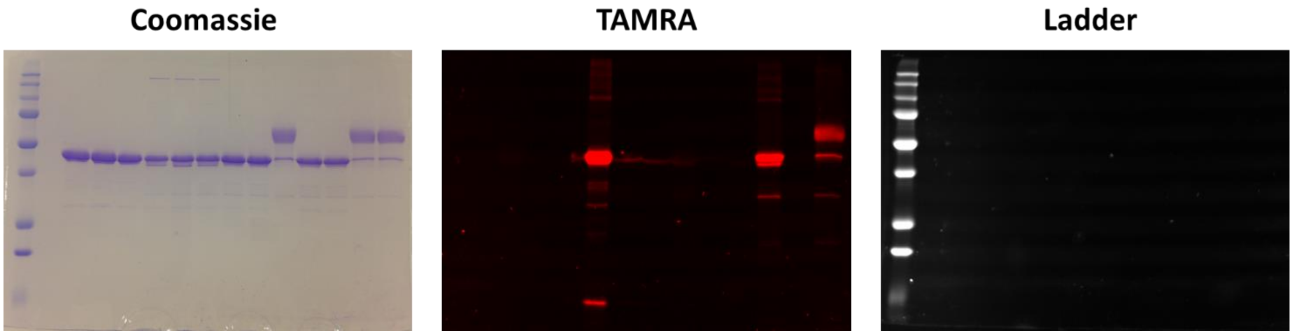

***Figure S18***. Unedited gel images for Figure 3B, including Coomassie-stained (left), TAMRA (middle) and fluorescent ladder (left) channels.

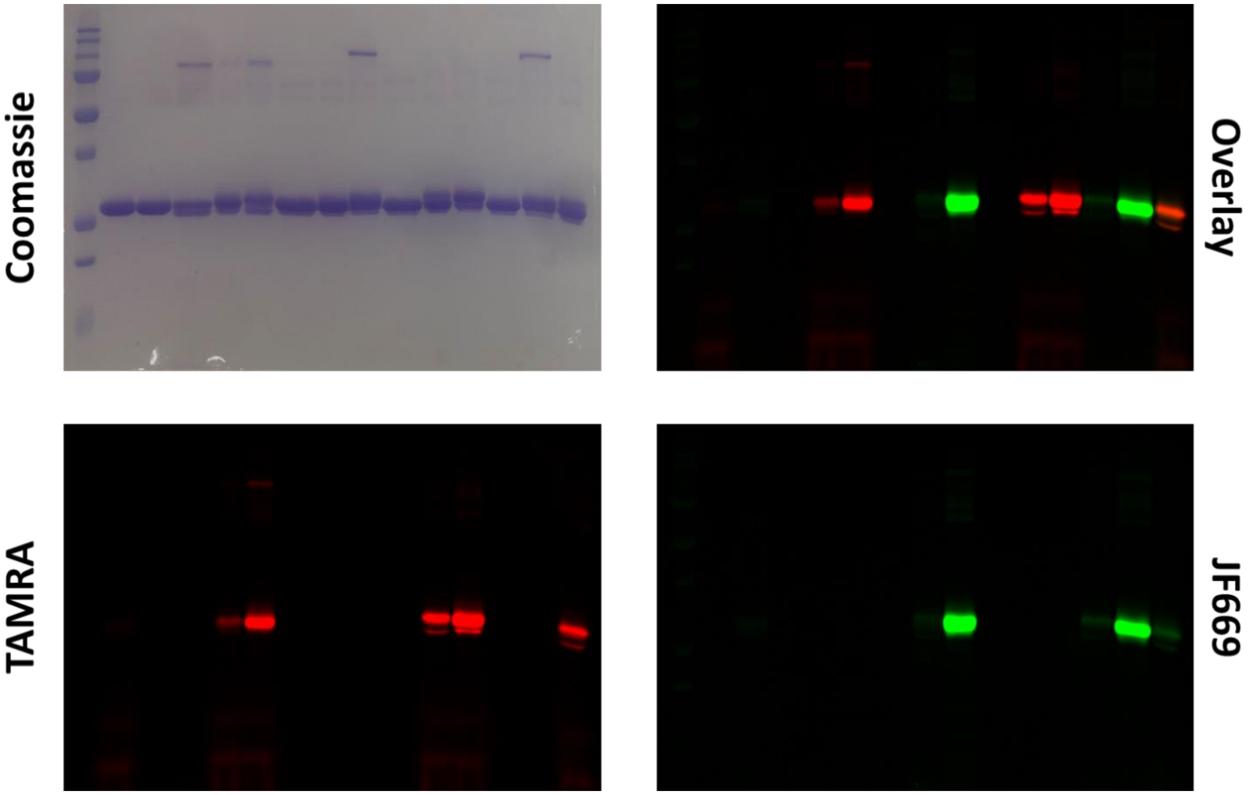

***Figure S19***. Unedited gel images for Figure 4A, including Coomassie-stained (top left), TAMRA (bottom left), JF669 (bottom right) channels, and their overlay (top right).

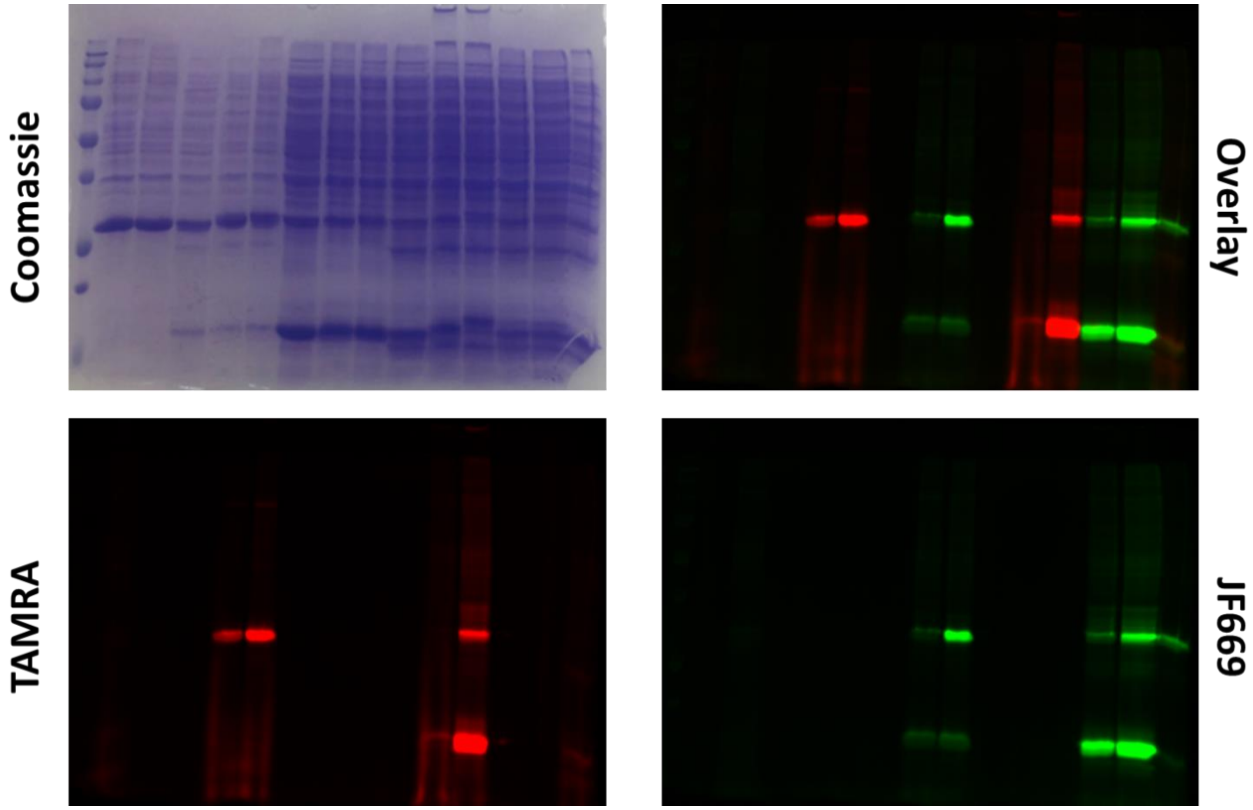

***Figure S20***. Unedited gel images for Figure 4C, including Coomassie-stained (top left), TAMRA (bottom left), JF669 (bottom right) channels, and their overlay (top right). Note, lower, high-abundance bands correspond to truncation product, which is a common byproduct of GCE and can experience specific (and non-specific) labeling due to their abundance and distribution.

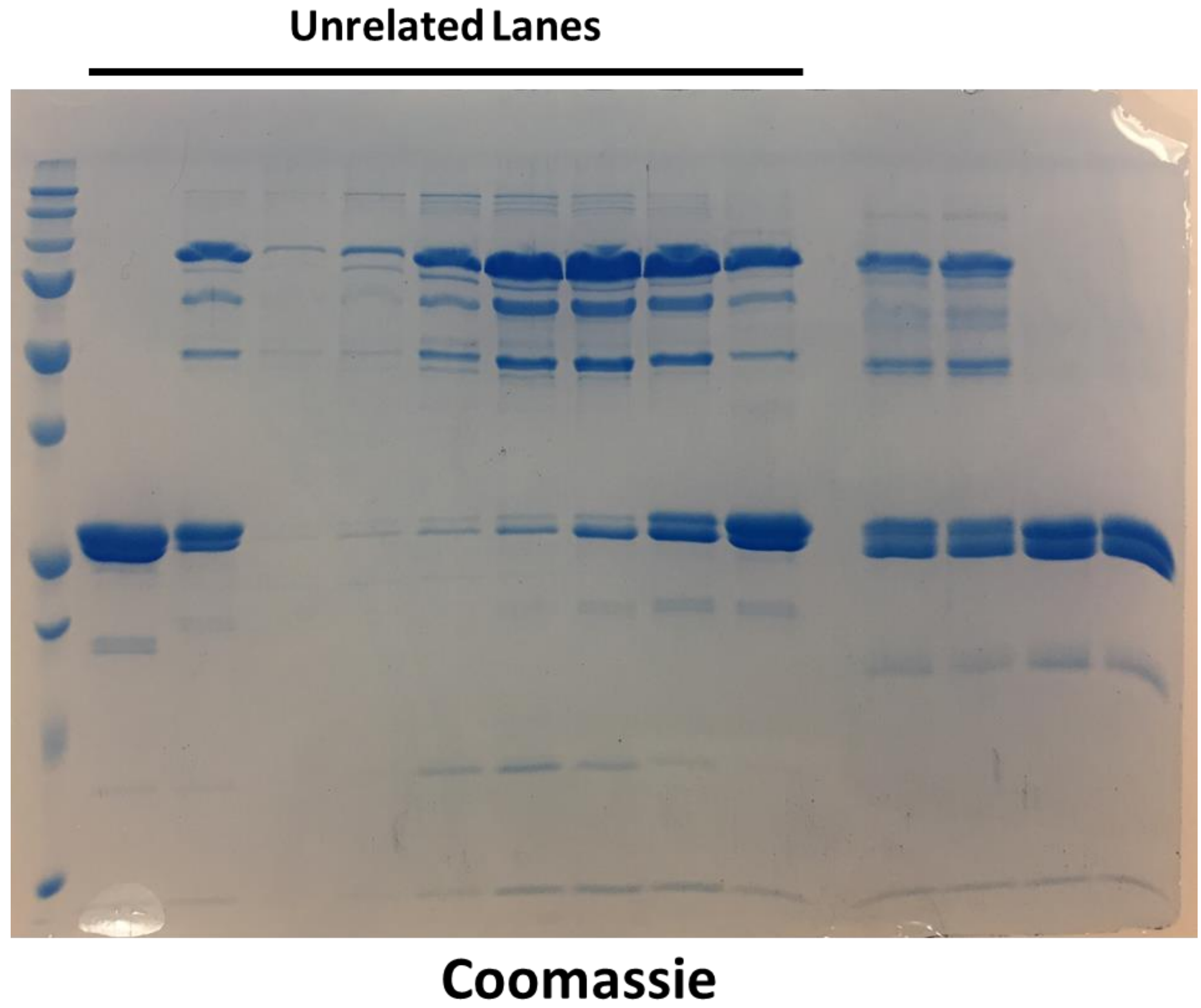

***Figure S21***. Unedited gel image for Figure 5B. Top bar indicates unrelated lanes

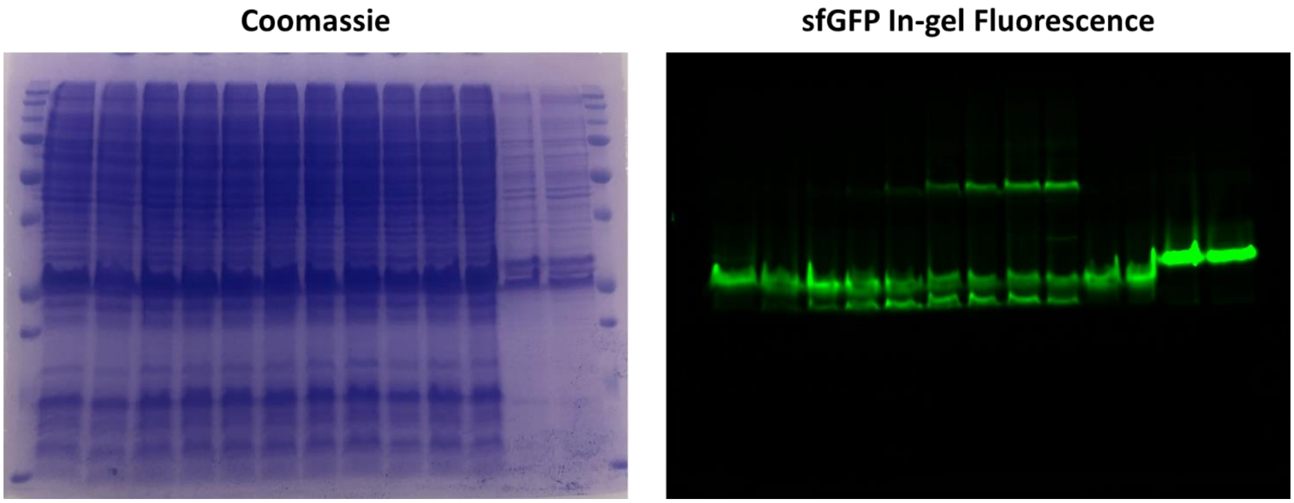

***Figure S22***. Unedited gel images for Figure 5D, including Coomassie-stained (left), and sfGFP in-gel fluorescence (right).

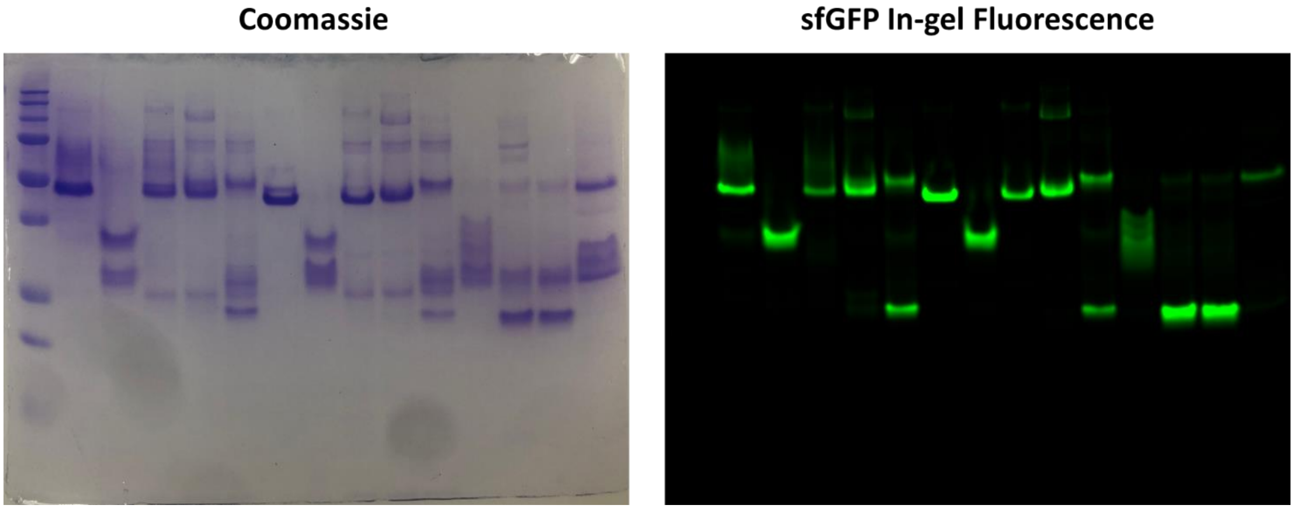

***Figure S23***. Unedited gel images for Figure 6B, including Coomassie-stained (left), and sfGFP in-gel fluorescence (right).

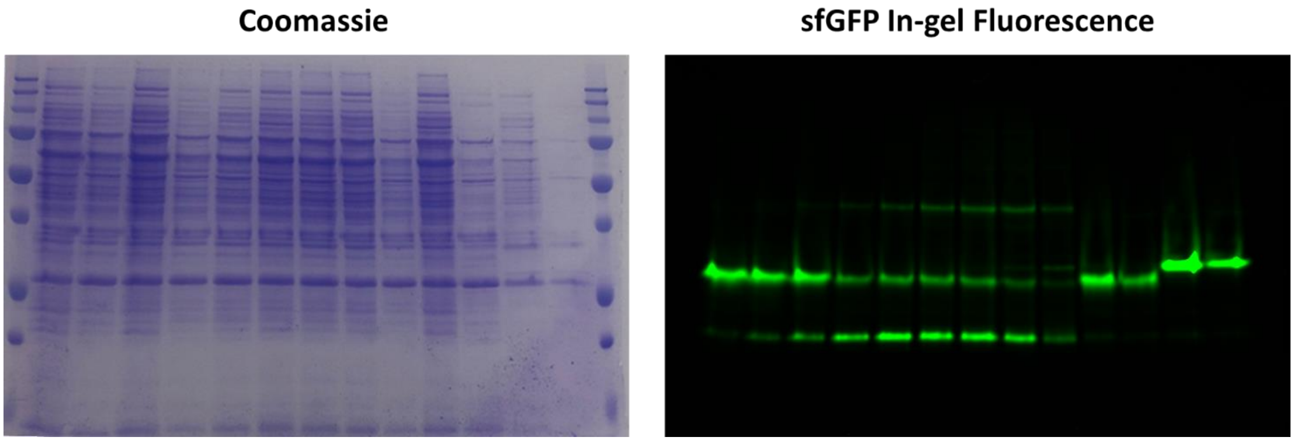

***Figure S24***. Unedited gel images for Figure 6B, including Coomassie-stained (left), and sfGFP in-gel fluorescence (right).

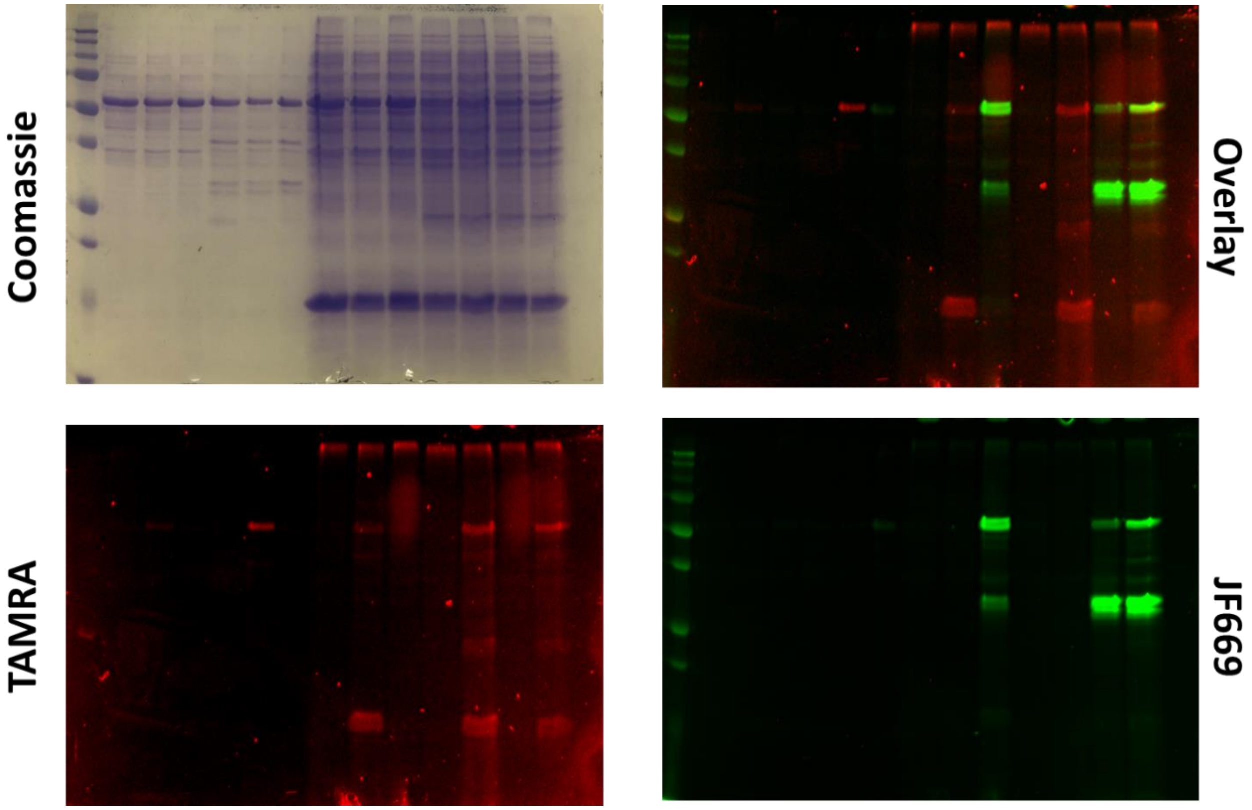

***Figure S25***. Unedited gel images for Figure 7B, including Coomassie-stained (top left), TAMRA (bottom left), JF669 (bottom right) channels, and their overlay (top right). Note, lower, high-abundance bands correspond to truncation product, which is a common byproduct of GCE and can experience specific (and non-specific) labeling due to their abundance and distribution.

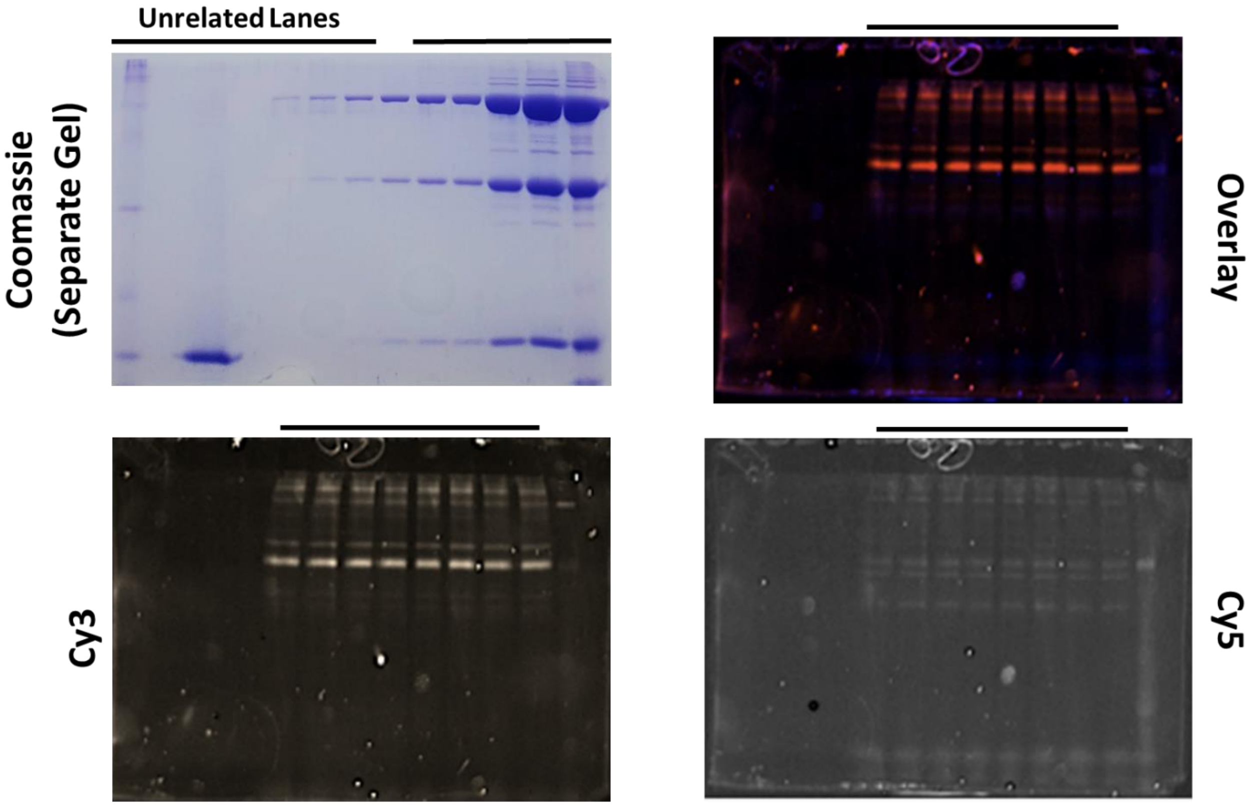

***Figure S26***. Unedited gel images for Figure 8B, including Coomassie-stained (top left), Cy3 (bottom left), Cy5 (bottom right) channels, and their overlay (top right). Bars indicate unrelated lanes. Note the Coomassie-stained gel (top left) is a separate gel from those presented in the other panels.

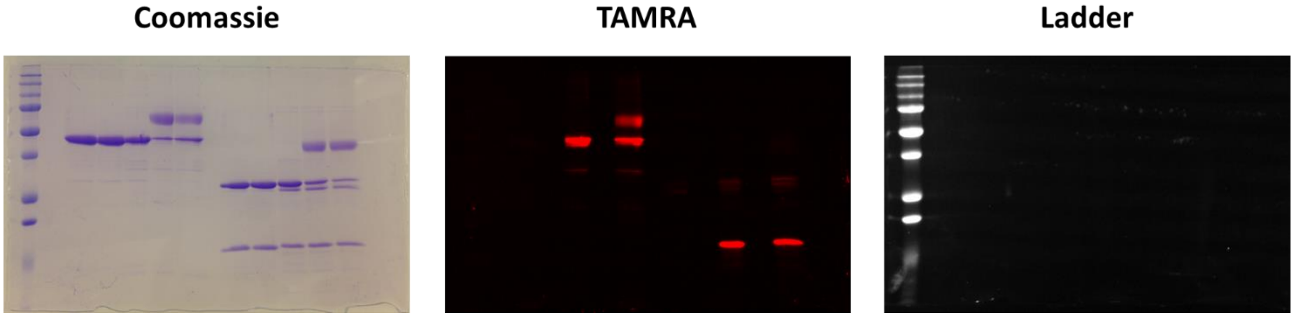

***Figure S27***. Unedited gel images for Figure S4B, including Coomassie-stained (left), TAMRA (middle) and fluorescent ladder (left) channels.

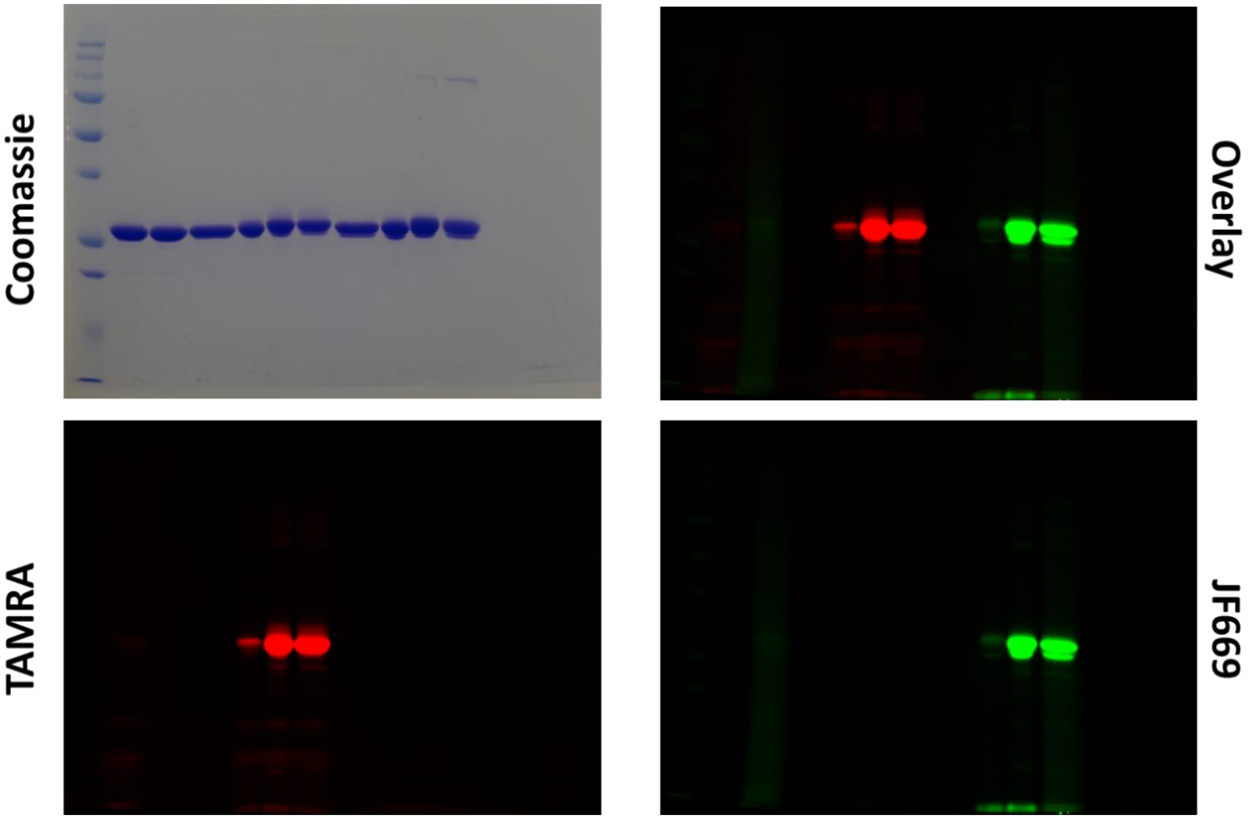

***Figure S28***. Unedited gel images for Figure S5A, including Coomassie-stained (top left), TAMRA (bottom left), JF669 (bottom right) channels, and their overlay (top right).

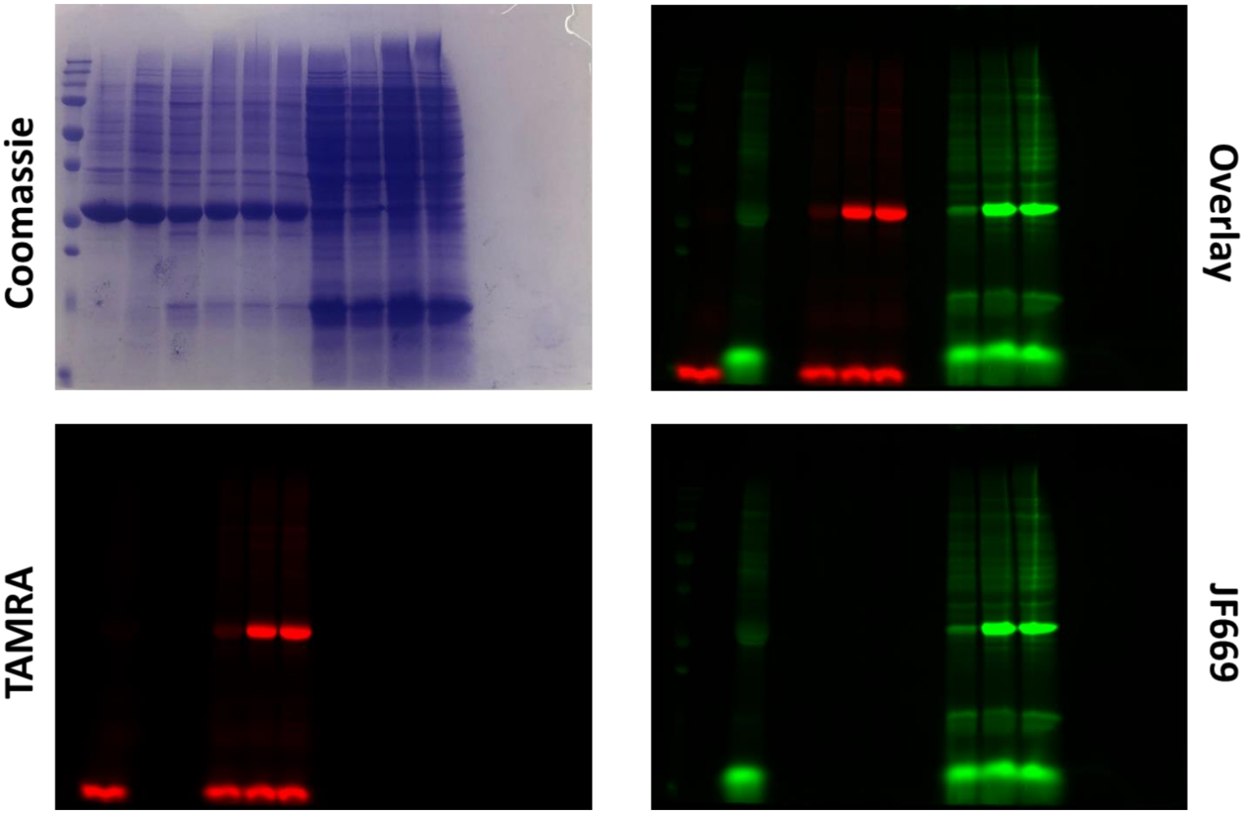

***Figure S29***. Unedited gel images for Figure S5C, including Coomassie-stained (top left), TAMRA (bottom left), JF669 (bottom right) channels, and their overlay (top right). Note, lower, high-abundance bands correspond to truncation product, which is a common byproduct of GCE and can experience specific (and non-specific) labeling due to their abundance and distribution.

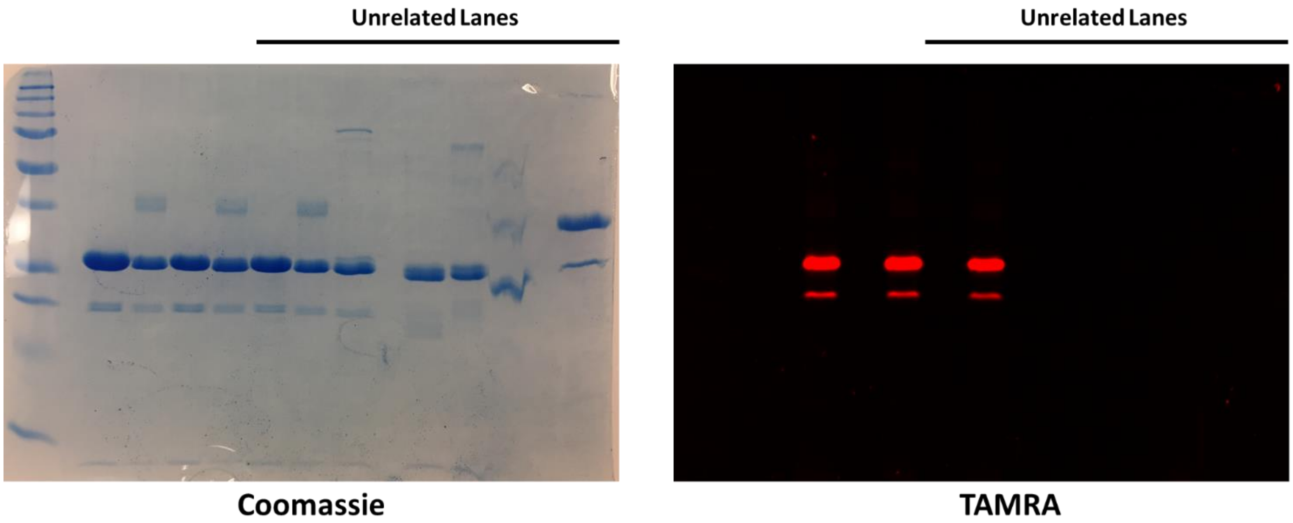

***Figure S30*** Unedited gel images for Figure S8C, including Coomassie-stained (left) and TAMRA (right) channels. Top bars indicate unrelated lanes

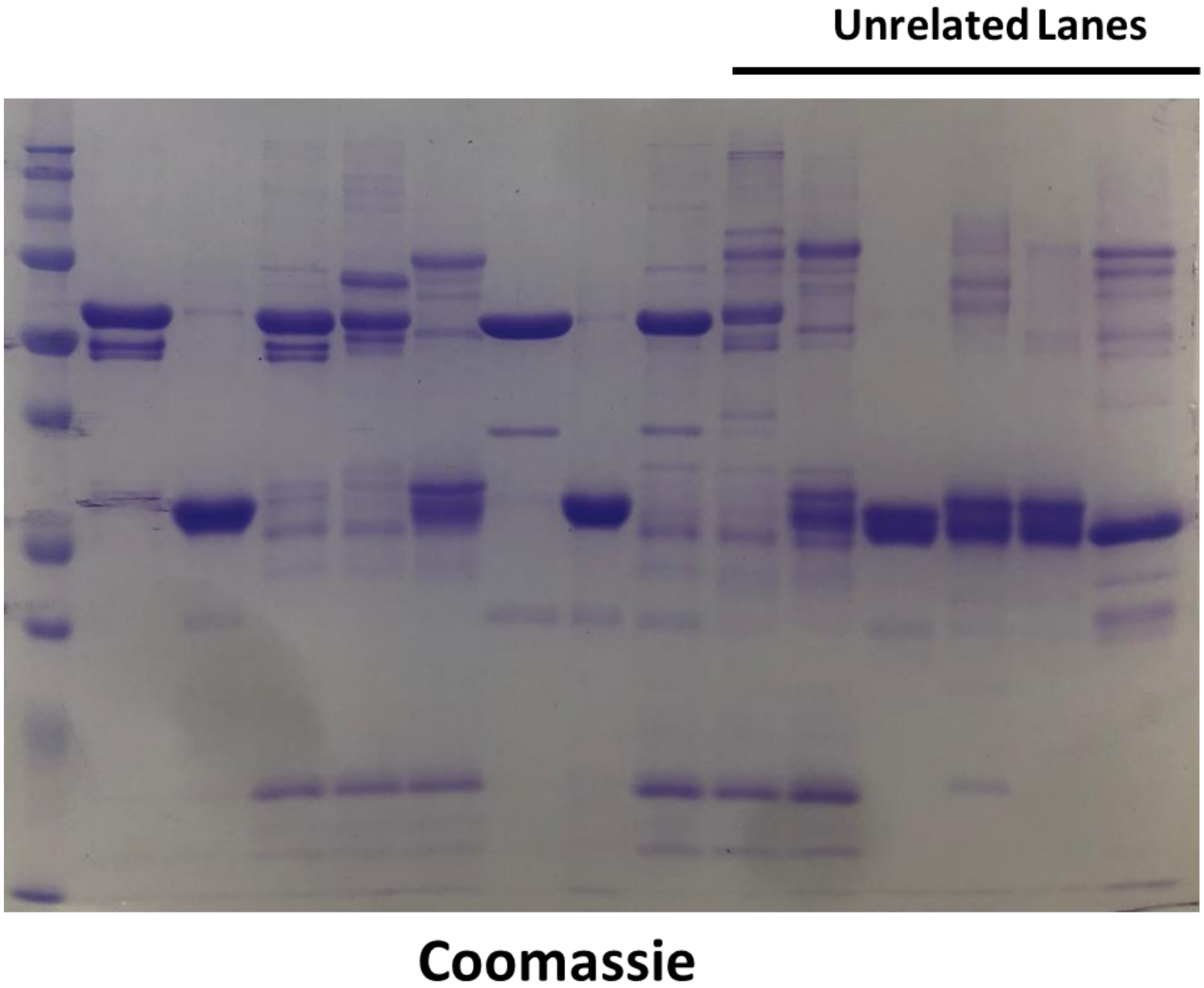

*Figure S31*. Unedited gel image for Figure S11. Top bar indicates unrelated lanes

#### Supplementary Tables

| No. | Plasmid Name | Promoter(s) | Ori | Resistance | Size (bp) | Primers | Template | Backbone | Cloning Approach | Reference | Addgene |
|---|---|---|---|---|---|---|---|---|---|---|---|
| **1** | pEVOL-pAzF[TAG] | araBAD (aaRS1)<br>glnS (aaRS2)<br>proK (tRNA) | p15A | Cm | 6104 | 1-6 | 48215 | pEVOL (BglII/PstI) | SLiCE | This Study | 164579 |
| **2** | pUltraI-pAzF[TAG] | tac (aaRS)<br>proK (tRNA) | CloDF13<br>(cdf) | Spec | 4977 | - | - | - | - | [20] | 48215 |
| **3** | pUltraI-Tet3.0[TAA] | tac (aaRS)<br>proK (tRNA) | CloDF13<br>(cdf) | Spec | 5297 | 6-12 | [1], 4 | pUltraI (NotI; PCR) | SLiCE | This Study | 164580 |
| **4** | pDule1-Tet3.0[TAA] | lpp (aaRS)<br>lpp (tRNA) | p15A | Tet | 6672 | 13-14 | [1] | N/A | QuikChange | This Study | - |
| **5** | pBAD-SUMO[WT]-sfGFP[WT] | araBAD | pBR322 | Amp | 5144 | 15-16 | 8 | pBAD (NcoI/XhoI) | SLiCE | This Study | - |
| **6** | pBAD-SUMO[E35TAG]-sfGFP[WT] | araBAD | pBR322 | Amp | 5144 | 15-16 | 9 | pBAD (NcoI/XhoI) | SLiCE | This Study | - |
| **7** | pBAD-SUMO[E35TAA]-sfGFP[WT] | araBAD | pBR322 | Amp | 5144 | 15-16 | 10 | pBAD (NcoI/XhoI) | SLiCE | This Study | - |
| **8** | pET28-SUMO[WT]-sfGFP[WT] | T7 | pBR322 | Kan | 6261 | - | - | - | - | [9] | - |
| **9** | pET28-SUMO[E35TAG]-sfGFP[WT] | T7 | pBR322 | Kan | 6261 | 17-20 | 8 | - | SLiCE | This Study | - |
| **10** | pET28-SUMO[E35TAA]-sfGFP[WT] | T7 | pBR322 | Kan | 6261 | 17, 20-22 | 8 | pET28 (NcoI/XhoI) | SLiCE | This Study | - |
| **11** | pET28-SUMO[E35TAG+T102TAA]-sfGFP[WT] | T7 | pBR322 | Kan | 6261 | 17-20, 25-26 | 8 | pET28 (NcoI/XhoI) | SLiCE | This Study | - |
| **12** | pET28-SUMO[E35TAA+T102TAG]-sfGFP[WT] | T7 | pBR322 | Kan | 6261 | 17, 20-24 | 8 | pET28 (NcoI/XhoI) | SLiCE | This Study | - |
| **13** | pET28-SUMO[WT]-sfGFP[N253TAG] | T7 | pBR322 | Kan | 6261 | - | - | - | - | [9] | - |
| **14** | pET28-SUMO[WT]-sfGFP[N253TAA] | T7 | pBR322 | Kan | 6261 | 17, 20, 30-31 | 13 | pET28 (NcoI/XhoI) | SLiCE | This Study | - |
| **15** | pET28-SUMO[E35TAG]-sfGFP[N253TAA] | T7 | pBR322 | Kan | 6261 | 17-20 | 13 | pET28 (NcoI/XhoI) | SLiCE | This Study | - |
| **16** | pET28-sfGFP[WT] | T7 | pBR322 | Kan | 5952 | - | - | - | - | [10] | 85493 |
| **17** | pET28-sfGFP[N150TAG] | T7 | pBR322 | Kan | 5952 | - | - | - | - | [10] | 85492 |
| **18** | pET28-sfGFP[N150TAA] | T7 | pBR322 | Kan | 5952 | 27, 30-32 | 16 | pET28 (NcoI/XhoI) | SLiCE | This Study | 164581 |
| **19** | pET28-sfGFP[D134TAG+N150TAA] | T7 | pBR322 | Kan | 5952 | 27-32 | 16 | pET28 (NcoI/XhoI) | SLiCE | This Study | 164582 |
| **20** | pET28-mTagBFP[WT] | T7 | pBR322 | Kan | 5928 | 33-34 | [30] | - | - | - | 34632 |
| **21** | pET28-mTagBFP[Y14TAG] | T7 | pBR322 | Kan | 5928 | 33-35 | 20 | pET28 (NcoI/XhoI) | SLiCE, Overlap Extension | This Study | - |
| **22** | pET28-mTagBFP[Y14TAA] | T7 | pBR322 | Kan | 5928 | 33-34, 36 | 20 | pET28 (NcoI/XhoI) | SLiCE, Overlap Extension | This Study | - |
| **23** | pET28-mTagBFP[T105TAG] | T7 | pBR322 | Kan | 5928 | 33-34, 37-38 | 20 | pET28 (NcoI/XhoI) | SLiCE | This Study | - |
| **24** | pET28-mTagBFP[T105TAA] | T7 | pBR322 | Kan | 5928 | 33-34, 39-40 | 20 | pET28 (NcoI/XhoI) | SLiCE | This Study | - |
| **25** | pET28-mTagBFP[D112TAG] | T7 | pBR322 | Kan | 5928 | 33-34, 41-42 | 20 | pET28 (NcoI/XhoI) | SLiCE | This Study | - |
| **26** | pET28-mTagBFP[D112TAA] | T7 | pBR322 | Kan | 5928 | 33-34, 43-44 | 20 | pET28 (NcoI/XhoI) | SLiCE | This Study | - |
| **27** | pET28-mTagBFP[T144TAG] | T7 | pBR322 | Kan | 5928 | 33-34, 45-46 | 20 | pET28 (NcoI/XhoI) | SLiCE | This Study | - |
| **28** | pET28-mTagBFP[T144TAA] | T7 | pBR322 | Kan | 5928 | 33-34, 47-48 | 20 | pET28 (NcoI/XhoI) | SLiCE | This Study | - |
| **29** | pET28-mTagBFP[E200TAG] | T7 | pBR322 | Kan | 5928 | 33-34, 49-50 | 20 | pET28 (NcoI/XhoI) | SLiCE | This Study | - |
| **30** | pET28-mTagBFP[E200TAA] | T7 | pBR322 | Kan | 5928 | 33-34, 51-52 | 20 | pET28 (NcoI/XhoI) | SLiCE | This Study | - |
| **31** | pET28-mTagBFP[N206TAG] | T7 | pBR322 | Kan | 5928 | 33-34, 53-54 | 20 | pET28 (NcoI/XhoI) | SLiCE | This Study | - |
| **32** | pET28-mTagBFP[N206TAA] | T7 | pBR322 | Kan | 5928 | 33-34, 55-56 | 20 | pET28 (NcoI/XhoI) | SLiCE | This Study | - |
| **33** | pET28-mTagBFP2[WT]-sfGFP[WT] | T7 | pBR322 | Kan | 6702 | 32-33, 57, 60 | 16, 20 | pET28 (NcoI/XhoI) | SLiCE | This Study | - |
| **34** | pET28-mTagBFP2[T105TAG]-sfGFP[N400TAA] | T7 | pBR322 | Kan | 6702 | 32-33, 57, 60 | 18, 23 | pET28 (NcoI/XhoI) | SLiCE | This Study | - |
| **35** | pET28-sfGFP[WT]-mTagBFP2[WT] | T7 | pBR322 | Kan | 6702 | 27, 32, 58-59 | 16, 20 | pET28 (NcoI/XhoI) | SLiCE | This Study | - |
| **36** | pET28-sfGFP[N150TAA]-mTagBFP2[T363TAG] | T7 | pBR322 | Kan | 6702 | 27, 32, 58-59 | 18, 23 | pET28 (NcoI/XhoI) | SLiCE | This Study | - |
| **37** | pETduet-mTagBFP2[WT]+sfGFP[WT] | T7 | pBR322 | Kan | 6812 | 33-34, 61-64 | 16, 20 | pETduet (NcoI/XhoI) | SLiCE, Overlap Extension | This Study | - |
| **38** | pETduet-mTagBFP2[T105TAG]+sfGFP[N150TAA] | T7 | pBR322 | Kan | 6812 | 33-34, 61-64 | 18, 23 | pETduet (NcoI/XhoI) | SLiCE, Overlap Extension | This Study | - |
| **39** | pETduet-sfGFP[WT]+mTagBFP2[WT] | T7 | pBR322 | Kan | 6812 | 27, 65-67 | 16, 20 | pETduet (NcoI/XhoI) | SLiCE, Overlap Extension | This Study | - |
| **40** | pETduet-sfGFP[N150TAA]+mTagBFP2[T105TAG] | T7 | pBR322 | Kan | 6812 | 27, 65-67 | 18, 23 | pETduet (NcoI/XhoI) | SLiCE, Overlap Extension | This Study | - |
| **41** | pETduet-sfGFP[WT]+mTagBFP2[WT] | T7 | pBR322 | Kan | 6812 | 27, 65-67 | 16, 20 | pETduet (NcoI/XhoI) | SLiCE, Overlap Extension | This Study | - |
| **42** | pETduet-sfGFP[N150TAA]+mTagBFP2[T105TAG] | T7 | pBR322 | Kan | 6812 | 27, 65-67 | 18, 23 | pETduet (NcoI/XhoI) | SLiCE, Overlap Extension | This Study | - |

***Table S1***. Plasmids used in this study, along with additional salient features, including promoters, origin of replication, resistance, size, and details pertaining to cloning approaches. For the cloning approach, the insert fragments were amplified using the indicated primers (number correspond to those in table S3) from an indicated template (numbers corresponding to plasmids in this table, or originating from the indicated reference in superscript), and were combined with a linearized backbone for SLiCE (restriction enzymes indicated in parentheses). In some cases, PCR fragments were fused prior to SLiCE using overlap extension PCR (indicated in cloning approach column). For QuikChange (plasmid 4), the entire plasmid was amplified using the indicated primers from the indicated template prior to Dpn1 digestion. If not prepared in this study, the reference (superscript; numbering consistent with supplemental references) and Addgene ID for the corresponding plasmids are indicated.

| Strain No. | Plasmid 1 | Plasmid 2 | Plasmid 3 | Cell Line | Resistance |
|---|---|---|---|---|---|
| 1 | pET28-SUMO[WT]-sfGFP[WT] | pEVOL-pAzF[TAG] | - | BL21(DE3) | Kan + Cm |
| 2 | pET28-SUMO[E35TAG]-sfGFP[WT] | pEVOL-pAzF[TAG] | - | BL21(DE3) | Kan + Cm |
| 3 | pET28-SUMO[WT]-sfGFP[WT] | pUltraI-pAzF[TAG] | - | BL21(DE3) | Kan + Spec |
| 4 | pET28-SUMO[E35TAG]-sfGFP[WT] | pUltraI-pAzF[TAG] | - | BL21(DE3) | Kan + Spec |
| 5 | pET28-SUMO[WT]-sfGFP[WT] | pUltraI-Tet3.0[TAA] | - | BL21(DE3) | Kan + Spec |
| 6 | pET28-SUMO[E35TAA]-sfGFP[WT] | pUltraI-Tet3.0[TAA] | - | BL21(DE3) | Kan + Spec |
| 7 | pET28-SUMO[WT]-sfGFP[WT] | pDule1-Tet3.0[TAA] | - | BL21(DE3) | Kan + Tet |
| 8 | pET28-SUMO[E35TAA]-sfGFP[WT] | pDule1-Tet3.0[TAA] | - | BL21(DE3) | Kan + Tet |
| 9 | pBAD-SUMO[WT]-sfGFP[WT] | pEVOL-pAzF[TAG] | - | DH10B | Amp + Cm |
| 10 | pBAD-SUMO[E35TAG]-sfGFP[WT] | pEVOL-pAzF[TAG] | - | DH10B | Amp + Cm |
| 11 | pBAD-SUMO[WT]-sfGFP[WT] | pUltraI-pAzF[TAG] | - | DH10B | Amp + Spec |
| 12 | pBAD-SUMO[E35TAG]-sfGFP[WT] | pUltraI-pAzF[TAG] | - | DH10B | Amp + Spec |
| 13 | pBAD-SUMO[WT]-sfGFP[WT] | pUltraI-Tet3.0[TAA] | - | DH10B | Amp + Spec |
| 14 | pBAD-SUMO[E35TAA]-sfGFP[WT] | pUltraI-Tet3.0[TAA] | - | DH10B | Amp + Spec |
| 15 | pBAD-SUMO[WT]-sfGFP[WT] | pDule1-Tet3.0[TAA] | - | DH10B | Amp + Tet |
| 16 | pBAD-SUMO[E35TAA]-sfGFP[WT] | pDule1-Tet3.0[TAA] | - | DH10B | Amp + Tet |
| 17 | pET28-SUMO[E35TAG]-sfGFP[WT] | pUltraI-Tet3.0[TAA] | - | BL21(DE3) | Kan + Spec |
| 18 | pET28-SUMO[E35TAA]-sfGFP[WT] | pEVOL-pAzF[TAG] | - | BL21(DE3) | Kan + Cm |
| 19 | pET28-sfGFP[N150TAG] | pEVOL-pAzF[TAG] | pUltraI-Tet3.0[TAA] | BL21(DE3) | Kan + Cm + Spec |
| 20 | pET28-SUMO[WT]-sfGFP[WT] | pEVOL-pAzF[TAG] | pUltraI-Tet3.0[TAA] | BL21(DE3) | Kan + Cm + Spec |
| 21 | pET28-SUMO[E35TAG+T102TAA]-sfGFP[WT] | pEVOL-pAzF[TAG] | pUltraI-Tet3.0[TAA] | BL21(DE3) | Kan + Cm + Spec |
| 22 | pET28-SUMO[E35TAA+T102TAG]-sfGFP[WT] | pEVOL-pAzF[TAG] | pUltraI-Tet3.0[TAA] | BL21(DE3) | Kan + Cm + Spec |
| 23 | pET28-SUMO[WT]-sfGFP[WT] | pUltraI-pAzF[TAG] | pDule1-Tet3.0[TAA] | BL21(DE3) | Kan + Spec + Tet |
| 24 | pET28-SUMO[E35TAG+T102TAA]-sfGFP[WT] | pUltraI-pAzF[TAG] | pDule1-Tet3.0[TAA] | BL21(DE3) | Kan + Spec + Tet |
| 25 | pET28-sfGFP[WT] | - | - | BL21(DE3) | Kan |
| 26 | pET28-sfGFP[N150TAG] | pEVOL-pAzF[TAG] | - | BL21(DE3) | Kan + Cm |
| 27 | pET28-sfGFP[N150TAA] | pUltraI-Tet3.0[TAA] | - | BL21(DE3) | Kan + Spec |
| 28 | pET28-sfGFP[D134TAG+N150TAA] | pEVOL-pAzF[TAG] | pUltraI-Tet3.0[TAA] | BL21(DE3) | Kan + Cm + Spec |
| 29 | pET28-SUMO[E35TAG]-sfGFP[N253TAA] | pEVOL-pAzF[TAG] | pUltraI-Tet3.0[TAA] | BL21(DE3) | Kan + Cm + Spec |
| 30 | pET28-SUMO[WT]-sfGFP[N253TAG] | pEVOL-pAzF[TAG] | - | BL21(DE3) | Kan + Cm |
| 31 | pET28-SUMO[WT]-sfGFP[N253TAA] | pUltraI-Tet3.0[TAA] | - | BL21(DE3) | Kan + Spec |
| 32 | pET28-mTagBFP[WT | - | - | BL21(DE3) | Kan |
| 33 | pET28-mTagBFP[Y14TAG] | pEVOL-pAzF[TAG] | - | BL21(DE3) | Kan + Cm |
| 34 | pET28-mTagBFP[Y14TAA] | pUltraI-Tet3.0[TAA] | - | BL21(DE3) | Kan + Spec |
| 35 | pET28-mTagBFP[T105TAG] | pEVOL-pAzF[TAG] | - | BL21(DE3) | Kan + Cm |
| 36 | pET28-mTagBFP[T105TAA] | pUltraI-Tet3.0[TAA] | - | BL21(DE3) | Kan + Spec |
| 37 | pET28-mTagBFP[D112TAG] | pEVOL-pAzF[TAG] | - | BL21(DE3) | Kan + Cm |
| 38 | pET28-mTagBFP[D112TAA] | pUltraI-Tet3.0[TAA] | - | BL21(DE3) | Kan + Spec |
| 39 | pET28-mTagBFP[T144TAG] | pEVOL-pAzF[TAG] | - | BL21(DE3) | Kan + Cm |
| 40 | pET28-mTagBFP[T144TAA] | pUltraI-Tet3.0[TAA] | - | BL21(DE3) | Kan + Spec |
| 41 | pET28-mTagBFP[E200TAG] | pEVOL-pAzF[TAG] | - | BL21(DE3) | Kan + Cm |
| 42 | pET28-mTagBFP[E200TAA] | pUltraI-Tet3.0[TAA] | - | BL21(DE3) | Kan + Spec |
| 43 | pET28-mTagBFP[N206TAG] | pEVOL-pAzF[TAG] | - | BL21(DE3) | Kan + Cm |
| 44 | pET28-mTagBFP[N206TAA] | pUltraI-Tet3.0[TAA] | - | BL21(DE3) | Kan + Spec |
| 45 | pET28-mTagBFP2[WT]-sfGFP[WT] | - | - | BL21(DE3) | Kan |
| 46 | pET28-mTagBFP2[T105TAG]-sfGFP[N400TAA] | pEVOL-pAzF[TAG] | pUltraI-Tet3.0[TAA] | BL21(DE3) | Kan + Cm + Spec |
| 47 | pET28-sfGFP[WT]-mTagBFP2[WT] | - | - | BL21(DE3) | Kan |
| 48 | pET28-sfGFP[N150TAA]-mTagBFP2[T363TAG] | pEVOL-pAzF[TAG] | pUltraI-Tet3.0[TAA] | BL21(DE3) | Kan + Cm + Spec |
| 49 | pET28-sfGFP[WT]-mTagBFP2[T363TAG] | pEVOL-pAzF[TAG] | - | BL21(DE3) | Kan + Cm |
| 50 | pET28-sfGFP[N150TAA]-mTagBFP2[WT] | pUltraI-Tet3.0[TAA] | - | BL21(DE3) | Kan + Spec |
| 51 | pETduet-mTagBFP2[WT]+sfGFP[WT] | - | - | BL21(DE3) | Kan |
| 52 | pETduet-mTagBFP2[T105TAG]+sfGFP[N150TAA] | pEVOL-pAzF[TAG] | pUltraI-Tet3.0[TAA] | BL21(DE3) | Kan + Cm + Spec |
| 53 | pETduet-sfGFP[WT]+mTagBFP2[WT] | - | - | BL21(DE3) | Kan |
| 54 | pETduet-sfGFP[N150TAA]+mTagBFP2[T105TAG] | pEVOL-pAzF[TAG] | pUltraI-Tet3.0[TAA] | BL21(DE3) | Kan + Cm + Spec |

***Table S2***. *E. coli* strains used in this study, including plasmid combinations, cell line, and resistance.

| No. | Primer | Sequence | Size (bp) |
|---|---|---|---|
| 1 | pEVOL-Mj-aaRS1 Fwd | TGGGCTAACAGGAGGAATTAGATCTATGGACGAATTTGAAATGATAAAGAGAAACAC | 57 |
| 2 | pEVOL-Mj-aaRS1(TAA>TGA) Rev | CCAAGCTGGAGACCGTTTAAACTCATCATAATCTCTTTCTAATTGGCTCTAAAATCTTTATAAGTTC | 67 |
| 3 | pEVOL_inter-aaRS Fwd | TGAGTTTAAACGGTCTCCAGCTTGG | 25 |
| 4 | pEVOL_inter-aaRS Rev | ATGGGATTCCTCAAAGCGTAAACAA | 25 |
| 5 | pEVOL-Mj-aaRS2 Fwd | TTGTTTACGCTTTGAGGAATCCCATATGGACGAATTTGAAATGATAAAGAGAAACA | 56 |
| 6 | pEVOL-Mj-aaRS2(TAA>TGA) Rev | GGCAATTTAGCGTTTGAAACTGCAG TCATAATCTCTTTCTAATTGGCTCTAAAATCTTTATAAGTTCTTC | 70 |
| 7 | pUltra1-Mb-aaRS Fwd | AATTTCACAAAGGAGGTGCGGCCGC ATGGATAAAAAACCGCTGGATGT | 48 |
| 8 | pUltra1-Mb-aaRS(TAA>TGA) Rev | CTGGAGACCGTTTAAACGCGGCCGC TCA CAGGTTCGTGCTAATGCC | 46 |
| 9 | pUltraI-Mb-tRNA Fwd | TGCGCAGTAAGATGCGCCCCGCATTGGAAACCTGATCATGTAGATCGAAC | 50 |
| 10 | pUltraI-Mb-tRNA Rev | GCTCGTTGAGCAGGCTTTTCGAATTTGGCGGAAACCCCG | 39 |
| 11 | pEVOL/pUltraI-Backbone(-tRNA) Rev | AATGCGGGGCGCATCTTACTGCGCA | 25 |
| 12 | pEVOL/pUltraI-Backbone(-tRNA) Fwd | AATTCGAAAAGCCTGCTCAACGAGC | 25 |
| 13 | Mb-tRNA-TAG>TAA Fwd | GATCGAACGGACTTTA AATCCGTTCAGCCGGGTTAGATTCCCGGGG | 46 |
| 14 | Mb-tRNA-TAG>TAA Rev | GCTGAACGGATTTAAAGTCCGTTCGATCTACATGATCAGGTTTCC | 45 |
| 15 | pBAD-SUMO Fwd | GTTTTTTGGGCTAACAGGAGGAATTAACCATGGGCAGCGACTCAGAAGTCAATC | 54 |
| 16 | pBAD-sfGFP-6xHis (TGA Stop) Rev | ATGGTACCAGCTGCAGATCTCGAGCTTCAGTGGTGGTGGTGGTGGTGCTCGAGGCTGC | 58 |
| 17 | pET28-SUMO-sfGFP Fwd | TTTAACTTTAAGAAGGAGATATACCATGGGCAGCGACTCAGAAGTCAATC | 50 |
| 18 | SUMO-E35TAG Rev | GATCTTGAAGAAGATCTATGAAGATCCATCGGACACCTTTAAATTG | 46 |
| 19 | SUMO-E35TAG Fwd | GATGGATCTTCATAGATCTTCTTCAAGATCAAAAAGACCACTCC | 44 |
| 20 | pET28-SUMO-sfGFP Rev | GCAGCCGGATCTCAGTGGTGGTGGTGGTGGTGCTCGAGGCTGCCTTTATACAGTTCATCC | 60 |
| 21 | SUMO-E35TAA Rev | GATCTTGAAGAAGATTTATGAAGATCCATCGGACACCTTTAAATTG | 46 |
| 22 | SUMO-E35TAA Fwd | GATGGATCTTCATAAATCTTCTTCAAGATCAAAAAGACCACTCC | 44 |
| 23 | SUMO-T102TAG Rev | CACCTTTGCTAACAGAGCCCTAACCGGATCCACCAATCTGTTCTCTGTGAGCCTC | 55 |
| 24 | SUMO-T102TAG Fwd | ACAGATTGGTGGATCCGGTTAGGGCTCTGTTAGCAAAGGTGAAGAACTGTTTACCG | 56 |
| 25 | SUMO-T102TAA Rev | CACCTTTGCTAACAGAGCCTTAACCGGATCCACCAATCTGTTCTCTGTGAGCCTC | 55 |
| 26 | SUMO-T102TAA Fwd | ACAGATTGGTGGATCCGGTTAAGGCTCTGTTAGCAAAGGTGAAGAACTGTTTACCG | 56 |
| 27 | pET28-sfGFP Fwd | TTTAACTTTAAGAAGGAGATATACCATGGTTAGCAAAGGTGAAGA | 45 |
| 28 | sfGFP-D134TAG Rev | CAGAATGTTGCCCTATTCTTTAAAATCAATACCTTTCAGTTCAATG | 45 |
| 29 | sfGFP-D134TAG Fwd | GATTTTAAAGAATAGGGCAACATTCTGGGTCATAAACTGGAATAT | 46 |
| 30 | sfGFP-N150TAA Rev | GGTAATATACACTTAATGGCTGTTGAAATTATATTCCAGTTTATG | 45 |
| 31 | sfGFP-N150TAA Fwd | TTCAACAGCCATTAAGTGTATATTACCGCCGATAAACAGAAAAATG | 46 |
| 32 | pET28-sfGFP Rev | ATCTCAGTGGTGGTGGTGGTGGTGCTCGAGGCTGCCTTTATACAGTTCATCCATA | 55 |
| 33 | pETDuet-mTagBFP2 Fwd | ATTTTGTTTAACTTTAAGAAGGAGATATACCATGAGCGAGCTGATTAAGGAGAACA | 56 |
| 34 | pETDuet-mTagBFP2 (TGA Stop) Rev | TCAGTGGTGGTGGTGGTGGTGCTCGAGATTAAGCTTGTGCCCCAGTTTGCTA | 52 |
| 35 | mTagBFP2-Y14TAG Fwd | ATGAGCGAGCTGATTAAGGAGAACATGCACATGAAGCTGTAGATGGAGGG | 50 |
| 36 | mTagBFP2-Y14TAA Fwd | ATGAGCGAGCTGATTAAGGAGAACATGCACATGAAGCTGTAAATGGAGGG | 50 |
| 37 | mTagBFP2-T105TAG Rev | CTGGTGTCCTGCTAAGCGGTCAGCA | 25 |
| 38 | mTagBFP2-T105TAG Fwd | TGCTGACCGCTTAGCAGGACACCAG | 25 |
| 39 | mTagBFP2-T105TAA Rev | CTGGTGTCCTGTTAAGCGGTCAGCA | 25 |
| 40 | mTagBFP2-T105TAA Fwd | TGCTGACCGCTTAACAGGACACCAG | 25 |
| 41 | mTagBFP2-D112TAG Rev | ATGAGGCAGCCCTACTGGAGGCTGG | 25 |
| 42 | mTagBFP2-D112TAG Fwd | CCAGCCTCCAGTAGGGCTGCCTCAT | 25 |
| 43 | mTagBFP2-D112TAA Rev | ATGAGGCAGCCTTACTGGAGGCTGG | 25 |
| 44 | mTagBFP2-D112TAA Fwd | CCAGCCTCCAGTAAGGCTGCCTCAT | 25 |
| 45 | mTagBFP2-T144TAG Rev | TACAGCGTCTCCTAGAAGGCCTCCC | 25 |
| 46 | mTagBFP2-T144TAG Fwd | GGGAGGCCTTCTAGGAGACGCTGTA | 25 |
| 47 | mTagBFP2-T144TAA Rev | TACAGCGTCTCTTAGAAGGCCTCCC | 25 |
| 48 | mTagBFP2-T144TAA Fwd | GGGAGGCCTTCTAAGAGACGCTGTA | 25 |
| 49 | mTagBFP2-E200TAG Rev | TCCTTGATTCTCTACAGTCTGTAGT | 25 |
| 50 | mTagBFP2-E200TAG Fwd | ACTACAGACTGTAGAGAATCAAGGA | 25 |
| 51 | mTagBFP2-E200TAA Rev | TCCTTGATTCTTTACAGTCTGTAGT | 25 |
| 52 | mTagBFP2-E200TAA Fwd | ACTACAGACTGTAAAGAATCAAGGA | 25 |
| 53 | mTagBFP2-N206TAG Rev | ACGTAGGTCTCCTAGTTGGCCTCCT | 25 |
| 54 | mTagBFP2-N206TAG Fwd | AGGAGGCCAACTAGGAGACCTACGT | 25 |
| 55 | mTagBFP2-N206TAA Rev | ACGTAGGTCTCTTAGTTGGCCTCCT | 25 |
| 56 | mTagBFP2-N206TAA Fwd | AGGAGGCCAACTAAGAGACCTACGT | 25 |

***Table S3***. Primers used in this study

| Gene | | Sequence |
|---|---|---|
| ***Mj*pAzFRS** | | ATGGACGAATTTGAAATGATAAAGAGAAACACATCTGAAATTATCAGCGAGGAAGAGTTAAGAGAGGTTTTAAAAAAAGATGAAAAATCTGCTCTGATAGGTTTTGAACCAAGTGGTAAAATACATTTAGGGCATTATCTCCAAATAAAAAAGATGATTGATTTACAAAATGCTGGATTTGATATAATTATAGTGTTGGCTGATTTACATGCCTATTTAAACCAGAAAGGAGAGTTGGATGAGATTAGAAAAATAGGAGATTATAACAAAAAAGTTTTTGAAGCAATGGGGTTAAAGGCAAAATATGTTTATGGAAGTGAATGGATGCTTGATAAGGATTATACACTGAATGTCTATAGATTGGCTTTAAAAACTACCTTAAAAAGAGCAAGAAGGAGTATGGAACTTATAGCAAGAGAGGATGAAAATCCAAAGGTTGCTGAAGTTATCTATCCAATAATGCAGGTTAATGGTCCTCATTATCTTGGCGTTGATGTTGCAGTTGGAGGGATGGAGCAGAGAAAAATACACATGTTAGCAAGGGAGCTTTTACCAAAAAAGGTTGTTTGTATTCACAACCCTGTCTTAACGGGTTTGGATGGAGAAGGAAAGATGAGTTCTTCAAAAGGGAATTTTATAGCTGTTGATGACTCTCCAGAAGAGATTAGGGCTAAGATAAAGAAAGCATACTGCCCAGCTGGAGTTGTTGAAGGAAATCCAATAATGGAGATAGCTAAATACTTCCTTGAATATCCTTTAACCATAAAAAGGCCAGAAAAATTTGGTGGAGATTTGACAGTTAATAGCTATGAGGAGTTAGAGAGTTTATTTAAAAATAAGGAATTGCATCCAATGGATTTAAAAAATGCTGTAGCTGAAGAACTTATAAAGATTTTAGAGCCAATTAGAAAGAGATTATGA |
| ***Mj*tRNA$^{CUA}$** | | CCGGCGGTAGTTCAGCAGGGCAGAACGGCGGACTCTAAATCCGCATGGCAGGGGTTCAAATCCCCTCCGCCGGACCA |
| ***Mb*Tet3.0RS** | | ATGGATAAAAAACCGCTGGATGTGCTGATTAGCGCGACCGGCCTGTGGATGAGCCGTACCGGCACCCTGCATAAAATCAAACATCATGAAGTGAGCCGCAGCAAAATCTATATTGAAATGGCGTGCGGCGATCATCTGGTGGTGAACAACAGCCGTAGCTGCCGTACCGCGCGTGCGTTTCGTCATCATAAATACCGCAAAACCTGCAAACGTTGCCGTGTGAGCGATGAAGATATCAACAACTTTCTGACCCGTAGCACCGAAAGCAAAAACAGCGTGAAAGTGCGTGTGGTGAGCGCGCCGAAAGTGAAAAAAGCGATGCCGAAAAGCGTGAGCCGTGCGCCGAAACCGCTGGAAAATAGCGTGAGCGCGAAAGCGAGCACCAACACCAGCCGTAGCGTTCCGAGCCCGGCGAAAAGCACCCCGAACAGCAGCGTTCCGGCGTCTGCGCCGGCACCGAGCCTGACCCGCAGCCAGCTGGATCGTGTGGAAGCGCTGCTGTCTCCGGAAGATAAAATTAGCCTGAACATGGCGAAACCGTTTCGTGAACTGGAACCGGAACTGGTGACCCGTCGTAAAAACGATTTTCAGCGCCTGTATACCAACGATCGTGAAGATTATCTGGGCAAACTGGAACGTGATATCACCAAATTTTTTGTGGATCGCGGCTTTCTGGAAATTAAAAGCCCGATTCTGATTCCGGCGGAATATGTGGAACGTATGGGCATTAACAACGACACCGAACTGAGCAAACAAATTTTCCGTGTGGATAAAAACCTGTGCCTGCGTCCGATGCTGGCCCCGACCGGTTATAACTATTTGCGTAAACTGGATCGTATTCTGCCGGGTCCGATCAAAATTTTTGAAGTGGGCCCGTGCTATCGCAAAGAAAGCGATGGCAAAGAACACCTGGAAGAATTCACCATGGTTGGTTTTGCTCAAATGGGCAGCGGCTGCACCCGTGAAAACCTGGAAGCGCTGATCAAAGAATTCCTGGATTATCTGGAAATCGACTTCGAAATTGTGGGCGATAGCTGCATGGTGTATGGCGATACCCTGGATATTATGCATGGCGATCTGGAACTGAGCAGCGCGGTGGTGGGTCCGGTTAGCCTGGATCGTGAATGGGGCATTGATAAACCGTGGATTGGCGCGGGTTTTGGCCTGGAACGTCTGCTGAAAGTGATGCATGGCTTCAAAAACATTAAACGTGCGAGCCGTAGCGAAAGCTACTATAACGGCATTAGCACGAACCTGTAA |
| ***Mm*tRNA$^{UUA}$** | | GGAAACCTGATCATGTAGATCGAACGGACTTTAAATCCGTTCAGCCGGGTTAGATTCCCGGGGTTTCCGCCA |
| **SUMO-sfGFP** | | ATGGGCAGCGACTCAGAAGTCAATCAAGAAGCTAAGCCAGAGGTCAAGCCAGAAGTCAAGCCTGAGACTCACATCAATTTAAAGGTGTCCGATGGATCTTCAGAGATCTTCTTCAAGATCAAAAAGACCACTCCTTTAAGAAGGCTGATGGAAGCGTTCGCTAAAAGACAGGGTAAGGAAATGGACTCCTTAAGATTCTTGTACGACGGTATTAGAATCCAAGCTGATCAGACCCCTGAAGATTTGGACATGGAGGATAACGATATTATTGAGGCTCACAGAGAACAGATTGGTGGATCCGGTACCGGCTCTGTTAGCAAAGGTGAAGAACTGTTTACCGGCGTTGTGCCGATTCTGGTGGAACTGGATGGTGATGTGAATGGCCATAAATTTAGCGTTCGTGGCGAAGGCGAAGGTGATGCGACCAACGGTAAACTGACCCTGAAATTTATTTGCACCACCGGTAAACTGCCGGTTCCGTGGCCGACCCTGGTGACCACCCTGACCTATGGCGTTCAGTGCTTTAGCCGCTATCCGGATCATATGAAACGCCATGATTTCTTTAAAAGCGCGATGCCGGAAGGCTATGTGCAGGAACGTACCATTAGCTTCAAAGATGATGGCACCTATAAAACCCGTGCGGAAGTTAAATTTGAAGGCGATACCCTGGTGAACCGCATTGAACTGAAAGGTATTGATTTTAAAGAAGATGGCAACATTCTGGGTCATAAACTGGAATATAATTTCAACAGCCATAATGTGTATATTACCGCCGATAAACAGAAAAATGGCATCAAAGCGAACTTTAAAATCCGTCACAACGTGGAAGATGGTAGCGTGCAGCTGGCGGATCATTATCAGCAGAATACCCCGATTGGTGATGGCCCGGTGCTGCTGCCGGATAATCATTATCTGAGCACCCAGAGCGTTCTGAGCAAAGATCCGAATGAAAAACGTGATCATATGGTGCTGCTGGAATTTGTTACCGCCGCGGGCATTACCCACGGTATGGATGAACTGTATAAAGGCAGCCTCGAGCACCACCACCACCACCACTGA |
| **sfGFP*** | | ATGGTTAGCAAAGGTGAAGAACTGTTTACCGGCGTTGTGCCGATTCTGGTGGAACTGGATGGTGATGTGAATGGCCATAAATTTAGCGTTCGTGGCGAAGGCGAAGGTGATGCGACCAACGGTAAACTGACCCTGAAATTTATTTGCACCACCGGTAAACTGCCGGTTCCGTGGCCGACCCTGGTGACCACCCTGACCTATGGCGTTCAGTGCTTTAGCCGCTATCCGGATCATATGAAACGCCATGATTTCTTTAAAAGCGCGATGCCGGAAGGCTATGTGCAGGAACGTACCATTAGCTTCAAAGATGATGGCACCTATAAAACCCGTGCGGAAGTTAAATTTGAAGGCGATACCCTGGTGAACCGCATTGAACTGAAAGGTATTGATTTTAAAGAAGATGGCAACATTCTGGGTCATAAACTGGAATATAATTTCAACAGCCATAATGTGTATATTACCGCCGATAAACAGAAAAATGGCATCAAAGCGAACTTTAAAATCCGTCACAACGTGGAAGATGGTAGCGTGCAGCTGGCGGATCATTATCAGCAGAATACCCCGATTGGTGATGGCCCGGTGCTGCTGCCGGATAATCATTATCTGAGCACCCAGAGCGTTCTGAGCAAAGATCCGAATGAAAAACGTGATCATATGGTGCTGCTGGAATTTGTTACCGCCGCGGGCATTACCCACGGTATGGATGAACTGTATAAAGGCAGC[CTCGAG]CTCGAGCACCACCACCACCACCACTGA |
| **mTagBFP2** | | ATGAGCGAGCTGATTAAGGAGAACATGCACATGAAGCTGTACATGGAGGGCACCGTGGACAACCATCACTTCAAGTGCACATCCGAGGGCGAAGGCAAGCCCTACGAGGGCACCCAGACCATGAGAATCAAGGTGGTCGAGGGCGGCCCTCTCCCCTTCGCCTTCGACATCCTGGCTACTAGCTTCCTCTACGGCAGCAAGACCTTCATCAACCACACCCAGGGCATCCCCGACTTCTTCAAGCAGTCCTTCCCTGAGGGCTTCACATGGGAGAGAGTCACCACATACGAAGACGGGGGCGTGCTGACCGCTACCCAGGACACCAGCCTCCAGGACGGCTGCCTCATCTACAACGTCAAGATCAGAGGGGTGAACTTCACATCCAACGGCCCTGTGATGCAGAAGAAAACACTCGGCTGGGAGGCCTTCACCGAGACGCTGTACCCCGCTGACGGCGGCCTGGAAGGCAGAAACGACATGGCCCTGAAGCTCGTGGGCGGGAGCCATCTGATCGCAAACGCCAAGACCACATATAGATCCAAGAAACCCGCTAAGAACCTCAAGATGCCTGGCGTCTACTATGTGGACTACAGACTGGAAAGAATCAAGGAGGCCAACAACGAGACCTACGTCGAGCAGCACGAGGTGGCAGTGGCCAGATACTGCGACCTCCCTAGCAAACTGGGGCACAAGCTTAATCTCGAGCACCACCACCACCACCACTGA |
| **sfGFP-mTagBFP2** | | ATGGTTAGCAAAGGTGAAGAACTGTTTACCGGCGTTGTGCCGATTCTGGTGGAACTGGATGGTGATGTGAATGGCCATAAATTTAGCGTTCGTGGCGAAGGCGAAGGTGATGCGACCAACGGTAAACTGACCCTGAAATTTATTTGCACCACCGGTAAACTGCCGGTTCCGTGGCCGACCCTGGTGACCACCCTGACCTATGGCGTTCAGTGCTTTAGCCGCTATCCGGATCATATGAAACGCCATGATTTCTTTAAAAGCGCGATGCCGGAAGGCTATGTGCAGGAACGTACCATTAGCTTCAAAGATGATGGCACCTATAAAACCCGTGCGGAAGTTAAATTTGAAGGCGATACCCTGGTGAACCGCATTGAACTGAAAGGTATTGATTTTAAAGAAGATGGCAACATTCTGGGTCATAAACTGGAATATAATTTCAACAGCCATAATGTGTATATTACCGCCGATAAACAGAAAAATGGCATCAAAGCGAACTTTAAAATCCGTCACAACGTGGAAGATGGTAGCGTGCAGCTGGCGGATCATTATCAGCAGAATACCCCGATTGGTGATGGCCCGGTGCTGCTGCCGGATAATCATTATCTGAGCACCCAGAGCGTTCTGAGCAAAGATCCGAATGAAAAACGTGATCATATGGTGCTGCTGGAATTTGTTACCGCCGCGGGCATTACCCACGGTATGGATGAACTGTATAAAGGCAGCGGCAGTGGTTCTGGTGAAAACCTGTATTTCCAGTCCGGCAGCGGCTCGGGTATGAGCGAGCTGATTAAGGAGAACATGCACATGAAGCTGTACATGGAGGGCACCGTGGACAACCATCACTTCAAGTGCACATCCGAGGGCGAAGGCAAGCCCTACGAGGGCACCCAGACCATGAGAATCAAGGTGGTCGAGGGCGGCCCTCTCCCCTTCGCCTTCGACATCCTGGCTACTAGCTTCCTCTACGGCAGCAAGACCTTCATCAACCACACCCAGGGCATCCCCGACTTCTTCAAGCAGTCCTTCCCTGAGGGCTTCACATGGGAGAGAGTCACCACATACGAAGACGGGGGCGTGCTGACCGCTACCCAGGACACCAGCCTCCAGGACGGCTGCCTCATCTACAACGTCAAGATCAGAGGGGTGAACTTCACATCCAACGGCCCTGTGATGCAGAAGAAAACACTCGGCTGGGAGGCCTTCACCGAGACGCTGTACCCCGCTGACGGCGGCCTGGAAGGCAGAAACGACATGGCCCTGAAGCTCGTGGGCGGGAGCCATCTGATCGCAAACGCCAAGACCACATATAGATCCAAGAAACCCGCTAAGAACCTCAAGATGCCTGGCGTCTACTATGTGGACTACAGACTGGAAAGAATCAAGGAGGCCAACAACGAGACCTACGTCGAGCAGCACGAGGTGGCAGTGGCCAGATACTGCGACCTCCCTAGCAAACTGGGGCACAAGCTTAATCTCGAGCACCACCACCACCACCACTGA |
| **mTagBFP2-sfGFP** | | ATGAGCGAGCTGATTAAGGAGAACATGCACATGAAGCTGTACATGGAGGGCACCGTGGACAACCATCACTTCAAGTGCACATCCGAGGGCGAAGGCAAGCCCTACGAGGGCACCCAGACCATGAGAATCAAGGTGGTCGAGGGCGGCCCTCTCCCCTTCGCCTTCGACATCCTGGCTACTAGCTTCCTCTACGGCAGCAAGACCTTCATCAACCACACCCAGGGCATCCCCGACTTCTTCAAGCAGTCCTTCCCTGAGGGCTTCACATGGGAGAGAGTCACCACATACGAAGACGGGGGCGTGCTGACCGCTACCCAGGACACCAGCCTCCAGGACGGCTGCCTCATCTACAACGTCAAGATCAGAGGGGTGAACTTCACATCCAACGGCCCTGTGATGCAGAAGAAAACACTCGGCTGGGAGGCCTTCACCGAGACGCTGTACCCCGCTGACGGCGGCCTGGAAGGCAGAAACGACATGGCCCTGAAGCTCGTGGGCGGGAGCCATCTGATCGCAAACGCCAAGACCACATATAGATCCAAGAAACCCGCTAAGAACCTCAAGATGCCTGGCGTCTACTATGTGGACTACAGACTGGAAAGAATCAAGGAGGCCAACAACGAGACCTACGTCGAGCAGCACGAGGTGGCAGTGGCCAGATACTGCGACCTCCCTAGCAAACTGGGGCACAAGCTTAATGGCAGTGGTTCTGGTGAAAACCTGTATTTCCAGTCCGGCAGCGGCTCGGGTATGGTTAGCAAAGGTGAAGAACTGTTTACCGGCGTTGTGCCGATTCTGGTGGAACTGGATGGTGATGTGAATGGCCATAAATTTAGCGTTCGTGGCGAAGGCGAAGGTGATGCGACCAACGGTAAACTGACCCTGAAATTTATTTGCACCACCGGTAAACTGCCGGTTCCGTGGCCGACCCTGGTGACCACCCTGACCTATGGCGTTCAGTGCTTTAGCCGCTATCCGGATCATATGAAACGCCATGATTTCTTTAAAAGCGCGATGCCGGAAGGCTATGTGCAGGAACGTACCATTAGCTTCAAAGATGATGGCACCTATAAAACCCGTGCGGAAGTTAAATTTGAAGGCGATACCCTGGTGAACCGCATTGAACTGAAAGGTATTGATTTTAAAGAAGATGGCAACATTCTGGGTCATAAACTGGAATATAATTTCAACAGCCATAATGTGTATATTACCGCCGATAAACAGAAAAATGGCATCAAAGCGAACTTTAAAATCCGTCACAACGTGGAAGATGGTAGCGTGCAGCTGGCGGATCATTATCAGCAGAATACCCCGATTGGTGATGGCCCGGTGCTGCTGCCGGATAATCATTATCTGAGCACCCAGAGCGTTCTGAGCAAAGATCCGAATGAAAAACGTGATCATATGGTGCTGCTGGAATTTGTTACCGCCGCGGGCATTACCCACGGTATGGATGAACTGTATAAAGGCAGCCTCGAGCACCACCACCACCACCACTGA |
| ***Sc*RPA** | **RPA70** | ATGAGCAGCGTGCAGCTGAGCCGTGGTGACTTCCACAGCATCTTTACCAACAAACAGCGTTACGATAACCCGACCGGTGGCGTGTACCAAGTTTATAACACCCGTAAAAGCGACGGTGCGAACAGCAACCGTAAGAACCTGATCATGATTAGCGATGGCATCTATCACATGAAAGCGCTGCTGCGTAACCAGGCGGCGAGCAAGTTCCAGAGCATGGAGCTGCAACGTGGTGACATCATTCGTGTGATCATTGCGGAGCCGGCGATTGTTCGTGAGCGTAAGAAATACGTGCTGCTGGTTGACGATTTTGAACTGGTGCAGAGCCGTGCGGACATGGTTAACCAAACCAGCACCTTCCTGGATAACTATTTTAGCGAGCACCCGAACGAAACCCTGAAGGACGAGGACATCACCGACAGCGGTAACGTGGCGAACCAGACCAACGCGAGCAACGCGGGCGTTCCGGATATGCTGCACAGCAACAGCAACCTGAACGCGAACGAGCGTAAATTCGCGAACGAAAACCCGAACAGCCAAAAGACCCGTCCGATCTTTGCGATTGAGCAGCTGAGCCCGTACCAAAACGTGTGGACCATCAAAGCGCGTGTTAGCTATAAGGGTGAAATTAAAACCTGGCACAACCAGCGTGGTGACGGCAAGCTGTTCAACGTTAACTTTCTGGATACCAGCGGTGAAATCCGTGCGACCGCGTTCAACGACTTTGCGACCAAATTCAACGAGATTCTGCAAGAAGGCAAGGTGTACTATGTTAGCAAGGCGAAACTGCAGCCGGCGAAACCGCAATTCACCAACCTGACCCACCCGTACGAGCTGAACCTGGACCGTGATACCGTGATCGAGGAATGCTTTGATGAAAGCAACGTTCCGAAGACCCACTTCAACTTTATCAAACTGGACGCGATTCAGAACCAAGAGGTGAACAGCAACGTGGATGTTCTGGGTATCATTCAGACCATTAACCCGCACTTCGAACTGACCAGCCGTGCGGGCAAGAAATTTGACCGTCGTGACATCACCATTGTGGACGATAGCGGTTTCAGCATCAGCGTTGGCCTGTGGAACCAGCAAGCGCTGGACTTTAACCTGCCGGAAGGTAGCGTTGCGGCGATTAAAGGCGTGCGTGTTACCGATTTCGGTGGCAAGAGCCTGAGCATGGGTTTTAGCAGCACCCTGATCCCGAACCCGGAGATTCCGGAAGCGTACGCGCTGAAGGGTTGGTATGACAGCAAAGGCCGTAACGCGAACTTCATCACCCTGAAACAAGAGCCGGGTATGGGTGGCCAAAGCGCGGCGAGCCTGACCAAGTTTATTGCGCAGCGTATCACCATTGCGCGTGCGCAAGCGGAGAACCTGGGTCGTAGCGAAAAGGGCGACTTCTTTAGCGTGAAAGCGGCGATCAGCTTCCTGAAGGTTGATAACTTTGCGTACCCGGCGTGCAGCAACGAGAACTGCAACAAGAAAGTGCTGGAACAGCCGGACGGCACCTGGCGTTGCGAGAAATGCGATACCAACAACGCGCGTCCGAACTGGCGTTATATCCTGACCATTAGCATCATTGACGAAACCAACCAACTGTGGCTGACCCTGTTCGACGATCAGGCGAAGCAACTGCTGGGCGTGGACGCGAACACCCTGATGAGCCTGAAAGAGGAAGATCCGAACGAGTTCACCAAGATCACCCAGAGCATTCAAATGAACGAGTACGATTTTCGTATCCGTGCGCGTGAAGACACCTACAACGATCAGAGCCGTATTCGTTATACCGTTGCGAACCTGCACAGCCTGAACTACCGTGCGGAGGCGGACTATCTGGCGGATGAACTGAGCAAGGCGCTGCTGGCGTGA |
| | **RPA32** | ATGGCGACCTACCAGCCGTATAACGAGTACAGCAGCGTGACCGGTGGCGGTTTCGAGAACAGCGAAAGCCGTCCGGGCAGCGGCGAGAGCGAAACCAACACCCGTGTGAACACCCTGACCCCGGTTACCATCAAGCAGATTCTGGAGAGCAAACAGGACATCCAAGATGGCCCGTTCGTTAGCCACAACCAAGAACTGCACCACGTGTGCTTTGTTGGTGTGGTTCGTAACATCACCGACCACACCGCGAACATCTTTCTGACCATTGAAGATGGCACCGGTCAGATTGAAGTGCGTAAGTGGAGCGAAGACGCGAACGATCTGGCGGCGGGTAACGATGATAGCAGCGGTAAAGGTTATGGCAGCCAGGTTGCGCAGCAATTCGAGATCGGCGGTTACGTGAAGGTTTTCGGTGCGCTGAAAGAATTTGGCGGTAAGAAAAACATTCAATATGCGGTGATCAAGCCGATTGACAGCTTTAACGAGGTGCTGACCCACCACCTGGAAGTTATCAAGTGCCACAGCATTGCGAGCGGTATGATGAAACAGCCGCTGGAGAGCGCGAGCAACAACAACGGCCAAAGCCTGTTCGTTAAGGACGATAACGATACCAGCAGCGGTAGCAGCCCGCTGCAGCGTATCCTGGAGTTCTGCAAGAAACAATGCGAAGGCAAAGATGCGAACAGCTTTGCGGTGCCGATCCCGCTGATTAGCCAGAGCCTGAACCTGGACGAAACCACCGTTCGTAACTGCTGCACCACCCTGACCGATCAAGGTTTCATTTACCCGACCTTTGACGATAACAACTTCTTTGCGCTGCATCATCATCACCACCACTGA |
| | **RPA14** | ATGGCCAGCGAAACACCAAGAGTTGACCCCACAGAAATCTCCAACGTCAATGCTCCTGTGTTTAGGATAATAGCGCAAATCAAATCACAACCCACTGAATCTCAGTTGATTTTACAATCGCCAACCATATCATCGAAAAACGGCAGCGAGGTTGAAATGATTACGTTGAACAATATTCGTGTATCTATGAATAAAACATTTGAGATCGACTCGTGGTATGAGTTTGTTTGCAGGAACAATGATGACGGCGAGCTAGGATTTTTGATTCTAGACGCTGTACTATGCAAATTCAAGGAGAACGAAGATTTAAGCTTAAACGGTGTGGTTGCTTTACAGAGACTATGTAAGAAATACCCAGAAATATACTGA |

***Table S4***. DNA sequences of the key proteins in this study. Highlighted residues indicate location of mutagenesis to TAG (red), TAA (blue), or either (pink) codons within the various constructs used herein. *sfGFP variants containing N150TAA and D134TAG+N150TAA modifications possess additional "LE" residues at the N-terminal region preceding the 6x His-tag, the sequence of which is indicated in brackets ("[]").

| No. | Strain | Protein | Yield (mg/L) | ε ($A_{280}$) |
|---|---|---|---|---|
| **1** | N/A | SUMO[WT]-sfGFP[WT] | 300†[9] | 25,360 |
| **2** | 2 | SUMO[pAzF]-sfGFP[WT] | 46* | 25,360 |
| **3** | 31 | SUMO[WT]-sfGFP[Tet3.0] | 11 | 37,014 |
| **4** | 29 | SUMO[pAzF]-sfGFP[Tet3.0] | 11 | 37,014 |
| **5** | N/A | sfGFP[WT] | 220†[32] | 24,080 |
| **6** | 26 | sfGFP[pAzF] | 132* | 24,080 |
| **7** | 27 | sfGFP[Tet3.0] | 29 | 35,734 |
| **8** | 28 | sfGFP[Dual] | 26 | 35,734 |
| **9** | 32 | mTagBFP2[WT] | 93 | 26,740 |
| **10** | 33 | mTagBFP2-Y14[pAzF] | 83 | 26,740 |
| **11** | 34 | mTagBFP2-Y14[Tet3.0] | 8 | 38,394 |
| **12** | 35 | mTagBFP2-T105[pAzF] | 79 | 26,740 |
| **13** | 41 | mTagBFP2-E200[pAzF] | 34 | 26,740 |
| **14** | 43 | mTagBFP2-N206[pAzF] | 92 | 26,740 |
| **15** | 51 | mTagBFP2[WT] | 71 | 50,600** |
| | | sfGFP[WT] | 20 | 83,000** |
| **16** | 52 | mTagBFP2[pAzF] | 67 | 50,600** |
| | | sfGFP[Tet3.0] | 10 | 83,000** |
| **17** | 53 | sfGFP[WT] | 15 | 83,000** |
| | | mTagBFP2[WT] | 88 | 50,600** |
| **18** | 54 | sfGFP[Tet3.0] | 9 | 83,000** |
| | | mTagBFP2[pAzF] | 113 | 50,600** |
| **19** | 45 | mTagBFP2[WT]-sfGFP[WT] | 16 | 62,474 |
| **20** | 46 | mTagBFP2[pAzF]-sfGFP[Tet3.0] | 2 | 62,474 |
| **21** | 47 | sfGFP[WT]-mTagBFP2[pAzF] | 34 | 62,474 |
| **22** | 48 | sfGFP[Tet3.0]-mTagBFP2[pAzF] | 2 | 62,474 |

***Table S5***. Yields of select proteins used in this study in mg/L of media (as estimated by amount of purified protein isolated from a 50 mL expression). Indicated strain numbers correspond to those reported in table S2. "*" Indicates that yield was calculated under resin-limiting conditions and is therefore an underestimation of full yield. "†" Indicates that the reported yield was determined previously in the indicated reference. For protein concentration determinations ($M^{-1}$ $cm^{-1}$), the molar extinction coefficient is listed for each protein (note, for Tet3.0-containing proteins, 11,654 $M^{-1}$ $cm^{-1}$ was added to ε to account for strong absorption at this wavelength by this ncAA). ** To determine concentrations of sfGFP and mTagBFP2 in mixtures, the ε for sfGFP at 485 nm and for mTagBFP2 at 399 nm were used. All molar extinction coefficients values were estimated from protein sequences *in silico*.

| Figure | Spectrum | Peak | Observed Mass (Da) | Predicted Mass (Da) | Difference (Da) | Residue at D134 | Residue at N150 | Modification(s) | Notes |
|---|---|---|---|---|---|---|---|---|---|
| 3B | sfGFP$^{WT}$ | sfGFP$^{WT}$ | 27828 | - | - | Asp | Asn | - | |
| | | sfGFP$^{WT}$* | 27697 | 27696 | +1 | Asp | Asn | -Met | |
| | sfGFP$^{pAzF}$ | sfGFP$^{pAzF}$ | 27902 | 27902 | 0 | Asp | pAzF | - | |
| | | sfGFP$^{pAzF}$† | 27873 | 27876 | -3 | Asp | pAmF | pAzF Reduction | |
| | | sfGFP$^{pAzF}$* | 27771 | 27770 | +1 | Asp | pAzF | -Met | |
| | sfGFP$^{Tet3.0}$ | sfGFP$^{Tet3.0}$ | 28197 | 28197 | 0 | Asp | Tet3.0 | - | |
| | | sfGFP$^{Tet3.0}$* | 28066 | 28065 | +1 | Asp | Tet3.0 | -Met | |
| | sfGFP$^{Dual}$ | sfGFP$^{Dual}$ | 28270 | 28271 | -1 | pAzF | Tet3.0 | - | |
| | | sfGFP$^{Dual}$† | 28244 | 28244 | 0 | pAmF | Tet3.0 | pAzF Reduction | |
| | | sfGFP$^{Dual}$* | 28139 | 28138 | +1 | pAzF | Tet3.0 | -Met | |
| S2B | sfGFP$^{WT}$ | sfGFP$^{WT}$ | 27828 | - | - | Asp | Asn | - | |
| | | sfGFP$^{WT}$* | 27696 | 27696 | 0 | Asp | Asn | -Met | |
| | sfGFP$^{TAG}$+pEVOL-pAzF$^{TAG}$+pAzF | sfGFP$^{pAzF}$ | 27902 | 27902 | 0 | Asp | pAzF | - | |
| | | sfGFP$^{pAzF}$* | 27770 | 27770 | 0 | Asp | pAzF | -Met | |
| | sfGFP$^{TAG}$ + pEVOL-pAzF$^{TAG}$ + Tet3.0 | sfGFP$^{Phe}$ | 27861 | 27861 | 0 | Asp | Phe | - | |
| | | sfGFP$^{Asn}$ | 27827 | 27828 | -1 | Asp | Asn? | - | Near-Cognate Suppression |
| | | sfGFP$^{Asn}$* | 27696 | 27696 | 0 | Asp | Asn? | -Met | Near-Cognate Suppression |
| | sfGFP$^{TAG}$ + pEVOL-pAzF$^{TAG}$ -ncAA | sfGFP$^{Phe}$ | 27861 | 27861 | 0 | Asp | Phe | - | |
| | | sfGFP$^{Asn}$ | 27827 | 27828 | -1 | Asp | Asn? | - | Near-Cognate Suppression |
| | | sfGFP$^{Asn}$* | 27696 | 27696 | 0 | Asp | Asn? | -Met | Near-Cognate Suppression |
| S2D | sfGFP$^{WT}$ | sfGFP$^{WT}$ | 27828 | - | - | Asp | Asn | - | |
| | | sfGFP$^{WT}$* | 27696 | 27696 | 0 | Asp | Asn | -Met | |
| | sfGFP$^{TAA}$+pUltral-Tet3.0$^{TAA}$+Tet3.0 | sfGFP$^{Tet3.0}$ | 28197 | 28197 | 0 | Asp | Tet3.0 | - | |
| | | sfGFP$^{Tet3.0}$* | 28066 | 28065 | +1 | Asp | Tet3.0 | -Met | |
| | sfGFP$^{TAG}$+pUltral-Tet3.0$^{TAA}$+Tet3.0 | sfGFP$^{Tet3.0}$ | 27955 | 27955 | 0 | Asp | Tet3.0 | - | Near-Cognate Suppression |
| | | sfGFP$^{Tet3.0}$* | 27825 | 27823 | +2 | Asp | Tet3.0 | - | Near-Cognate Suppression |
| | sfGFP$^{TAG}$ +pEVOL-pAzF$^{TAG}$ +p-UltraTet3.0$^{TAA}$ +pAzF/Tet3.0 | sfGFP$^{pAzF}$ | 27902 | 27902 | 0 | Asp | pAzF | - | |
| | | sfGFP$^{pAzF}$* | 27770 | 27770 | 0 | Asp | pAzF | -Met | |
| S5A | sfGFP$^{WT}$ | sfGFP$^{WT}$ | 27828 | - | - | Asp | Asn | - | |
| | | sfGFP$^{WT}$* | 27696 | 27697 | -1 | Asp | Asn | -Met | |
| | sfGFP$^{pAzF}$ | sfGFP$^{pAzF}$ | 27902 | 27902 | 0 | Asp | pAzF | - | |
| | | sfGFP$^{pAzF}$† | 27873 | 27876 | -3 | Asp | pAmF | pAzF Reduction | |
| | | sfGFP$^{pAzF}$* | 27771 | 27770 | +1 | Asp | pAzF | -Met | |
| | sfGFP$^{pAzF}$ + DBCO-$NH_2$ | sfGFP$^{pAzF}$-DBCO-$NH_2$ | 28178 | 28178 | 0 | Asp | pAzF-DBCO-$NH_2$ | SPAAC Conjugation | |
| | | sfGFP$^{pAzF}$-DBCO-$NH_2$* | 28047 | 28046 | +1 | Asp | pAzF-DBCO-$NH_2$ | SPAAC Conjugation, -Met | |
| | | sfGFP$^{pAzF}$† | 27874 | 27876 | -2 | Asp | pAmF | pAzF Reduction | |
| S5B | sfGFP$^{WT}$ | sfGFP$^{WT}$ | 27828 | - | - | Asp | Asn | - | |
| | | sfGFP$^{WT}$* | 27696 | 27697 | -1 | Asp | Asn | -Met | |
| | sfGFP$^{Tet3.0}$ | sfGFP$^{Tet3.0}$ | 28197 | 28197 | 0 | Asp | Tet3.0 | - | |
| | | sfGFP$^{Tet3.0}$* | 28066 | 28065 | +1 | Asp | Tet3.0 | -Met | |
| | sfGFP$^{Tet3.0}$ + sTCO-OH | sfGFP$^{Tet3.0}$-sTCO-OH | 28321 | 28321 | 0 | Asp | Tet3.0-sTCO-OH | IEDDA Conjugation | |
| | | sfGFP$^{Tet3.0}$-sTCO-OH* | 28190 | 28189 | +1 | Asp | Tet3.0-sTCO-OH | IEDDA Conjugation, -Met | |
| S5C | sfGFP$^{WT}$ | sfGFP$^{WT}$ | 27828 | - | - | Asp | Asn | - | |
| | | sfGFP$^{WT}$* | 27696 | 27697 | -1 | Asp | Asn | -Met | |
| | sfGFP$^{Dual}$ | sfGFP$^{Dual}$ | 28270 | 28271 | -1 | pAzF | Tet3.0 | - | |
| | | sfGFP$^{Dual}$† | 28244 | 28244 | 0 | pAmF | Tet3.0 | pAzF Reduction | |
| | | sfGFP$^{Dual}$* | 28138 | 28138 | 0 | pAzF | Tet3.0 | -Met | |
| | sfGFP$^{Dual}$ + DBCO-$NH_2$ | sfGFP$^{Dual}$-DBCO-$NH_2$ | 28546 | 28547 | -1 | pAzF-DBCO-$NH_2$ | Tet3.0 | SPAAC Conjugation | |
| | | sfGFP$^{Dual}$-DBCO-$NH_2$* | 28415 | 28415 | 0 | pAzF-DBCO-$NH_2$ | Tet3.0 | SPAAC Conjugation, -Met | |
| | | sfGFP$^{Dual}$ | 28268 | 28271 | -3 | pAzF | Tet3.0 | - | Incomplete SPAAC Reaction |
| | | sfGFP$^{Dual}$† | 28244 | 28244 | 0 | pAmF | Tet3.0 | pAzF Reduction | |
| | | sfGFP$^{Dual}$†* | 28112 | 28113 | -1 | pAmF | Tet3.0 | pAzF Reduction -Met | |
| S5D | sfGFP$^{WT}$ | sfGFP$^{WT}$ | 27828 | - | - | Asp | Asn | - | |
| | | sfGFP$^{WT}$* | 27696 | 27697 | -1 | Asp | Asn | -Met | |
| | sfGFP$^{Dual}$ | sfGFP$^{Dual}$ | 28270 | 28271 | -1 | pAzF | Tet3.0 | - | |
| | | sfGFP$^{Dual}$† | 28244 | 28244 | 0 | pAmF | Tet3.0 | pAzF Reduction | |
| | | sfGFP$^{Dual}$* | 28138 | 28138 | 0 | pAzF | Tet3.0 | -Met | |
| | sfGFP$^{Dual}$ + sTCO | sfGFP$^{Dual}$-sTCO-OH | 28394 | 28395 | -1 | pAzF | Tet3.0-sTCO-OH | IEDDA Conjugation | |
| | | sfGFP$^{Dual}$-sTCO-OH† | 28368 | 28369 | -1 | pAmF | Tet3.0-sTCO-OH | IEDDA Conjugation, pAzF Reduction | |
| | | sfGFP$^{Dual}$-sTCO-OH* | 28263 | 28263 | 0 | pAzF | Tet3.0-sTCO-OH | IEDDA Conjugation, -Met | |
| S5E | sfGFP$^{WT}$ | sfGFP$^{WT}$ | 27828 | - | - | Asp | Asn | - | |
| | | sfGFP$^{WT}$* | 27696 | 27697 | -1 | Asp | Asn | -Met | |
| | sfGFP$^{Dual}$ | sfGFP$^{Dual}$ | 28270 | 28271 | -1 | pAzF | Tet3.0 | - | |
| | | sfGFP$^{Dual}$† | 28244 | 28244 | 0 | pAmF | Tet3.0 | pAzF Reduction | |
| | | sfGFP$^{Dual}$* | 28138 | 28138 | 0 | pAzF | Tet3.0 | -Met | |
| | sfGFP$^{Dual}$ +DBCO-$NH_2$ +sTCO-OH | sfGFP$^{Dual}$-DBCO-$NH_2$/sTCO-OH | 28670 | 28671 | -1 | pAzF-DBCO-$NH_2$ | Tet3.0-sTCO-OH | SPAAC Conjugation, IEDDA Conjugation | |
| | | sfGFP$^{Dual}$-DBCO-$NH_2$/sTCO-OH* | 28540 | 28539 | +1 | pAzF-DBCO-$NH_2$ | Tet3.0-sTCO-OH | SPAAC Conjugation, IEDDA Conjugation, -Met | |
| | | sfGFP$^{Dual}$-sTCO-OH | 28393 | 28395 | -2 | pAzF | Tet3.0-sTCO-OH | IEDDA Conjugation | Incomplete SPAAC Reaction |
| | | sfGFP$^{Dual}$-sTCO-OH† | 28368 | 28369 | -1 | pAmF | Tet3.0-sTCO-OH | IEDDA Conjugation, pAzF Reduction | Incomplete SPAAC Reaction |
| | | sfGFP$^{Dual}$-sTCO-OH†* | 28238 | 28237 | +1 | pAmF | Tet3.0-sTCO-OH | IEDDA Conjugation, pAzF Reduction, -Met | Incomplete SPAAC Reaction |

***Table S6***. Index of all indicated peaks present in all mass spectra presented in this study. Note that the mass measurement error is ±1 Da. All predicted masses were calculated based on the observed WT mass.
